# Development of the Early Childhood Duodenum across Ancestry, Geography and Environment

**DOI:** 10.64898/2025.12.15.694427

**Authors:** Joshua de Sousa Casal, Krishnan Raghunathan, Chelsea Asare, Abigail Plone, Nazanin Moradinasab, Junaid Iqbal, Lianna F. Wood, Elsy M. Ngwa, Xia Chen, S. Fisher Rhoads, Clara Baek, Dur-e Shahwar, Neha S. Dhaliwal, Madison Wong, Max Garrity-Janger, Lily P. Gillette, Stephanie Regis, Fatima Zulqarnain, Asra Usmani, Jason D. Boisvert, Casey R. Johnson, Jackson Larlee, Michael D. Anderson, Daniel Zeve, Elisa Saint-Denis, Thomas G. Wichman, Jeffrey La, Ashish Jain, Liang Sun, Lauren Scudari, Natalie N. Bhesania, Zehra Jamil, Michelle Galeas-Pena, Adam R. Greene, Aneeta Hotwani, Fedaa Najdawi, Shyam S. Raghavan, Donald E. Brown, Christopher A. Moskaluk, Heather H. Burris, Piotr Sliz, Phyllis R. Bishop, Scott B. Snapper, Kamran Sadiq, Sarah C. Glover, Muhammad Imran Nisar, Sana Syed, Jocelyn A. Silvester, Jose Ordovas-Montanes, Jay R. Thiagarajah

## Abstract

During early childhood, the proximal small intestinal mucosa plays a central role in growth, metabolism, immune priming, and neuronal development. Yet the cellular architecture and environmental responsiveness of the human small intestinal mucosa during this period remain poorly defined. Here, we generate a comprehensive cellular and spatial map of the duodenum from 87 children aged 6 months to 13 years, representing diverse ancestries and geographic contexts. This atlas integrates single-cell transcriptomic and spatial profiling with data on diet, social drivers of health, and environmental exposures. Using these data, we define mucosal cellular composition and chart its developmental trajectory in early childhood. Comparative analyses of children residing in the United States (US) and Pakistan reveal a differentiated enterocyte subset expressing the aquaglyceroporin, *AQP10* (AQP10^+^ enterocyte), that is enriched in children from the US. We show that emergence of this enterocyte state depends on lipid exposure to intestinal stem cells and correlates with dietary fat intake. We also identify a previously-undescribed thyrotropin-releasing hormone (TRH^+^) enteroendocrine cell and provide evidence for a local endocrine–epithelial–lymphocyte circuit. Our work establishes a detailed framework for pediatric duodenal mucosal development and illuminates how intestinal cellular dynamics are shaped by age and environment.

## Main

Early childhood is a critical period of human development, marked by rapid anthropometric growth and maturation of multiple organ systems. Meeting energetic demands during this phase depends on the efficient assimilation of nutrients and fluids through the gastrointestinal (GI) tract. Within the small intestine, the duodenum plays a uniquely central role: it is the primary site where acidic gastric contents are neutralized and where micronutrients, macronutrients, bile acids and enzymes converge to regulate nutrient absorption, host defense, and hormone production. Diet, genetics, and environmental exposures shape duodenal mucosal function, influencing metabolic programming, neurodevelopment, and physical growth trajectories. Understanding how the mucosal surface of the proximal small intestine changes at a cellular level during childhood is therefore essential for elucidating mechanisms of both healthy development and disease susceptibility across the lifespan.

Although significant progress has been made in mapping the human GI tract at single-cell resolution, these efforts have focused predominantly on the adult distal intestine, including the ileum and the colon^1–14^. Prior analyses have defined major epithelial, stromal, and immune populations in these tissues and provided insight into disease-associated states^15–27^. With the expanding scale and resolution of cellular atlases, we have begun to gain deeper insight into rare and previously understudied cell types, including nutrient-sensing enteroendocrine cells defined by their distinctive hormone-expression programs, as well as the recently described BEST4⁺ and INFLARE populations.^10,20,27–30^ Mapping multiple regions of the intestine has revealed that absorptive enterocytes constitute a highly heterogeneous population, varying both along the length of the gut and across individual villi.^6,13,31^ These cells deploy distinct repertoires of metabolite transporters that underpin regional specialization in nutrient uptake. Single-cell–level resolution now provides deeper insight into how this expansive surface senses and selectively absorbs nutrients in a finely zonated manner while simultaneously maintaining barrier defense against pathogens.

Recent efforts have resulted in the generation of a working community-guided atlas by the Human Cell Atlas Gut Bionetwork. However, these endeavors have highlighted a critical gap: the lack of early childhood representation.^32^ Developmentally focused single-cell studies have primarily characterized *in utero* intestinal formation, revealing the emergence of various major cell lineages^33–38^. However, *in utero* development represents a biological window driven by organogenesis rather than environmental adaptation. In contrast, early childhood represents a window of environmental exposures that facilitate rapid growth, immune education, and dynamic metabolic transitions. To date, this has not been systematically profiled at single-cell resolution in the duodenum. In addition, there has been limited inclusion of participants with diverse ancestry and environmental exposures across human single-cell transcriptomic datasets from all organs, reducing the generalizability and translational relevance of current resources^32,33^. A detailed, ancestrally and environmentally inclusive atlas of the pediatric duodenum is therefore needed to illuminate the cellular programs that shape growth, metabolism, and immunity during early childhood.

To address these gaps, we established the Gut-AGE project—an international, multi-omics initiative designed to map the childhood upper intestine at single-cell resolution across diverse ancestries, geographies, and environmental contexts. Here, we present an integrated atlas of the duodenum generated from 88 children of which 87 were aged 6 months to 13 years residing in the United States (US) and Pakistan. This dataset captures transcriptional, compositional, and spatial features of the duodenal mucosa, complemented by detailed information on tissue morphology, diet, social drivers of health, and environmental exposures (available at www.gut-age.com, cellxgene.cziscience.com).

Our analyses reveal epithelial, stromal, and immune cell types and cell transcriptional states that are conserved across all participants, establishing a robust reference framework for interpreting healthy variation across populations and for contextualizing genomic and transcriptomic studies of pediatric disease. Analysis of age-dependent changes uncovered major epithelial and immune transitions. We established that these mucosal cell shifts occur predominantly within the first four years of life and stabilize thereafter. The breadth of our cohort allowed detailed interrogation of rare cell types and variations in cellular states. Leveraging this resolution, we identified a diet-adapted absorptive AQP10^+^ late enterocyte state and uncovered an intrinsic mucosal hormonal circuit centered on a previously unrecognized TRH^+^ enteroendocrine cell subtype.

### Cohort Enrollment, Structure, and Metadata

To map duodenal mucosa within representative populations, we prospectively enrolled children undergoing diagnostic esophagogastroduodenoscopy (EGD) for indications with a high likelihood of being non-diagnostic for active mucosal disease (e.g., foreign body retrieval, non-specific abdominal pain, and reflux). See **Extended table 1, 2, Supplementary table 1** and **Methods** for full indications and inclusion/exclusion criteria. To capture variability in human intestinal composition across geographic environments, individuals were enrolled across three sites in the US (Boston, Massachusetts; Jackson, Mississippi; and Charlottesville, Virginia), and one site in Pakistan (Karachi, Sindh) **(Figure 1A)**. Metadata and biospecimens were collected. Blood was used for PBMC isolation and whole genome sequencing. Duodenal biopsies were processed for single-cell RNA-sequencing (scRNA-seq), bulk RNA-sequencing, histology, and multiplexed immunofluorescence (MxIF) imaging **(Figure 1A)**.

**Figure 1:**
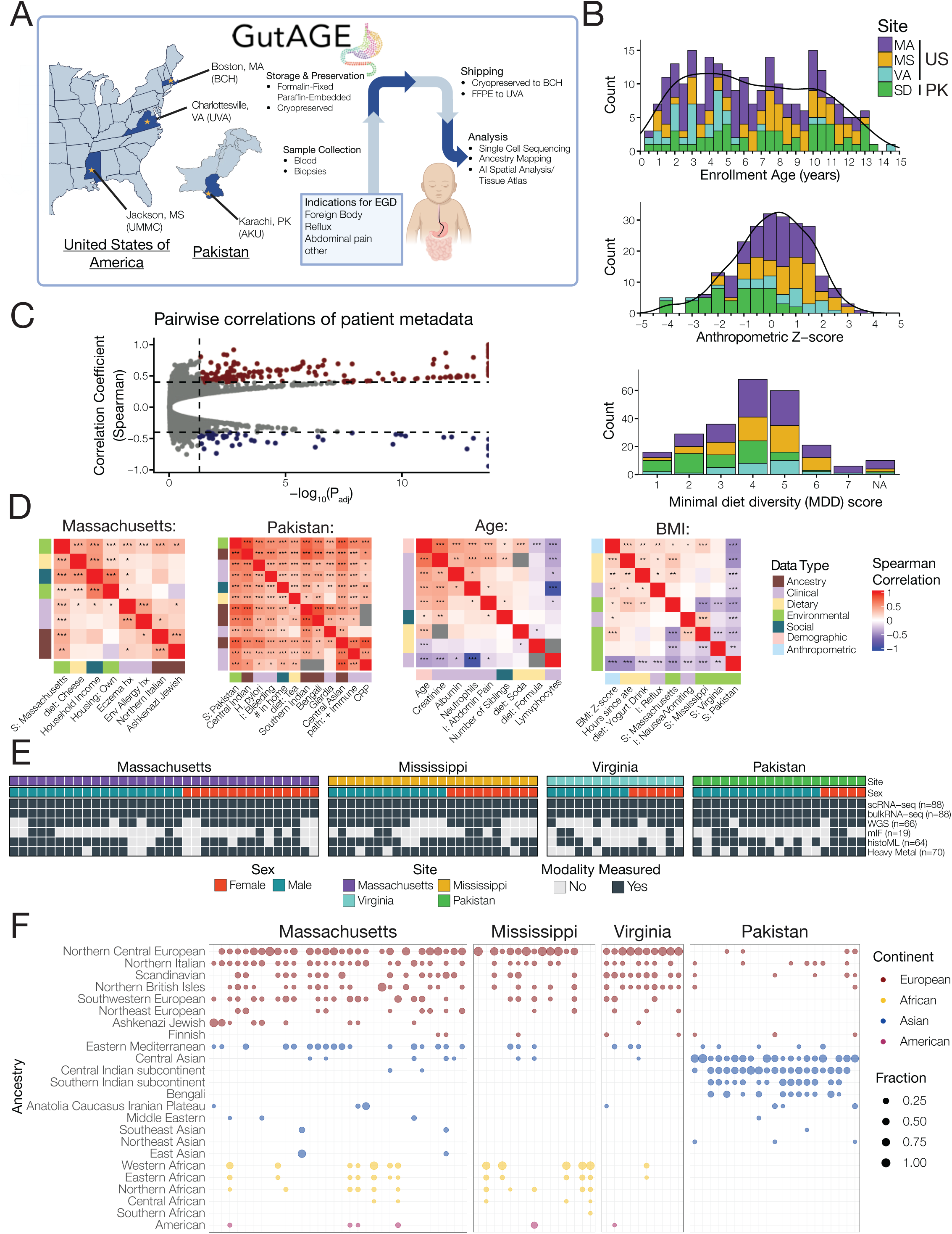
Gut-AGE captures representative variation in the pediatric duodenum. **A.** Schematic overview of patient recruitment and sample collection across three sites in the United States and one site in Pakistan. EGD, esophagogastroduodenoscopy. FFPE, formalin-fixed paraffin-embedded. **B.** Distribution of enrollment age (top), anthropometry z-scores (WHO growth curves; Weight/length under 2 years old, BMI over 2 years old, middle), and minimum diet diversity (MDD) score (bottom) across the Gut-AGE cohort. **C.** Volcano plot of pairwise correlations across all metadata variables surveyed. Color represents significant correlations (red, correlation coefficient > 0.5; blue, correlation coefficient < −0.5). **D.** Correlation matrices of highly associated metadata modules, colored by data category and Spearman correlation coefficient. S, site; hx, history; I, endoscopy indication; path: + immune, pathology description of elevated immune cells in duodenal biopsy. **E.** Representation of complementary data modalities in participants with both single-cell and bulk RNA-seq datasets. n_sc_ denotes the number of samples shared between the specified modality and scRNA-seq. n_T_ denotes the total number of participants evaluated using the specified modality. scRNA-seq, single-cell RNA-sequencing; WGS, whole genome sequencing; MxIF, multiplexed immunofluorescence; histoML, machine learning of hematoxylin and eosin stains. **F.** Dot plot of genetic ancestry deconvolution of participants within the Gut-AGE study, color represents continent of origin and dot size depicts fraction of genetic ancestry.

Enrolled children ranged from 0.5-13 years of age with a median age of 6 **(Figure 1B)**. Anthropometric (weight/length if age < 2 years, BMI if age >2 years) z-scores were normally distributed with a median of 0.14 (95% CI= −0.02 – 0.42, SD = 1.49) **(Figure 1B),** consistent with the anthropometric distribution of a healthy population. Notably, these z-scores were lower in the Pakistan cohort, representative of the pediatric community population from which participants were recruited^39^. Surveys were used to collect detailed clinical, environmental, and socioeconomic data. Survey items included medical and medication history, feeding and dietary data, and social drivers of health, such as perceived stress and food security (Hunger Vital Sign^TM^)^40,41^. Given the importance of diet on mucosal function we modified the World Health Organization, Infant and Young Child Feeding Minimum Dietary Diversity (MDD) indicator to assess dietary diversity in non-breastfeeding children who had fasted for endoscopy. Specifically, we assessed the number of different food groups consumed over a ‘typical’ 24-hour recall period and used a score of 4 as the threshold for adequate dietary diversity (**Supplementary table 1**). Using this approach, the majority of US participants had adequate dietary diversity, (median: 4, SD: ±1.4), whereas participants in Pakistan tended to be below this cutoff (median: 3, SD:±1.3) (**Figure 1B)**.

Given the breadth of the metadata obtained, we conducted pairwise Spearman’s rank correlation analyses to assess trends in the cohort data structure (**Figure 1C**, **Extended data 1A**). We found several clusters of associations related to site, age and BMI (**Figure 1D**). Notable associations included participants enrolled in Massachusetts and a history of eczema, and participants from Pakistan and *Giardia* colonization. To assess duodenal mucosa composition in a tissue context, we generated multiple types of data across the cohort to complement scRNA-seq (**Figure 1E**), including spatial data through multiplexed protein analysis (MxIF) and machine-learning based histological analysis (histo-ML). We also recorded individual-level ancestry based on supervised admixture genotype analysis of whole genome sequencing (WGS). An overview of continent and region-level data (**Figure 1F**) outlines the varied and wide distribution of genetic ancestry across our cohort - an important criterion for establishing a generalized human reference atlas.

### An Atlas of Duodenal Cellular Diversity in Early Childhood

We report duodenal scRNA-seq data of 226,706 cells from 88 participants after removal of doublets and poor-quality cells (quality control cutoffs summarized in **Supplementary data 1 and Supplementary table 2)**. To reduce costs and better normalize technical batch effects, multiple participants were pooled and then later demultiplexed based on unique SNPs within each individuals transcriptome^42,43^. Following genetic demultiplexing and quality control filtering, we isolated an average of 2580 cells per participant, with a median interparticipant doublet rate of 5.92% (SD:±5.25), all of which were removed for downstream analysis **(Supplementary data 1)**. When selecting participants to pool, we ensured that biological variables (such as age and site) were orthogonal to technical variables (such as participant pool and sequencing batch). Consequently, although small percentage of the dataset variance is associated with technical variables, their association with biological variables have been reduced allowing for more effective dataset integration **(Supplementary data 2)**.

Top level cell clustering identified 8 major cell types, which were annotated as epithelial cells, T cells, B cells, plasma cells, myeloid cells, granulocytes, endothelial cells, and stromal cells **(Figure 2A)**. Leveraging our large cell number and diversity, we used iterative clustering to subdivide and identify the cell subtypes and states within clusters. Overall, we identified 128 end cell states, which were evaluated and annotated by experts within the field and compared to previously published annotations.^1–3,5,6,8^ For each cell subtype and state, we provide a comprehensive glossary of marker genes and our associated rationale **(Supplementary: cell glossary).** In addition, a full list of differentially expressed marker genes are provided in **Supplementary table 3-6.** To understand cell state distance and the extent of their similarity, we performed tiered clustering within a taxonomical tree **(Figure 2B)**. As expected, end cell states primarily cluster with other states that originated from the same major cell type. One interesting exception are mucosal mast cells, which exhibit greater transcriptional similarity to epithelial tuft cells than to other immune cells. Most identified cell states were well represented across our cohort **(Extended data 2A)** and highly conserved across age, sex, and geographic site **(Extended data 2B-C)**.

**Figure 2:**
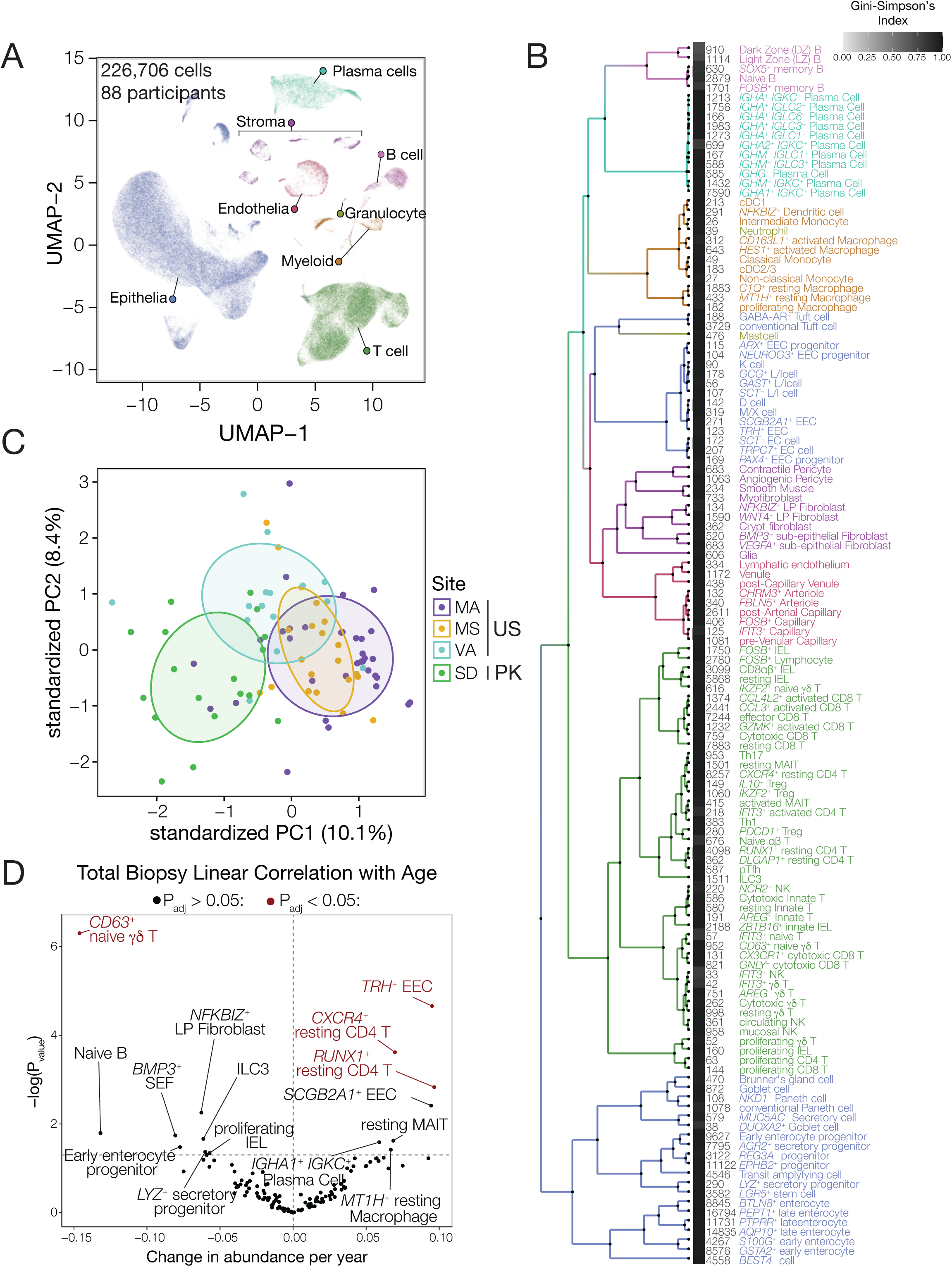
Cellular composition within the duodenum across age and geography. **A.** Uniform manifold approximation and projection (UMAP) display of 226,706 cells across duodenal biopsies from 88 participants, colored by broad cell identity. **B.** Taxonomic tree of Gut-AGE end cell states constructed using the ARBOL package, including squares depicting cohort representation (Gini-Simpson’s index) and the number of cells in the dataset for each cell state. Labels and branches are colored by broad cell identity. **C.** Principal component analysis of the cell composition for each individual participant. Each point represents a single participant, and distance represents differences in biopsy composition. Points are colored by site and ellipses are representative of site centroids at a confidence interval of 50%. **D.** Volcano plot depicting linear associations between cellular abundance and age in which each point represents a cell state. A linear model was fit for the abundance of each cell state across age, slope was plotted on the X axis, and the corresponding unadjusted p value was plotted on the Y axis. Significant points (p < 0.05) were labelled, and color represents points that remained significant after p value correction (FDR method). LP, lamina propria; SEF, sub-epithelial fibroblast; EEC, enteroendocrine cell; ILC, innate lymphoid cell; MAIT, mucosal-associated invariant T cell; IEL, intra-epithelial lymphocyte.

Among significant associations with participant metadata, geographic site was the largest driver of variance in cell state abundance. Specifically, we find an increase in representation of immune cell states, such as an undescribed DLGAP^+^ CD4 T cell state and resting IELs, in the Pakistan cohort. Genes within the DLGAP family encode scaffold proteins that localize to neuronal dendrites and have been implicated neuronal synaptic formation and glutamate signaling, but their exact role in T cells warrants further investigation^44,45^. Additional marker genes for these populations are summarized in our **cell glossary,** as well as in **Supplementary table 6**. Pakistan participants also had elevated proportions of REG3A^+^ progenitor cells, which have transcriptional similarities to a cell state implicated in damage and regenerative responses **(Extended data 2D)**.^46,47^ We identified eight cell states that were present in a small proportion of participants (<25%), including germinal center B cells, inflammatory monocytes, neutrophils and cell states with strong interferon signatures. The presence of these cell types and states may indicate an ongoing immune response in these participants.

To quantify interindividual differences in cell composition in an unbiased fashion, we calculated the proportional representation of each bottom-level cell state in each biopsy and performed principal component analysis (PCA) of sample composition **(Figure 2C; Methods)**. While there was considerable overlap in the duodenal cellular composition of biopsies from US sites (US-MA, US-MS, and US-VA), biopsies from the Pakistan site (PK-SD) exhibited a distinct cellular composition relative to the US cohort that primarily separated in PC1. PC1 variance was driven in the negative direction (more associated with PK-SD participants) by venules and multiple immune cell populations, including IELs, B cells and T follicular helper cells (Tfhs). In contrast, epithelial cell and tissue-resident macrophage states drove PC1 in the positive direction **(Extended data 2D)**. These findings indicate that aspects of childhood duodenal cellular composition are associated with participant country of residence, potentially attributable to differences in ancestry and cumulative environmental exposures.

We observed that age was not associated with large differences in overall biopsy composition or diversity, as measured using the Shannon or Gini-Simpson index **(Extended data 2E)**. Thus, overall duodenal cellular composition appears to be relatively conserved between 6 months and 13 years of age. To identify individual cell states that vary with age, we fitted linear regression models with cell state abundances as dependent variables. We found multiple naïve lymphocyte cell states were negatively associated with age, including a CD63^+^ naïve ψ8 T cells **(Figure 2D, Extended data 2F)**. In contrast, CD4^+^ T cell states, IgA^+^ Plasma cells, and mucosal-associated invariant T-cells (MAIT) cells were more abundant among older individuals. However, small variations in overall cell state representation due to biopsy sampling may obscure differences in cell state composition when analyzed as a function of all cells in the biopsy. Therefore, to better account for any bias related to sampling of other cell types (e.g., epithelial cells), we analyzed changes in lymphocyte cell states as a proportion of total T/NK/ILCs. This revealed that ILC3s, naive αβ T cells and mucosal NK cells are also negatively associated with age in addition to the trends observed in the previous analysis of all cell states **(Extended data 2G, H)**. We infer that these changes in the first four years of life likely represent a broad shift from embryonically seeded naïve and innate-like lymphocyte populations towards tissue resident adaptive populations as individuals encounter environmental antigen through microbial exposures and diet.

### Spatial Architecture of the Duodenum defined by Multiplexed Tissue Imaging and Morphological Feature Mapping

To understand how duodenal cells are spatially related within the context of both age and site we conducted multiplex immunofluorescence (MxIF) analysis of duodenal tissues from our cohort. Tissue microarrays (TMA) containing multiple intestinal biopsies were generated from formalin-fixed paraffin-embedded blocks. All TMAs were stained and imaged using a custom 30-plex panel using our recently developed SPECTRE-Plex method^48^. Raw fluorescence images underwent initial processing involving generation of an optimal tissue focus map, exposure bracketing, alignment, stitching and background correction (**Figure 3A and Supplementary data 3,4,5)**. Images were segmented to identify cells based on a three-stain trained deep-learning algorithm (Cellpose3).^49^ Segmented cells were annotated using a hierarchical classification method (**Supplementary data 6, 7**). Following annotation, custom analytical pipelines were used to derive tissue cell proportions, intestinal brush border measurements, areas of cell enrichment and cell neighborhood analyses. **Figure 3B** shows a series of representative multiplexed tissues, highlighting identification of different epithelial, immune and stromal cell types, functional markers such as the intestinal digestive enzyme lactase and markers of cell proliferation state. Cell-specific antigens allowed coordinate mapping by cell-type (**Figure 3C**) and spatial cell proportion analysis (**Extended data 3A, 3B**).

**Figure 3:**
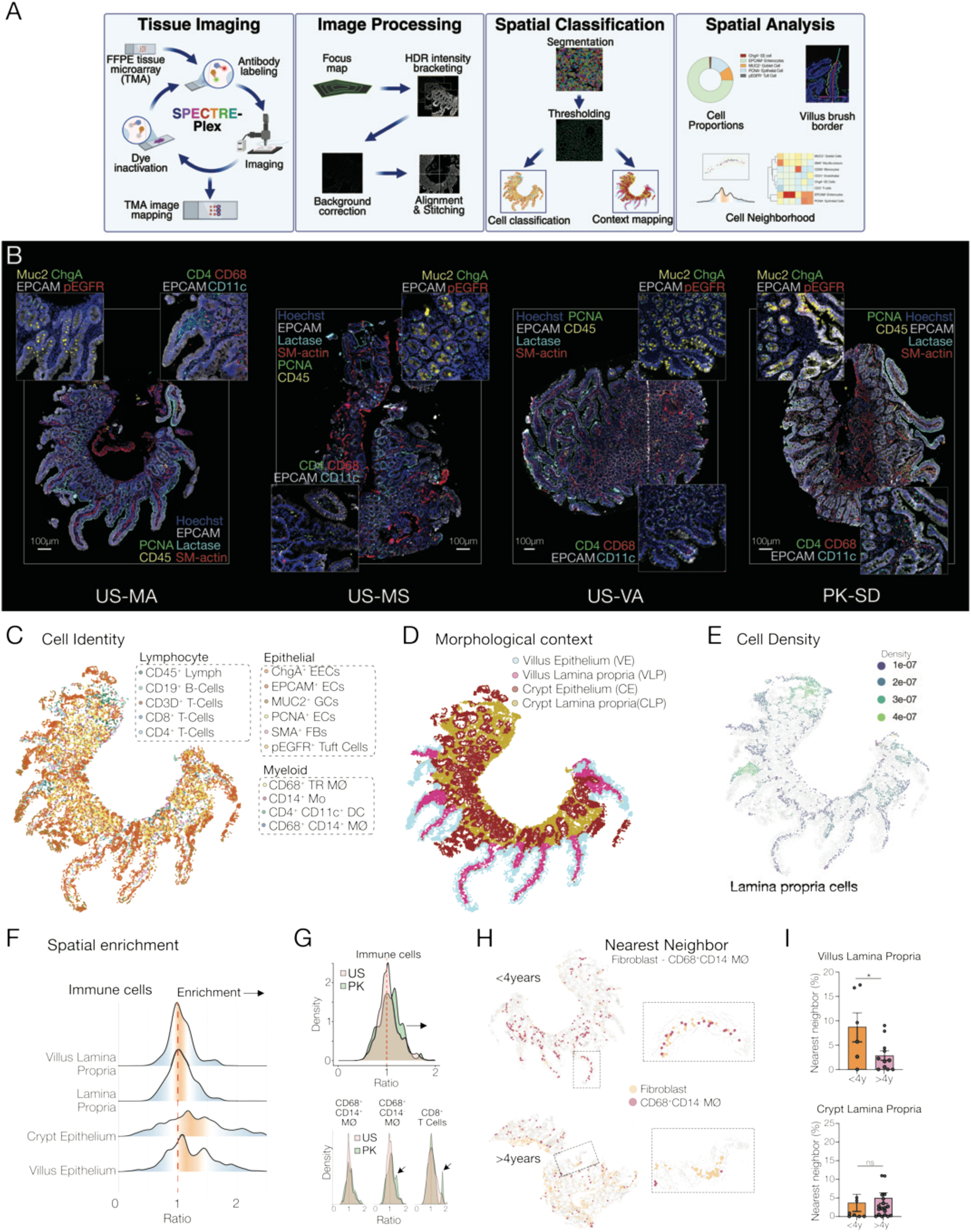
Multiplex-Immunofluorescence reveals the cellular geography of duodenal development. **A.** Schematic of the Multiplex Immunofluorescence (MxIF) pipeline. HDR-High Dynamic Range **B.** Representative MxIF images and corresponding insets showing the spatial distribution of select proteins in the duodenal tissue from each of the four sites. (3.0 year-old male from US-MA, 8.25 year-old female from US-MS, 5.75 year-old male from US-VA, and 0.5 year-old female from PK-SD). Each main panel shows the tissues stained with Hoechst (blue), SM-actin (red), Lactase (cyan), EPCAM (grey), PCNA (green), and CD45 (yellow). One inset shows the staining for Hoechst (blue), Muc2 (yellow), ChgA (green), EPCAM (grey), and pEGFR (red). The other inset shows the staining for Hoechst (blue), CD4 (green), CD68 (red), CD11c (cyan), and EPCAM (grey). **C.** Representative pointillistic image showing spatial distribution of all cell types in a biopsy from the United States (3.0 year-old male from US-MA). **D.** Representative context map showing the four different morphological regions of the intestine. (3.0 year-old male from US-MA). **E.** Representative immune cell density plot showing the relative regional enrichment of immune cells in a biopsy from the United States (3.0 year-old male from US-MA). **F.** Cell density plots of relative spatial enrichment of immune cells in the different tissue regions of the duodenum. Ratio greater than 1 implies the immune cells are more spatially clustered together while ratio less than 1 implies immune cells are spatially distanced. **G.** (Top) Cell density plots of spatial enrichment of all immune cells in the villus lamina propria of duodenum grouped by country (Pakistan and the US). (Bottom) Cell density plots comparing spatial enrichment of CD68^+^CD14^+^ and CD68^+^CD14^-^ macrophage and CD8^+^ T Cell populations grouped by participant country (Pakistan and the US). **H.** (Top) Representative pointillistic map showing the spatial relationship of fibroblasts and CD68^+^CD14^-^ macrophages in a biopsy from a child less than 4 years old (1 year old male from US-MA) and (Bottom) from a child over 4 years old (5.75 year-old female from US-MA). Insets show the spatial relationship within a single well-oriented villus region from these biopsies. **I.** Quantification of change in spatial proximity of CD68^+^ macrophages to fibroblast in the villus lamina propria in young children (age < 4 years) and in older children (age > 4years) in the villus lamina propria (top) and crypt lamina propria (bottom).

Complex tissues are composed of morphological and structural features unique to the specific organ and intrinsic to tissue function. In the duodenum, this is represented by crypt glands and villi as one example or the epithelial barrier and the underlying lamina propria as another. Most existing methods conduct spatial analysis agnostic to these contextual features. We recently developed a series of methods based on mathematical morphological operations that we extended to MxIF images to map small intestinal morphological features (**Figure 3D, Extended data 3C**)^50^. These feature maps can be used to provide additional contextual information to aid interpretation of spatial analyses such as cell density and relative enrichment and cell-cell neighborhoods (**Figures 3E-I**). Spatial enrichment analysis of tissue immune cells - a relational measure of cell density within an identified local environment - showed relative enrichment within both the villus and crypt epithelial compartments (**Figure 3F**). Analysis of immune cell enrichment within the villus lamina propria showed increased enrichment in tissues from Pakistan relative to those from the US. This enrichment was largely driven by two cell sub-types, CD68^+^CD14^-^ macrophages and CD8^+^ T cells (**Figure 3G**). Given the early age-dependent variation in immune populations observed during the first four years in our overall scRNA-seq analyses and the known immune regulatory role of tissue stromal fibroblasts, we conducted tissue nearest-neighbor analysis dichotomized by age (> or <4 years) (**Figure 3H**)^51–53^. We found a significantly increased percentage of CD68^+^CD14^-^ macrophages in close proximity to tissue fibroblasts, specifically within the villus lamina propria in the youngest participants (**Figure 3I**). Importantly, we observed no significant change in relative populations of these cells between the two age groups (**Extended data 3D-E)**. These data are consistent with the scRNA-seq data which showed both fibroblast and macrophages were similarly abundant between the two age groups (**Extended data 3F-G)**.

### AI-guided Histology Analysis Identifies Site-Specific Duodenal Features

To investigate architectural variation between tissues in an unbiased fashion, we developed a self-supervised learning (SSL)-based AI model for extracting Whole Slide Image (WSI)-level features from H&E-stained duodenal biopsies collected across all our sites **(Extended data 4A)**. Over 45,000 non-overlapping patches, each 256×256 pixels, were extracted from the WSIs with exclusion of any patches containing less than 30% tissue **(Extended data 4B).** Patches were then used to fine-tune the pre-trained UNI, a large Vision Transformer model, to produce 1024-dimensional embeddings using the DINOv2 self-supervised learning approach (**Figure 4A**)^54–56^. Following the initial cluster centroid setting, a GMM-based approach was fitted to each WSI, with each mixture component representing a distinct histological feature (**Figure 4A**). The frequency of the eight histological features identified varied across sites (**Figure 4C** and **Extended data 4D)**. PCA was performed on the extracted histological features, of which the first two principal components were primarily associated with differences across geography **(Figure 4B)**. A Random Forest classifier was then trained using these features to predict whether a WSI is from Pakistan or the US. The classifier achieved a weighted F1 score of 0.94, with feature 5 and feature 6 being the most important for classification **(Extended data 4C)**.

**Figure 4:**
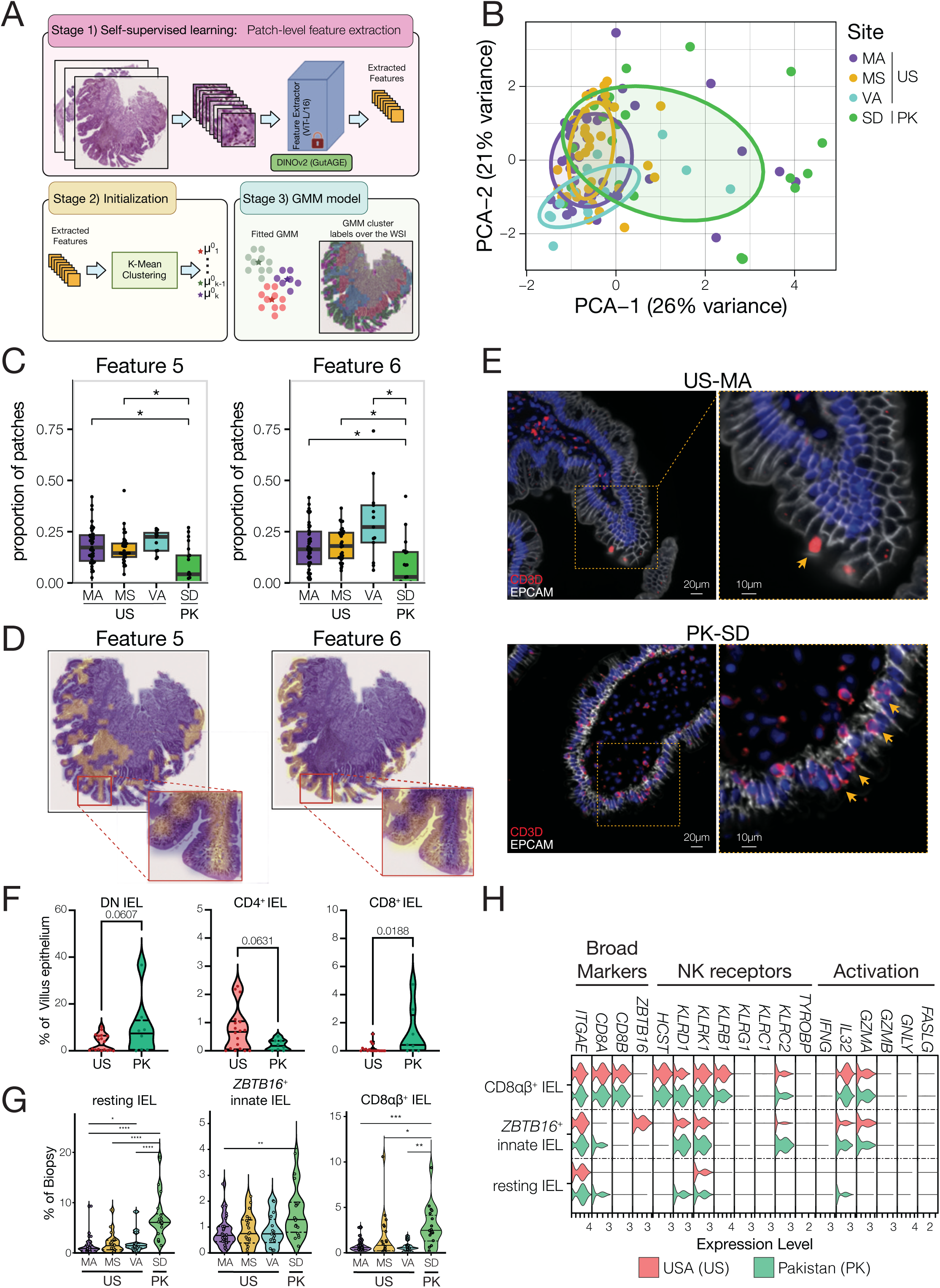
Unbiased machine-learning identifies differences in villus composition across geographic sites. **A.** Schematic overview of patch breakdown, feature extraction and feature identification by k-means clustering. **B.** Principal component analysis of patch-level features for each individual participant biopsy. Each point represents a single participant, and distance represents differences in the representation of identified patch features. Points are colored by site and ellipses are representative of site centroids at a confidence interval of 50%. **C.** Comparison of proportional patch assignment of Feature 5 and Feature 6 across enrollment sites. For each biopsy, feature proportion was calculated by dividing the number of patches assigned a specific feature by the total number of patches identified. **D.** Representative whole slide images of biopsies colored by regions associated with a specific feature. **E.** Representative MxIF images of a biopsy from the United States (Boston, Male, 35 months, top) and a biopsy from Pakistan (Male, 55 months, bottom) stained for EPCAM (white), CD3D (red) and DAPI. Yellow arrows highlight IELs and dotted yellow boxes denote enlarged insets (right). **F and G.** Violin plots of the relative proportion of each intraepithelial lymphocyte population (IEL) identified by MxIF **(F)** or scRNA-seq **(G)**. IEL abundance was divided by the total number of cells identified within the villus **(F)** or all cells within the biopsy **(G)**. DN, double negative. **H.** Violin plot depicting gene expression of IEL identity markers across IEL states. Color denotes country of enrollment. * p < 0.05, ** p < 0.01, ***p < 0.001, **** p < 0.0001.

The visualization of these features was overlaid on the WSIs and interpreted by three pathologists **(Figure 4B–D, Extended data 4E–F)** (Fleiss’ Kappa 0.83). We next evaluated whether there were significant differences in each of the specific WSI-level features across different geographic sites. We found that feature 5 and feature 6, which were annotated to mark the villus lamina propria and villus epithelium respectively, had significantly lower frequency in biopsies from Pakistan compared to any of the US sites (**Figure 4C, Extended data 4D**). Given these feature annotations, we assessed site associated changes in the composition of epithelial and immune cells within the villus by MxIF in tissue from the US and Pakistan. Overall, we found an increase in the number of intraepithelial lymphocytes (IELs) in biopsies from Pakistan and more specifically a significant increase in CD8^+^ IELs (**Figure 4E, 4F**). We did not observe a difference in the proportion of various epithelial populations between the two sites **(Extended data 4H)**. In agreement with our MxIF findings, we found that, within our scRNA-seq data, the proportions of both resting IELs and CD8αβ^+^ IELs were elevated in biopsies from Pakistan compared to biopsies from the US **(Figure 4G)**. In contrast, natural IELs (defined here by their expression of *CD8A* and *ZBTB16*) were similarly abundant across countries^57–59^. Prior investigations of human IEL function have demonstrated that pathogenic subsets are enriched in cytotoxic (*GZMA, GZMB, GNLY*) and inflammatory features, such as *KLRC2* and *IFNG*^60–65^. Evaluating the expression of these markers across IEL subsets and geographic sites, we found little evidence of an activated response, with no detectable expression of *IFNG*, *GZMB*, or *GNLY* **(Figure 4H)**. *GZMA* was expressed in multiple IEL populations, with similar expression in all CD8^+^ subsets and reduced relative to activated CD8 T cells **(Extended data 4I)**. *KLRC2*, which has been positively associated with IELs found in celiac disease, was strongly expressed in CD8^+^ IELs in all participants^63^. Therefore, site-level differences in IEL abundance likely reflect a normal, homeostatic adaptation to local antigen exposure rather than a pathogenic process.

### Distinct Absorptive Enterocyte Lineages Reveal Functional Specialization to Environment and Diet

The epithelium is critical for nutrient processing and assimilation in the small intestine. This is especially important during early childhood growth and development. We therefore investigated variability in epithelial composition within our scRNA-seq data in greater detail. In total, we identified 118,805 epithelial cells across 88 participants (**Figure 5A**). We describe 14 epithelial cell subtypes and 35 states, including progenitor cells, absorptive enterocytes, and multiple secretory cells, using iterative sub-clustering. These included subsets of BEST4⁺/CFTR⁺ cells, MUC6⁺ Brunner’s submucosal gland cells, and MUC5AC^+^ mucus cells, similar to the surface mucus cells of the stomach^8,10,27,66–68^. Enterocytes form a continuous distribution from multipotent cells to terminally differentiated states^69^. Within this continuum, we also observed that differentiated enterocytes display striking functional specialization, with distinct groups exhibiting enrichment for discrete metabolic, immunoregulatory, or transport pathways that extend well beyond canonical nutrient absorption roles. To organize and contextualize absorptive cell states, we performed pseudotime-mapped trajectory analysis initiated from the LGR5⁺ intestinal stem cell (ISC) cluster (**Figure 5B**)^70^. Based on this analysis, we identified three distinct but related cell states among highly differentiated late enterocytes. We named these enterocyte states by their most upregulated genes (*PEPT1*, *PTPRR*, and *AQP10*), with each representing a unique state within the absorptive lineage.

**Figure 5:**
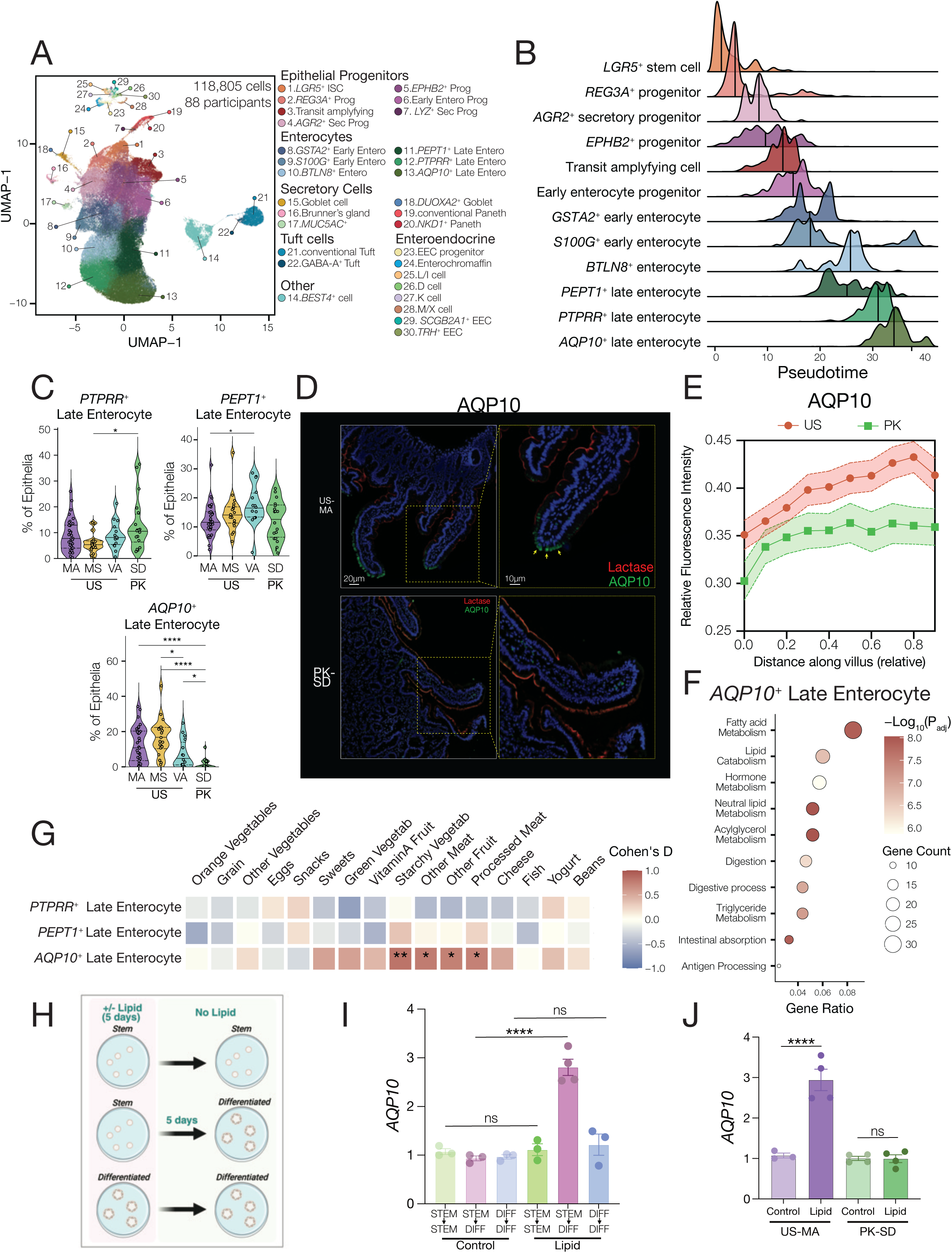
AQP10^+^ enterocytes are a lipid-responsive adaptation to Western diets. **A.** UMAP representation of 118,805 duodenal epithelial cells from 88 participants, colored by epithelial cell state. **B.** Ridge plot of pseudotime distribution for each progenitor and enterocyte state identified. Color corresponds to cell identity, and solid lines indicate the median value for each distribution. Pseudotime trajectory was calculated using Monocle3, originating from the LGR5^+^ intestinal stem cell cluster. **C.** Frequency of late enterocyte states as a proportion of epithelial cells by enrollment site. **D.** Representative MxIF images of biopsies from the US (Boston participant, Male, 35 months old) and Pakistan (Pakistan participant, female, 6 months old) stained with Lactase (red) and AQP10 (green). **E.** Quantification of AQP10 intensity along the intestinal villus by enrollment country. **F.** Top 10 GO terms enriched in genes significantly differentially expressed in AQP10^+^ late enterocytes relative to all other epithelial states. P values were corrected for FDR using the BH method. A full list of epithelial GO terms is available in **Supplementary table 7. G.** Associations between recent consumption of foods and late enterocyte state abundance. Color denotes effect size (Cohen’s D). Student’s t-test was used for statistical comparisons. **H.** Schematic representation of organoid experiment conditions. Organoids were cultured in the presence of a lipid cocktail composed of dietary lipids for 5 days. **I.** RNA expression of relative *AQP10* expression, split by the conditions outlined in **H**. **J.** Relative *AQP10* expression in differentiated organoids by site and lipid treatment. * p < 0.05, ** p < 0.01, ***p < 0.001, **** p < 0.0001.

Evaluating epithelial state across study sites, we note that most epithelial cell states are well represented and conserved across these distinct environments including intestinal stem cells, transit amplifying cells and goblet cells. However, we found that relative proportions of late absorptive enterocytes vary across environment (**Figure 5C**). Particularly striking was a significant country-level reduction in the relative proportion of AQP10^+^ enterocytes in participants recruited in Pakistan versus the US **(Figure 5C)**. Although AQP10^+^ enterocytes were the most abundant enterocyte state in US participants, they were rare in participants from Pakistan. MxIF staining of AQP10 verified our scRNA-seq findings and localized AQP10^+^ enterocytes to the villus tip (**Figure 5D**). Quantification of AQP10 localization within the epithelium showed that expression was lower, both overall and at villus tips, in children residing in Pakistan relative to children in the US (**Figure 5E**). Our findings identify AQP10 as a marker for a previously undescribed villus-tip enterocyte subset, unique to the proximal small intestine, that is differentially abundant across pediatric populations and geographies.^71,72^

To understand the potential physiological role of this AQP10^+^ cell state, we performed differential gene expression and GO term enrichment of each late enterocyte state relative to one another. AQP10^+^ enterocytes were highly enriched for gene pathways associated with lipid metabolism and absorption **(Figure 5F, Extended data 5A, B, C)**. Lipid digestion in the proximal intestine relies on enzymatic cleavage of dietary triglycerides within the lumen and subsequent epithelial absorption of the resultant cleavage products **(Extended data 5D)**^73^. The major products of luminally cleaved triglycerides have classically been assumed to be free fatty acids, and mono-acyl glycerols.^74^ Within the cytosol of enterocytes, these components are re-esterified via acyltransferases and packaged for systemic circulation^73^. In our epithelial cluster, only PTPRR⁺ and AQP10⁺ enterocytes express the full complement of genes required for lipid absorption and triglyceride re-esterification (**Extended data 5E**). In agreement with our observation regarding AQP10^+^ late enterocyte localization, prior work has demonstrated that lipid absorption within the small intestine occurs primarily within the upper third of villi.^75^ AQP10⁺ cells also express additional aquaporin family members, including the aquaglyceroporins AQP3 and AQP7 which transport glycerol as well as water across membranes^76,77^. Notably, AQP10 is present in humans, but not rodents, and exhibits pH-gated glycerol transport^76,78^. Emerging evidence suggests that free glycerol, either generated or delivered luminally, constitutes a small yet meaningful component of duodenal contents that fluctuates with diet and with the relative contributions of pancreatic and gastric lipases^79^. PTPRR⁺ and AQP10⁺ enterocytes therefore likely constitute the primary absorptive states responsible for lipid uptake in the duodenum, with AQP10⁺ late enterocytes further specialized to facilitate absorption of luminal free glycerol.

Compared to other late enterocytes, AQP10⁺ cells also showed increased expression of lipid-processing genes, including *APOA4, APOA1*, and *PRAP1*. In contrast, PEPT1⁺ enterocytes exhibited elevated expression of the sugar transporter *SLC2A2*, while PTPRR⁺ enterocytes showed increased expression of *IL15* and *BTNL8*, genes previously linked to support of tissue-resident T cells, specifically IELs (**Extended data 5F, G**)^80–82^. Therefore, PTPRR^+^ enterocytes may represent the major enterocyte responsible for IEL maintenance, especially in the Pakistan cohort. These patterns suggest a specialized role in carbohydrate absorption for PEPT1⁺ enterocytes and a role in immune regulation for PTPRR⁺ enterocytes. Overall, we identified multiple undescribed absorptive enterocytes in the duodenum, which possess distinct functional end states that contribute to nutrient acquisition and mucosal homeostasis, paralleling the diversity observed among secretory epithelial lineages.^69,83,84^ Notably, the relative abundance of these states varies across geographic sites, highlighting the natural physiological diversity of the epithelial surface.

The absorptive epithelium of the small intestine has been shown to remodel rapidly in response to changes in diet in mice^85,86^. Therefore, we hypothesized that since AQP10⁺ enterocytes express a glycerol channel alongside key lipid-absorption machinery, this state may be diet-responsive and specialized for luminal fat and glycerol uptake. To assess this, we related epithelial cell-state frequencies to dietary metadata (**Figure 5G**). Among absorptive late enterocytes, the abundance of AQP10⁺ cells was strongly associated with multiple dietary components, including processed and unprocessed meats and starchy vegetables. Meat intake is linked to higher levels of dietary fat, and processed foods often contain added glycerol, consistent with the possibility that AQP10⁺ cells expand in response to increased fat and free glycerol exposure in the U.S. diet^87–89^.

To directly interrogate lipid responsiveness, we generated participant-derived duodenal enteroids, exposed them to a physiologically relevant lipid mixture (**Methods**) or vehicle, and quantified *AQP10* expression (**Figure 5H**). U.S.-derived enteroids showed no change in *AQP10* expression when lipid treatment was applied to stem/progenitor–rich or fully differentiated cultures. However, when enteroids were exposed to lipids during the stem phase and then *subsequently* differentiated, *AQP10* expression increased significantly relative to controls (**Figure 5I**). In contrast, Pakistan-derived enteroids showed only minimal induction of *AQP10* under the same conditions (**Figure 5J**). These findings indicate that lipid exposure can act directly on intestinal stem cells to promote the emergence of the AQP10⁺ enterocyte state, suggesting a metabolic adaptation tuned to dietary components more prevalent in the US diet.

### Enteroendocrine Lineage Diversity and a Developmentally Emergent TRH Hormonal Circuit

Although enteroendocrine cells (EECs) comprise only a small fraction of the intestinal epithelium, they are central sensors of luminal nutrients and orchestrators of hormone release that regulate digestion, metabolism, and gut–brain communication^90,91^. To define which EEC subsets in the duodenum actively detect dietary components, and how these populations vary across environments and with age, we profiled 2,053 EECs from 88 participants (**Figure 6A**). To our knowledge, this represents the largest collection of unmanipulated human duodenal enteroendocrine cells profiled to date.

**Figure 6:**
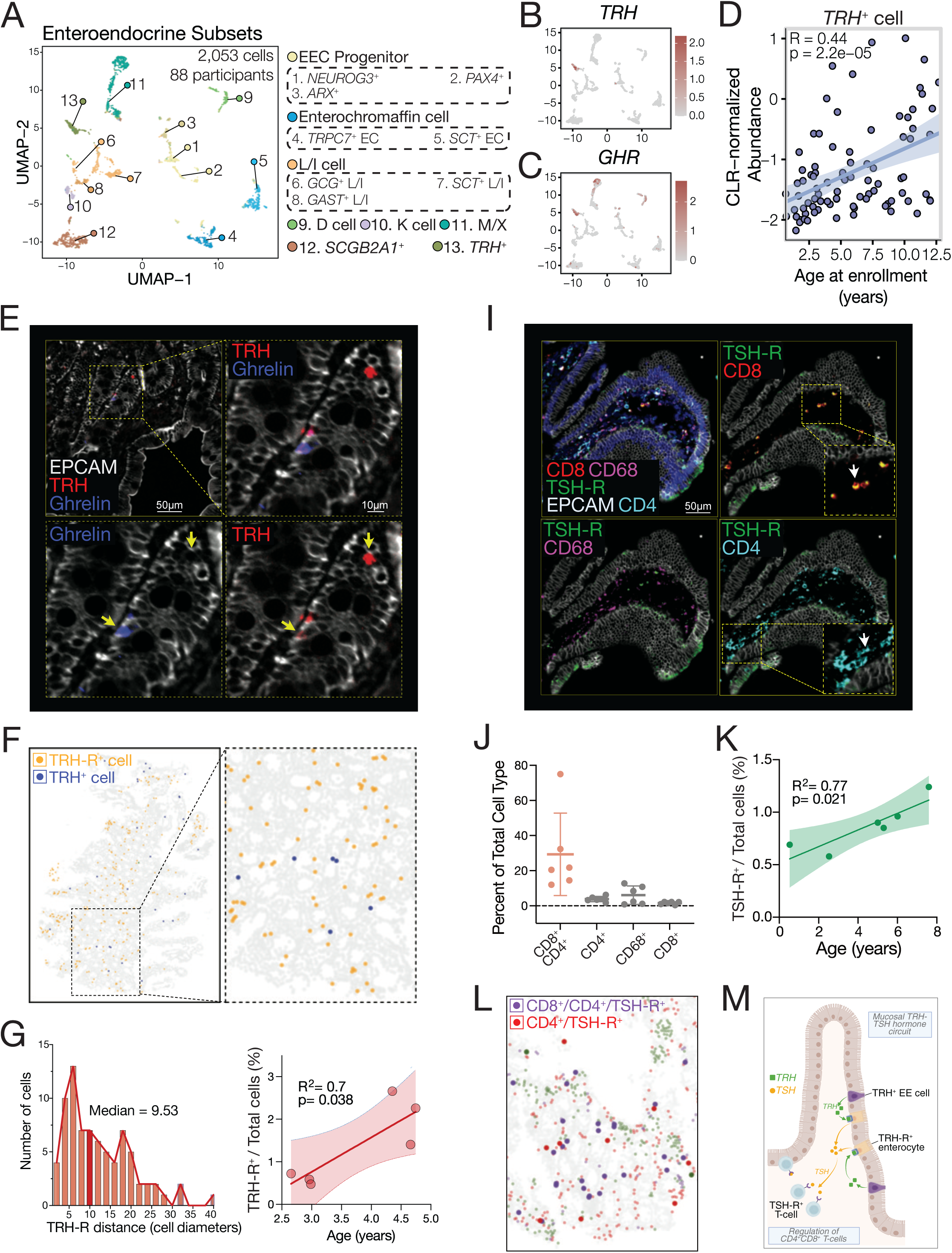
Identification of novel TRH^+^ enteroendocrine cells in the pediatric duodenum. **A.** UMAP representation of 2,053 enteroendocrine cells from 88 participants colored by cell state identity. EEC, enteroendocrine cell; EC, enterochromaffin cell. **B and C.** UMAP display of *TRH* (**B**) and GHR (**C**) expression in enteroendocrine cells. **D.** Normalized abundance of *TRH*^+^ EECs across participant age. Correlation coefficient and statistical significance was calculated using the Spearman rank correlation. CLR, centered log ratio. **E.** Representative MxIF image of pediatric duodenum (Boston, Male, 3.0 years-old) stained with EPCAM (white), TRH (red) and Ghrelin (blue). Yellow arrows highlight TRH^+^ EECs and dotted boxes denote enlarged insets. **F.** Representative pointillistic image of a biopsy from the United States (Virginia, Male, 4.75 years-old) showing spatial relationship of TRH-R^+^ epithelial cells (yellow dots) and TRH^+^ EECs (blue dots). Inset on the right shows an expanded view of the dotted box in the pointillist image. **G.** Quantification of the distance between TRH^+^ EECs and nearest TRHR^+^ epithelial cells measured in cell diameters. **H.** Linear association between participant age and TRHR^+^ cells as a percent of total cells detected by MxIF. **I.** Representative MxIF image (Pakistan, Female, 0.5 years-old) stained with TSH-R (green), EPCAM (white; epithelial cells), CD68 (purple; macrophages), CD4 (teal; T cells) and CD8 (red; T cells). Panels depict all stains (top left), CD8 T cells and TSH-R (top right), macrophages and TSHR (bottom left), and CD4 T cells and TSH-R (bottom right). **J.** Percentage of TSH-R^+^ immune cells within the total population of that cell type. **K.** Linear association between participant age and TSH-R^+^ cells as a percentage of total cells detected by MxIF. **L.** Representative pointillistic image of a biopsy (Pakistan, Female, 6.0 years-old) stained with CD4, CD8 and TSH-R. Purple points correspond to TSH-R^+^ DP T cells, and red points to TSH-R^+^ CD4^+^ T cells. **M.** Proposed model of TRH^+^ EEC – enterocyte – T cell communication axis. TRH^+^ EECs release TRH, which binds to TRH-R on epithelial cells to stimulate TSH release by epithelial cells. TSH then binds to TSH-R^+^ T cells.

Classical hormone-based markers identified serotonin-producing enterochromaffin cells (*TPH1*⁺, Enterochromaffin cells), somatostatin-producing D cells (*SST*⁺), and glucose-dependent insulinotropic peptide–producing K cells (*GIP*⁺). We also detected multi-hormone subsets, including ghrelin/motilin–expressing M/X cells (*GHRL*⁺/*MLN*⁺) and *CCK*/*PYY/GCG-*expressing L/I cells (**Extended data 6A**). Secretin (*SCT*) was expressed in both Enterochromaffin and L/I cells^30,92–95^.

Analysis of nutrient-sensing machinery revealed that long-chain fatty-acid receptors *FFAR1* and *FFAR4* were enriched in GLP1⁺ K cells and CCK/PYY⁺ L/I cells. *FFAR1* was also expressed by a previously undescribed NEUROD1⁺ EEC subset marked by *SCGB2A1* and the succinate receptor *SUCNR1.* EECs broadly expressed bile acid–sensing pathways, including marked upregulation of the amiloride-sensitive bile acid channel ASIC5 in GIP⁺ K cells^96^. Carbohydrate transporters (SGLT1/*SLC5A1*, GLUT2/*SLC2A2*, GLUT5/*SLC2A5*) were widely expressed, whereas GLUT1/*SLC2A1* was restricted to progenitors and subsets of Enterochromaffin and L/I cells. Peptide transporters PEPT1/*SLC15A1* and SNAT1/*SLC38A1* were similarly broadly expressed across EECs. Consistent with prior studies, the calcium-sensing receptor CaSR was enriched in L/I, D, and K cells (**Extended data 7A**)^29,97^. Glutamate receptors, which detect both dietary- and neuron-derived glutamate, showed a high degree of specificity to EEC subsets. Together, these data provide a cellular atlas of nutrient and neurotransmitter sensing across the pediatric duodenal EEC lineage **(Extended data 7B–D).** Further work will be needed to understand how individual receptors, channels, and transporters shape hormone release and EEC function.

In addition to mature enteroendocrine cells, we also identify progenitor populations positive for the transcription factor *NEUROG3*, which is transiently expressed during fate-specification and downregulated in mature cells^30,98^. These progenitors can be sub-divided into Aristaless-related homeobox^+^ (*ARX*^+^) and paired box 4 (*PAX4*^+^) cell subsets. *ARX^+^* progenitors represent a precursor of neuropeptide positive cells as *ARX* is expressed in all EEC subsets containing at least one neuropeptide. In contrast, PAX4^+^ cells likely represent an enterochromaffin cell precursor^99^.

Our extensive enteroendocrine dataset revealed that duodenal EECs express a range of hormones typically associated with extra-intestinal functions. We observed high levels of Parathyroid hormone-like hormone (*PTHLH*) in EC cells, Angiotensinogen (*AGT*) in M/X cells, and Growth hormone-releasing hormone (*GHRH*) in a subset of CCK⁺ L/I cells (**Extended data 6A**). These hormones indicate a role for enteroendocrine cells in regulating bone development and growth. However, the likelihood of a non-canonical role for these hormones cannot be excluded.

Notably, we also identified a previously undescribed EEC state marked by expression of thyrotropin-releasing hormone (*TRH*), a hormone classically restricted to the hypothalamus (**Figure 6B**)^100^. TRH⁺ EECs co-expressed transcripts encoding synaptic adhesion and guidance molecules (*SYT10, CDH10, NCAM2, NTNG1*) as well as Growth hormone receptor (*GHR*), suggesting potential interaction with the enteric nervous system and a link to growth-related pathways (**Figure 6C, Extended data 6C**)^101–103^. TRH⁺ EECs were present in most participants (62.3%) with a mean frequency of 5.8% of duodenal EECs, and their abundance increased significantly in children older than five years (**Extended data 6D, Figure 6D**).

MxIF confirmed TRH protein production in a subset of CHGA⁺EPCAM⁺ EECs, including occasional dual-positive TRH⁺GHRL⁺ cells (**Figure 6E, Extended data 8A and B**). In the hypothalamic–pituitary axis, TRH acts on TRH receptors (TRH-R) in the pituitary to induce secretion of Thyroid Stimulating Hormone (TSH), which then stimulates thyroid hormone production^100^. We therefore assessed expression of TRH pathway receptors in the duodenum. TRH-R was readily detected in a substantial fraction of EPCAM⁺ epithelial cells (**Figure 6F**), and spatial neighborhood analysis demonstrated that TRH-R⁺ epithelial cells lie in proximity to TRH⁺ EECs (median distance 9.53 cell diameters; **Figure 6G**). As with TRH⁺ EECs, epithelial TRH-R expression increased with age (**Figure 6H**). We also found that TSH-R, the receptor for TSH, is expressed by immune cells within the lamina propria and is particularly enriched in CD4⁺/CD8⁺ double-positive T cells (**Figure 6I, J**). TSH-R⁺ T cells, like TRH⁺ EECs and TRH-R⁺ epithelial cells, increased with age and were broadly distributed throughout the duodenal lamina propria (**Figure 6K, L**).

Together, these findings reveal an unexpected, age-associated duodenal hormonal circuit linking TRH-producing EECs, TRH-R⁺ epithelial cells, and TSH-R⁺ T cells. This circuit suggests a previously unrecognized pathway by which growth or developmental cues may interface with the homeostatic regulation of mucosal lymphocyte function (**Figure 6M**).

## Discussion

The Gut-AGE study was designed to define how the cellular architecture of the proximal human intestine is established and maintained during early childhood. By profiling participants from 6 months to 13 years of age across distinct geographic and environmental contexts, and by integrating extensive participant-level metadata with single-cell transcriptomics and spatial data, we chart duodenal maturation. Across sampled geographies, the first four years of life is a period of immune priming during which we observed a transition from naïve and innate lymphocytes to mature memory populations. However, immune cell populations differ across geographic environments, with IELs being more abundant in individuals from Pakistan than the US. Our findings document several previously undescribed cell states, including a lipid-absorptive AQP10^+^ late enterocyte, and an age associated TRH^+^ EEC. Altogether, we have generated a public resource that captures the landscape of the pediatric duodenum across age, ancestry and environment.

At single-cell resolution, most mucosal cell types and states were compositionally stable across age and geography, highlighting the robustness of early intestinal development. Nevertheless, the size and diversity of our cohort revealed previously unrecognized discrete, age-dependent and environmentally influenced modules of mucosal organization. For example, despite stable overall fibroblast and macrophage abundance, their spatial proximity within the villus lamina propria was elevated during the first four years of life. This association subsequently decreased later in life, suggesting coordinated maturation of stromal–immune interactions during early childhood.

Our data also capture a rapid reorganization of the lymphocyte compartment during the first four years of life, marked by a shift from naïve and innate-like subsets toward antigen-experienced phenotypes. This pattern is consistent with prior observations throughout the small and large intestine^14,104,105^ and likely reflects early exposure to dietary and microbial antigens. Given that naïve cells seed the intestine prenatally and that both microbial colonization and weaning drive immune maturation in murine models, the early transition observed here likely signifies the establishment of nascent adaptive immune surveillance in human infancy^34,35,106–110^.

Environmental context also shaped mucosal composition. Children enrolled in Pakistan displayed higher levels of induced CD8^+^ IELs and variation in enterocyte subsets relative to participants in the US, in the absence of pathological activation. Elevated IEL abundance is consistent with their proposed role in epithelial surveillance, particularly in settings with greater exposure to enteric pathobionts, reflected here by frequent colonization with *H.pylori* (58%) and *Giardia* (22%) in participants in Pakistan. As induced IELs arise in an antigen-dependent manner, these observations may indicate physiological adaptation to local microbial ecology^59^.

We identified a diet-associated enterocyte state defined by expression of *AQP10*, a human glycerol transporter that operates optimally in the acidic post-prandial duodenum^78,111^. This AQP10^+^ absorptive program was enriched across US sites and associated with dietary patterns including higher fat intake and consumption of highly processed foods. The over-representation of apolipoproteins and triglyceride metabolic pathway genes in AQP10^+^ enterocytes suggest a critical function in fat digestion. Organoid experiments indicate that lipid-mediated signaling in epithelial stem cells promotes this state, suggesting that responses to diet shape absorptive lineage allocation. However, lipid-induced adaptation responses were donor-specific, suggesting that a stem cell-centered epigenetic memory mechanism may link lineage decisions to prior dietary environments. These findings highlight a possible environmentally responsive epithelial program with direct relevance to nutritional health across diverse settings.

A key strength of the dataset is the unprecedented representation of enteroendocrine cells, which constitute only ∼1% of the intestinal epithelium. Consequently, they have previously been profiled in detail using organoid systems and genetic mouse models rather than native human tissue^28–30,93^. Our study provides the largest collection to date of unmanipulated human duodenal EECs profiled at single-cell level with corresponding spatial localization information. We uncovered a previously unrecognized endocrine circuit in the intestine, centered on a TRH-expressing enteroendocrine population. MxIF confirmed TRH^+^ EECs in the duodenal epithelium, and we observed expression of TRH receptors (TRH-R) on neighboring enterocytes and TSH receptors on nearby CD4^+^/CD8^+^ T cells. Given the short serum half-life (approximately 20 minutes) of TRH, these components likely function locally, raising the possibility of the duodenum co-opting select components of the hypothalamic–pituitary–thyroid axis^112^. Prior murine studies showing TRH-induced TSH production in intestinal epithelium and reduced IEL abundance in TSH-R-deficient mice support the plausibility of a local endocrine–immune circuit that develops during early childhood to regulate local lymphocyte numbers in response to growth and nutrient availability^113,114^. Overall, we propose that TRH^+^ enteroendocrine cells are a previously undescribed and distinct subset of enteroendocrine cells.

Despite its breadth, our study has several limitations that warrant consideration for future investigation. Differences in clinical indications and procedural protocols between sites could, in principle, introduce compositional bias unrelated to biology, although we observed no evidence that such factors influenced our results. Our cohort, though geographically distributed, represents only a subset of global environments, and longitudinal sampling will be required to disentangle temporal trajectories from inter-individual variation. Functional characterization of the inferred cellular circuits such as the putative TRH–TSH mucosal axis and diet-responsive AQP10^+^ cell state will require further mechanistic studies in primary human systems or model organisms capable of recapitulating these pathways.

Our data provides a cellular and spatial framework for understanding how the human duodenum matures across early childhood and adapts to the individual’s lived environment. They reveal conserved features of mucosal development, while identifying context-specific circuits and niches that may mediate growth, immune maturation and diet-dependent physiology. Expanding Gut-AGE to include longitudinal profiling, deeper metabolomic characterization, and integration with clinical, socio-economic and exposure data will be critical for understanding and confirming the links between environment and mucosal function suggested by our data.

## Methods

### Data Availability

Final annotated objects are available at the CellXGene portal (https://cellxgene.cziscience.com/collections/76642a11-3495-4a26-a670-f28936c4b093) and will be available through the Human Cell Atlas Initiative. Furthermore, final single-cell RNA-seq objects, MxIF images and supporting resources are also available through a dedicated project website (www.gut-age.com) for data visualization and exploration.

All original code is available as of the date of publication. All figures were generated and analysis was performed in R (v4.5.0) or Python v3.12.7 using third party packages highlighted throughout the methods, as well as standard analysis tools including conda (v24.9.2), numpy (v2.0.2), pandas (v2.2.3), opencv-python (v4.10.0.84), scikit-image (v0.24.0), scikit-learn (v1.5.2), scipy (v1.14.1), cellpose (v3.1.0), matplotlib (v3.9.4), tifffile (v2024.12.12), napari (v0.5.6), jupyter (v1.1.1), jupyterlab (v4.3.3), openpyxl (v3.1.5), plotly (v6.2.0), ipykernel (v6.29.5), napari-segment-blobs-and-things-with-membranes (v0.3.12), pip (v24.2), seaborn (v0.13.2), torch (v2.5.1), tqdm (v4.67.1), vitessce (v3.5.8) and the Tidyverse collection (v2.0.0). Any additional information required to reanalyze the data reported in this paper is available from the lead contact upon request.

### Participants and Sample Collection

#### Study Sites and study populations

The GutAGE study has four recruitment sites: Aga Khan University (AKU; Karachi, Pakistan), the University of Mississippi Medical Center (UMMC; Jackson, MS), the University of Virginia (UVA; Charlottesville, VA) and Boston Children’s Hospital (BCH; Boston, MA). These sites were selected to include children living in rural and urban environments across a range of resource and developing regions with varying climates. Sites were chosen to provide an opportunity to study children of diverse ancestry who have not historically been included in genetic research. Aga Khan University (AKU; Karachi, Pakistan) serves patients from across Pakistan including all local ethnic groups. The University of Mississippi Medical Center (UMMC; Jackson, MS) serves patients across Mississippi, including from rural communities with limited local healthcare resources. The African American population is genetically diverse and there are also sizeable Hispanic/Latin-X and Native American populations. The University of Virginia (UVA; Charlottesville, VA) serves patients from the entire state, as well as from rural areas in West Virginia and Tennessee. Boston Children’s Hospital (BCH; Boston, MA) serves patients from across Massachusetts, as well as from the greater New England region, other areas of the US, and those who seek care from other countries.

#### Ethics

This study was approved by the Institutional Review Board at BCH (IRB-00040075), upon which UMMC (2021V0695) and UVA (IRB-HSR210435) relied, and by the Ethical Review Committee at AKU (2023_7018_23972). Parent/Guardian consent was obtained for all participants. Assent was granted by participants above 7 years of age prior to any study procedures. At U.S. sites, participants were provided $50 for their participation. At AKU, the procedure cost and follow-up consultation fee were paid by the study.

#### Participant Selection

Children undergoing diagnostic esophagogastroduodenoscopy (EGD) for clinical indications with a high likelihood of being non-diagnostic for active mucosal disease (e.g., foreign body retrieval, non-specific abdominal pain, and reflux) and whose family were able to complete the survey in a language spoken by the study team were eligible to participate. Exclusion criteria were: chromosomal or genetic abnormalities, gastrointestinal disease other than eosinophilic esophagitis, abnormal laryngopharyngeal or gastrointestinal anatomy, neurodevelopmental disorder, coagulopathy or bleeding disorder, and connective tissue disorder. Samples from any participant with a clinical biopsy pathology report indicating disease (e.g., celiac disease, Crohn’s disease) were excluded from further analysis.

#### Participant Recruitment

Referral lists and clinic and procedure schedules were used to identify potentially eligible children. Those who were still potentially eligible after medical record review were contacted by the study team in conjunction with a previously scheduled gastroenterology clinic appointment, by telephone, or on the day of the endoscopy procedure. A research assistant explained the project to the family and obtained consent from the parent(s)/legal guardian(s) and assent from the participant when appropriate.

#### Data and Biospecimen collection

Clinical research assistants conducted a structured medical record review to extract indications for endoscopy, anthropometrics, and select laboratory studies. Interviews were conducted to obtain detailed participant medical, dietary, social and environmental history, as well as family history and self-reported ancestry. The complete participant registration form is available at dx.doi.org/10.17504/protocols.io.6qpvr4d3zgmk/v1^115^. Blood (8.5 mL) was collected along with up to 4 endoscopic mucosal pinch biopsies obtained by and at the discretion of the endoscopist performing the clinically indicated EGD procedure. Biopsy forceps were not standardized as they were not provided by the study.

#### Biopsy processing for live cell isolation, including scRNA-seq and organoid generation

One to two biopsies were immediately placed in 1mL Collection Media (Advanced DMEM/F12 1X media, 10% FBS, Penicillin/Streptomycin) on wet ice for transport to the local laboratory, where they were transferred into cryopreservation media (90% FBS, 10% DMSO) within 1 hour of collection then immediately slow frozen to −80°C in a Mr. Frosty™ (Thermo Fisher Scientific Nalgene, Waltham, MA).

#### Histopathology Slide preparation and digitization

One biopsy was fixed in 10% buffered neutral formalin for 4-24 hours at room temperature, then washed with 70% ethanol twice prior to storage in 70% ethanol at 4°C until paraffin embedding. Formalin-fixed biopsies were paraffin-embedded locally at AKU, BCH and UVA. BCH also embedded samples for UMMC. Embedded tissue was shipped to the University of Virginia (UVA), where each biopsy was oriented to allow for visualization of duodenal villi as confirmed by an expert gastrointestinal pathologist. To maximize efficiency during histology analyses, biopsy cores were taken from the embedded biopsies and placed onto tissue microarrays (TMA). Biopsies from BCH, UMMC and UVA were distributed across TMAs as much as possible given variable enrollment and accrual and each 2×3 array included biopsies from at least two centers. Biopsies from AKU were mounted together to facilitate the repatriation of unused tissue to Pakistan. Six 4 µm-thick slices from each microarray were each mounted on standard glass microscopy slides for immunofluorescence and hematoxylin and eosin (H&E) staining. Five unstained slides were dipped in paraffin before shipment to BCH for immunofluorescence studies. One hematoxylin and eosin (H&E)-stained slide was digitized at 40× magnification with a spatial resolution of 0.23 µm per pixel in both the X and Y directions using a Hamamatsu NanoZoomer S360 (C13220-01) at UVA.

### Whole genome sequencing and Ancestry imputation

Ancestry imputation was performed using Gencove’s ancestry deconvolution service^116,117^. For the majority of participants, snap-frozen whole blood aliquots were sent to Gencove and genomic DNA (gDNA) was isolated. For a small number of samples, gDNA was isolated on site at BCH and gDNA was delivered to Gencove for further processing. gDNA was sequenced using low pass whole genome sequencing (∼7X coverage). Variant calls were then imputed using Gencove’s proprietary pipeline and imputed variants were compared to known variant reference populations to estimate global ancestry admixtures.

### RNA-seq Methods

#### Tissue dissociation for single-cell and bulk RNA-seq

Biopsies processed for live cell processing were shipped frozen to BCH where they were processed for bulk and single cell RNA sequencing. First, biopsies were rapidly thawed at 37°C for 90 seconds then rinsed in 3 mL Hank’s buffered salt solution to remove residual DMSO. Next, whole biopsies were finely minced using a razor and tissue was digested in 10 mL complete RPMI (RPMI + Glutamax, 2% FBS, 10 mM HEPES, sodium pyruvate, non-essential amino acids and 1X penicillin:streptomycin) containing 100 ug/mL Liberase^TM^ TM and 100 ug/mL DNase I. Following 30 minutes at 37 °C with shaking at 220 rpm, samples were placed on ice and digestion was quenched using 100 µL 0.5M EDTA (pH 8.0) and 100 µL FBS. Digested samples were homogenized using a wide-bore 1mL pipette followed by a 10 mL serological pipette then passed through a 40µm filter. Cells were centrifuged at 500x*g* for 5 minutes, then resuspended in 300uL complete RPMI prior to counting via hemocytometer with Trypan Blue staining (1:1). Prior studies have used dissociation protocols that separate the epithelial layer from the underlying lamina propria^26,118,119^. In our experience, these methods were not suitable for the isolation of cells from cryopreserved biopsies. Therefore, this protocol was optimized to obtain cell yields suitable for scRNA-seq from frozen intestinal biopsies, while retaining cellular representation from both epithelial and lamina propria fractions.

#### Droplet-based single-cell RNA sequencing

For scRNA-seq all samples were processed and loaded at BCH. For each batch, 30,000 cells per individual participant were combined into pools of 3 individuals each. From each pool, a maximum of 30,000 cells were loaded into a single lane of a 10X 3’ Chromium Next GEM Single Cell 3’ v3.1 chip. Libraries were prepared using the corresponding kit with dual indices as per the manufacturer’s instructions. Library quality was verified by TapeStation 4200 (Agilent). Sequencing was performed at the Broad Institute Sequencing Core using either the NovaSeq 6000 or Novaseq X platform (Illumina), with a minimum targeted RNA read depth of 20,000 reads/cell **(Supplementary data 1).**

#### Bulk RNAseq

For each biopsy, 15,000 cells were lysed using 50 µL RLT buffer (Qiagen, RNeasy Kit) + 2-mercaptoethanol (1% v/v). Lysates were snap frozen on dry ice then stored at −80°C for batch processing. Libraries were prepared using the previously described SmartSEQ2 protocol^120^. Briefly, RNA was purified from 10 µL of cell lysate using 2.0X SPRIselect beads (Beckman-Coulter). Purified RNA was reverse transcribed into cDNA and whole transcriptome amplification was carried out. Libraries were fragmented and dual indexed using the Illumina Nextera XT Library Prep Kits and sequenced either on the NextSeq 2000 or NovaSeq 6000 platforms (Illumina). Sequencing was performed at the Broad Institute Sequencing Core.

#### Library pre-processing and variant calling

scRNA-seq libraries were demultiplexed and aligned to the GRCh38 human reference genome using the CellRanger toolkit (v8.0.1) provided through Cumulus tools (https://cumulus.readthedocs.io/en/stable/)^121^. Briefly, raw BCL files were demultiplexed using the cellranger mkfastq function, a wrapper of Illumina’s bcl2fastq. Afterwards, FASTQs were aligned to GRCh38 human genome and features were counted using the cellranger count function.

Following alignment, sample quality was evaluated based on CellRanger quality metrics and outliers were removed. In total 3 samples were omitted from further analysis because libraries were of low quality relative to all other samples.

Raw bulk RNA-seq libraries were demultiplexed using Illumina’s bcl2fasq package (v2.20). Participant-identifying SNPs were called using Genome Analysis Toolkit (GATK4) workflows provided by the Broad Institute (https://github.com/gatk-workflows) using default settings. Briefly, paired FASTQs were converted into unmapped BAM files and aligned to the GRCh38 genome. Duplicate marking and base recalibration was performed and haplotype caller was run to generate a variant call file (VCF) for each participant.

#### Genetic demultiplexing

SNP-based genetic demultiplexing of pooled scRNA-seq samples was performed as previously described^43,122,123^. Briefly, scRNA-seq pools were initially demultiplexed in a genotype-agonistic method through the Freemuxlet variation of the Popscle package (https://github.com/statgen/popscle). In summary, Freemuxlet performs unsupervised clustering of single cells based on SNPs identified within transcripts and assigns cells as a nameless donor or doublet. In addition to assigning cells, a VCF is generated for identifying SNPs of each donor. These VCFs are then compared to participant-identified VCFs generated from individual bulkRNA-seq and matched based on genotype similarity (vcf-match-sample-ids, https://github.com/hyunminkang/apigenome).

Patient-identified single cells are then paired with clinical metadata and inter-participant doublets are filtered from downstream analysis.

#### Technical variation

Great care was taken to limit the correspondence between technical variables and biological variables of interest. The three participants pooled into each technical batch were selected such that each batch contained a mix of ages, study sites and sex. Therefore, integration can be performed across the batch ID, which will integrate technical variation while being completely orthogonal to biological variation. To remove the residual effect of technical variation on our downstream analysis, we performed Harmony (v1.0) integration using default parameters across the pool ID variable^124^. Harmony integration was performed at each level of sub-clustering.

The impact of technical and biological variation on the GutAGE dataset was evaluated using previously published methods^125^. Briefly, linear regression was performed for each principal component against major technical and biological variables. The amount of variation explained by each technical variable is summarized in **Supplementary data 2.** In addition, Spearman’s correlation and Theil’s U was calculated between major biological and technical variables. Overall, technical variables were found to have little effect on the variation and structure of our dataset (**Extended data 1A, Supplementary data 2**).

#### Clustering and annotation

All downstream analysis was performed using Seurat (v5.1.0)^126^. Briefly, all participants were merged into one dataset, and an initial broad filtering was performed based on Unique Molecular Identifier (UMI) count (>250 & <5000), number of detected genes (>250) and percentage of mitochondrial genes (<75%). Cells were then normalized using scTransform (v0.4.1), followed by principal component analysis (PCA), Harmony integration, Louvain clustering and UMAP embedding. Different PCs and clustering resolutions were sampled to determine ideal resolution. Using a final resolution of 0.4 and 40 PC values, cluster markers were calculated and major cell lineages were defined based on known lineage markers (eg., *EPCAM*, *PTPRC, TRAC, CD79A, PECAM1, COL1A1,* etc.). Intra-sample doublets were identified and removed based on the co-expression of 2 or more conflicting lineage markers.

To identify fine-grained cell subsets and cell states, iterative clustering was performed as previously described^119,127^. Briefly, each broad cluster was separated, and finer grained filtering was performed with a UMI count and percent mitochondrial cutoff specific to each broad identity. This is necessary as broad cell types have distinct distributions in their RNA content. Therefore, a cutoff that may filter low quality cells of one type would filter out the majority of high-quality cells of another. Following filtering, cells were re-clustered and cell subsets were annotated based on prior knowledge, expert opinions, and previously published small intestine scRNA-seq datasets^1–3,5,6,8^. Iterative clustering also allowed us to identify and remove doublets composed of two distinct cell subtypes within the same cell type. Cells were annotated and doublets were removed. Finally, each annotated cell subtype was subsetted and clustered one additional time to identify cell states. Cell states were annotated by pairing the top differentially expressed marker gene with the cell subtype identity. Overall, we identified a total of 128 end cell states across 55 cell subtypes. Annotations were collectively agreed upon by multiple experts, and the identifying features of each cell state is summarized in **Supplementary: cell glossary**. To visualize the relationship of each cell state, a “phylogenetic tree” was constructed using the ARBOL package^119^. For each cell state, cosine distances to all other cell states were calculated based on the feature-centroid of each state in unreduced gene space. Hierarchical clustering was then performed to build a clustered tree with branch length showcasing the distance in gene expression space between each annotated cell state **(Figure 2B)**.

#### Differential gene expression

Differential gene expression (DGE) was performed at both the single-cell and the pseudobulked level. Single-cell DGE was performed using the Seurat FindMarkers function to identify cell markers for annotation. When performing FindMarkers, DGE was calculated using a Wilcoxon Rank Sum test and significant genes were filtered to include only genes expressed in a minimum of 30% of cells and with a log fold change of at least 0.5.

For all other comparisons, DGE was calculated on a pseudobulked level. Pseudobulked DGE was performed by first aggregating the gene expression of individual cells for each cell state:participant combination. After aggregation, differential gene expression was performed using the DEseq2 package set to default parameters.

#### Compositional analysis

Compositional analysis was performed as previously described^127^. After annotation of all cells and the removal of both inter- and intraparticipant doublets, cell state abundance was calculated for the entire biopsy as well as for each broad cell identity. Initially cell state frequencies were calculated for each participant by dividing the number of cells for each cell state by either the total number of cells in that biopsy or within the broad identity. A pseudocount was then generated by multiplying each frequency by 3000 (Cp3K), as this was approximately the average number of cells per sample (average=2580 post-filtering). Pseudocounts were then transformed relative to the geometric mean using a center-log ratio (CLR). Principal component analysis of sample composition was performed using the CLR transformed data.

The diversity of each sample was calculated using metrics previously employed in biodiversity analysis, including species richness, Shannon index of diversity, and the Gini-Simpson index of diversity^128,129^. Each cell state was treated as an individual species and by extension each biopsy could be viewed as a unique ecosystem. Species richness is a measure of the number of unique cell states observed in each biopsy. Both Shannon’s and Gini-Simpson’s index of diversity were calculated using the cell state frequency within the total biopsy.

#### Pseudotime analysis of enterocytes

The pseudotime trajectory of epithelial cells was calculated using the Monocle3 package (v1.4.26)^70^. Epithelial progenitor and enterocyte populations were initially subsetted and Louvain clustering, as well as UMAP coordinates were recalculated based on the populations present. A pseudotime trajectory was then calculated using the LGR5^+^ epithelial stem cell population as the root node. To evaluate the pseudotime distribution of each enterocyte cluster, the pseudotime values for all cells within a cluster were extracted. The distribution was then plotted, alongside the median and interquartile range.

### Self-Supervised Learning-Based Analysis of Duodenal Whole Slide Images

TMA images were digitally split to prepare a dataset of 110 H&E-stained duodenal biopsy whole slide images (WSIs), one per patient. This included participants from BCH (n = 43), UMMC (n = 33), AKU (n = 21), and UVA (n = 13) (**Extended data 4A**). We then developed a self-supervised learning (SSL)-based AI model to extract WSI-level features from this dataset, as described in detail below (**Figure 4A**).

The ultra-high resolution of these WSIs posed computational challenges for Vision Transformers (ViTs) and Convolutional Neural Networks (CNNs), which are not designed to handle inputs of such size directly^130^.Most studies bypass this requirement by using a two-stage methodology, first splitting the WSIs into small patches of manageable size (e.g., 256×256 pixels), then extracting patch-level embedding using a frozen neural network pre-trained on a large dataset, followed by aggregating the patch embeddings to generate a slide-level representation for each WSI using Multiple Instance Learning (MIL) classifiers trained on slide-level labels (e.g., diagnosis, disease subtype, or treatment response). However, in this study, there were no such labels available to employ supervised or weakly supervised learning approaches for slide-level representation.

Therefore, we adopted the alternative pipeline proposed by Song et al. (2024) to address the absence of labels (**Figure 4A)**^131^. In Stage 1, we fine-tuned a large Vision Transformer (ViT) using a self-supervised learning (SSL) method to extract 1024-dimensional patch embeddings. In Stages 2–3, we applied a GMM-based framework inspired by Panther to aggregate these patch embeddings into compact WSI-level representations^131^.

#### Patch-level Embedding Extraction

To obtain patch embeddings, we utilized UNI, a ViT-L/16 DINOv2 model with 1024-dimensional embeddings, pretrained on a large internal histology dataset as shown in Stage 1 (**Figure 4A**)^54–56^. Use of UNI for patch-level embedding extraction in histopathology allowed us to utilize domain-specific encoders trained on large histopathology datasets. This enables improved performance in capturing domain-specific visual patterns as opposed to traditional ImageNet-pretrained networks^54,132^.

We further fine-tuned this model on our duodenum dataset using the DINOv2 self-supervised learning approach, without the need for labeled data^56^. After fine-tuning, we froze the ViT-L/16 model and extracted 1024-dimensional embeddings for each patch across all WSIs.

#### Whole Slide Image (WSI)-level Feature Extraction

To obtain compact, unsupervised WSI representations, we adopted a Gaussian Mixture Model (GMM)-based approach, inspired by Panther^131^. In Stage 2 **(Figure 4A**), we applied K-means clustering to the patch-level embeddings across all WSIs to determine the initial cluster centroids. In Stage 3 (**Figure 4A**), a GMM was fitted per WSI, with each mixture component representing a distinct histological feature. The number of components was set to 8, as determined by the elbow method. This enabled WSI-level feature aggregation, allowing for the characterization of histologically distinct prototypes within the biopsy samples.

### MxIF Methods

#### Deparaffinization, Rehydration, Antigen Retrieval and Blocking

Unstained slides were deparaffinized at 60°C for 15 minutes, then treated with Histo-Clear (three separate 5-minute incubations, followed by at least four 10-minute incubations). They were then rehydrated using a series of aqueous ethanol solutions with decreasing concentrations (100%, 100%, 95%, 75%, 50%, and 0%), each for 5 minutes at room temperature. Slides were then incubated in antigen retrieval buffer and heated to 100°C for 20 minutes. Tris-EDTA pH 9.0 (Genemed, 10-0046) was used for panels that contained TSH-R and immune markers. Citrate buffer pH 6.0 (Abcam ab93678) was used for all other antigens. After cooling to room temperature for 20-40 minutes, slides were washed twice with PBS and incubated for 1 hour in blocking buffer [0.1% Triton-X-100 (Sigma-Aldrich, cat no. X100-100ML) and 5% bovine serum albumin (Sigma-Aldrich, cat no. A6003-25G) in 1x PBS (Corning, cat no. 21-040-CV)]. Finally, slides were washed twice with PBS before long-term storage in PBS at 4°C.

#### Imaging

Tissue microarrays were imaged using SPECTRE-Plex and our previously published method of custom cyclical immunofluorescence protocols^48^. Briefly, tissues are labeled with fluorophore conjugated antibodies, imaged, then the fluorophores are inactivated by incubating with meta-chloroperoxybenzoic acid. The entire protocol is conducted using a previously published automated custom Python program. The details about the antibodies used in our MxIF protocols are listed in **Supplementary table 11 and 12**.

### MxIF Image Processing

We have refined our previous image processing pipeline with two novel methods: HDR intensity bracketing and updated background subtraction.

#### HDR Intensity Bracketing

There are no definite solutions for autoexposure algorithms that are agnostic of fluorescence marker properties making it extremely difficult to determine exact exposure time for optimal dynamic range. Therefore, we modified a previously published strategy, HDR autoexposure, to increase the dynamic range of the image irrespective of the actual fluorescence properties of the channel to ensure at least one image for each fluorophore with sufficient, non-saturated exposure^133^. Specifically, we extrapolate pixel intensities to the highest exposure time taken and use simplified curves based on relative extrapolation errors and Poisson error based on the generalized steps in the HDR algorithm:

1. Take a number of images at different exposure times
2. Extrapolate all image intensities from their exposure time to that of the highest exposure time taken.
3. Generate weight curves based on relative error due to Poisson error and extrapolation error. In general, we want the highest weight on the brightest, non-saturated pixels that were extrapolated the least amount as that contains the most accurate information.
4. Merge all X images together using the weight curve to generate a single 32bit HDR image.

For brevity, we assume that with a given exposure time, pixel information accuracy (i.e., signal to noise ratio) is ∝ √*I*. For extrapolation, we use a linear camera response function that includes an exposure time offset that was specific to the camera used in the experiment.

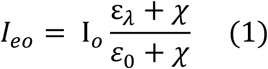

where Ie_o_ is the extrapolated pixel intensity at exposure time ε_λ_ and Io is the pixel intensity at exposure time ε_0_ and χ is the exposure time offset.

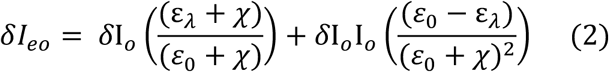

where 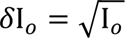

With equation 2 defining the error (noise), the signal to noise *vs* intensity curve can be quickly calculated for each exposure time while preserving the relative intensity of the images.

#### Background Subtraction

We observed in our data that Rolling ball background subtraction introduced significant artifacts, especially if the subtraction radius was not chosen correctly. This was especially prominent when processing a large image dataset that required a different radius for each fluorophore. To overcome this, we adapted a previously published dark frame background subtraction approach agnostic to the spatial particularities of our data^134^. This method generates a dark frame background image by applying a kernel to each pixel that corresponds to the value of the lowest intensity pixel in that kernel (as opposed to the kernel mean in case of rolling ball). Next, a low pass frequency filter is applied in Fourier space to remove any remaining signal from the background, and a dark frame image is generated. This dark frame image is subtracted from the original to obtain the final background subtracted image. We found that this method works well so long as the kernel is large enough to contain a single background pixel. Furthermore, we modified this method to reduce computational intensity. First the image is subsampled using a square pixel grid of 50×50. The lowest value of the pixels is used to determine the new intensity. Given that this image is now down sampled, we used B-spline interpolation to restore the original dimensions. Finally, a low pass filter is applied to the scaled-up background which is then subtracted from original image.

#### Main MxIF Cell Classification

All image processing steps were conducted using a custom Python pipeline in conjunction with standard Python libraries. First, all individual channels were down sampled twofold, normalized, and merged to generate a composite stain image. This normalized image was thresholded by Otsu’s method to create a binary image, which was dilated and the holes were filled using ndimage.binary_fill_holes. The number of tissues in each imaging run was defined manually. Within each tissue boundary, the largest contoured structure(s) were defined as the tissue mask which acts as the boundaries of the image with all subsequent processing conducted on images with the applied tissue mask (**Supplementary data 3)**. A top-hat filter was applied to all stains other than structural markers and bright stains (**Supplementary data 4)**.

Three channels (DAPI, EPCAM, and MUC2) were normalized using Min Max Scaler in the sklearn Python package. DAPI and MUC2 intensities were then averaged to create a composite image for segmentation. Segmentation was conducted using CellPose (Ver 3.1.0) with the normalized EPCAM image and the normalized DAPI and MUC2 composite image being the “chan to segment” and “chan2” inputs, respectively (**Supplementary data 5)**. Cellpose model “cyto3” was used with the default settings to generate cell masks. These cell masks were then dilated using segmentation.expand_labels with a distance of 3. A second set of masks were created specifically for fibroblasts using seeded_watershed from napari_segment_blobs_and_things_with_membranes. Using this pipeline, we observed that we could get segmentation sensitivity of 90% with high specificity.

The masks were used to determine the characteristic cellular intensity of each marker, along with the centroid coordinates for each cell in the image. Centroid coordinates were calculated using scipy.ndimage.center_of_mass. These data were compiled into 2 data frames, one using the original cell masks and the other using the modified fibroblast cell masks. A multi-round Otsu thresholding technique was used to threshold positive cells for each cell identity signal. For EPCAM, the top 10% of pixels within the original dilated cell masks were used, while for smooth muscle actin (SMA), all pixels within the modified fibroblast cell masks were used. For all other stains, all pixels within the original dilated cell masks were used to calculate thresholds. The first step was to clip the top 1 percent of highest intensity pixels (np.quantile(clip_img, q=0.99)). The first round of thresholding was implemented using skimage.filters.threshold_multiostu with 5 classes. Each pixel was assigned a threshold value of 0-4 based on the intensity, with the clipped pixels being 4 (highest threshold). The mean value was calculated using pixels corresponding to each cell. The cut-off threshold for each stain was manually determined and was consistent across all tissues. The second round of thresholding was conducted using only the pixels from cells greater than the specified threshold and using three classes (low, medium, high). The value for each cell was determined by calculating the median value of pixels within the cell. These thresholding outputs were used to determine cell type classifications using decision trees. A top-level decision tree classifies cells into 1 of 4 cell types (Epithelial, Immune, Stromal, Other) as outlined in **Supplementary data 6**. Next the level 1 cells were classified as subtypes or Level 2 classification using a decision tree outlined in **Supplementary data 7.**

For TSH-R, the initial processing steps were the same as the main antibody panel up until cell type classification, except that a composite of only DAPI and EPCAM was used for cell segmentation.

Cell type classification is determined using a set of decision trees. A top-level decision tree classifies cells into 1 of 3 cell types (Immune, Epithelial, Other). First, cells that are immune marker^+^ are classified as Immune cells. Next, cells that are immune marker^-^ but EPCAM^+^ are classified as epithelial cells. The remaining cells are classified as non-classified cells/others.

#### Enteroendocrine Cell MxIF classification

Initial steps were identical to the main multiplex IF image processing, except that a composite image consisting of only DAPI and EPCAM was used for cell segmentation. The thresholding protocol was used to classify enteroendocrine cells using Chromogranin A (CHGA) and EPCAM. For classifying the subtype of enteroendocrine cells, only cells that were both EPCAM^+^ and CHGA^+^ were considered.

The top 0.01 percent of highest intensity pixels (np.quantile(clip_img, q=0.9999)) were clipped and then the top 10% of pixels within each cell mask was used to calculate the mean pixel intensity per cell. Thresholding was performed using Otsu’s method (skimage.filters.threshold_otsu). In two biopsies (one from Mississippi and one from Virginia), one of the imaging regions was removed because of a high fraction of saturated pixels.

Cell type classification was done in two steps. For all stains, except TRH-R, each cell was initially classified by all stains that were above their given threshold (determined to be stain^+^). The classification categories were then consolidated if stain^+^ irrespective of NCAM/SUCNR1 (as these are receptors, not cell type markers). For instance, ghrelin^+^NCAM^-^ and ghrelin^+^NCAM^+^ were both classified as ghrelin positive cells. In the second step, TRH-R was classified using the determined stain threshold and included irrespective of EPCAM.

#### Context Mapping

To identify the tissue context, we have modified our previous work that uses mathematical morphology to classify the different architectures in the intestinal surface for fluorescence images^50^.

Specifically, we used two background subtracted images, DAPI to mark nucleus and EPCAM to mark epithelial cells. Segmentation maps were generated from these two images using different structuring element sizes. Next, ridge-based skeletons calculated using the distance transform from the input image were used to identify the villus-like protrusions and the villus base. Regions within the villus-like protrusions were then classified as villus epithelia (VE) and villus lamina propria (VLP) using the EPCAM segmentation map. This simultaneously enables identification of the tissue regions that are not villus. Next, the structuring element size was chosen to segment the crypt epithelium (CE) in the lamina propria region. The remaining region was classified as crypt lamina propria (LP). Each cell identified as described above in Cell Classification was then assigned to a tissue region.

#### Spatial Analysis

##### Cell Proportions

Cell counts were normalized for each context map region (LP, VLP, CE, VE).

##### Counting Intraepithelial Lymphocytes (IELs)

Cells located within the villus epithelium context map region and categorized under the “immune” cell type were defined as IELs. These include: CD3D CD45, CD3D, CD3D EPCAM(L), CD3D CD4 CD45, CD3D CD8a CD45, CD3D CD4, CD3D CD8a, CD3D CD4 CD8a, and CD3D CD4 CD8a CD45. Biopsies with fewer than 250 cells in the VE were excluded from the analysis. Finally, the ratio of IELs to total villus epithelial cells was calculated.

##### Immune Density

For each context map region (LP, VLP, CE, VE), two data frames were generated to hold cell density values. The first data frame included density values for all cells, excluding those classified as subtype “Other.” The second data frame contained only density values for “immune” cells. All density calculations were performed using the sklearn.neighbors.KernelDensity function, with a bandwidth of 300. These data frames were then combined with df.merge and aligned based on the centroid positions of the cells. The ratio between the “immune” cell density and total cell density was calculated for all cells, and only the entries corresponding to “immune” cells were retained in the final combined data frame. The ratios were categorized into bins with a width of 0.2, covering the range from 0 to 6. The count for each bin was then normalized by dividing by the total number of ratios.

##### Cell Neighborhood

The nearest neighbor of each fibroblast was calculated using the approximate nearest neighbor search algorithm implemented in R with the Wraps ANN Library^135^.

##### Brush Border Analysis

A tissue border mask was generated by sequentially applying an external gradient, a size filter (to remove small objects) and a dilation to the DAPI and EPCAM images. This mask was then integrated with each brush border stain to isolate the brush border signal at the tissue boundary. We then segmented the villus into equal sections along the villus base-tip axis using villus skeletonization part of the context map (see above) and measured the intensity of each stain along this axis.

##### Enteroendocrine Distance

The distance between each TRH^+^ enteroendocrine cell and TRH-R^+^ enterocyte was measured using np.linalg.norm to calculate distances and np.argmin to determine the shortest distance. Distance values were divided by 20 (average cell diameter) to calculate distance as number of cell lengths^136^.

### Organoid Methods

#### Organoid line establishment and maintenance

Human duodenum crypts were isolated from participant biopsy samples using established methods^137^. Isolated crypts were embedded in Growth Factor Reduced Phenol Red Free Matrigel® as 50 μL domes in 24-well plates (Corning Costar) and cultured in stem cell media consisting of RepliGut Growth Media (Altis biosystems, MED-RGM-200), 500 nM A83-01 (Sigma, SML0788), 10 μM Y-27632 ROCK inhibitor (Sigma, Y0503), and 250 nM CHIR99021 (Sigma, SML1046). Organoids were maintained at 37°C in 5% CO₂ with stem cell medium changes every 2 days and passaged every 4 days.

#### Organoid lipid responses

For lipid treatment experiments, organoids were initially cultured in stem cell media for 24 hours post-seeding. Organoids were treated by a 5-day exposure to stem cell medium supplemented with or without a defined lipid mixture (100 μM total concentration) comprised of linoleic acid (Cayman, 38649), palmitic acid (Cayman, 29558), oleic acid (Cayman, 29557), and docosahexaenoic acid (DHA; Cayman, 35872). Some organoids were collected at this stage and the rest were differentiated. To induce differentiation, organoids that had been treated with or without lipid mixture were transferred to RepliGut Maturation Media (Altis Biosystems, MED-RMM-100) and maintained for an additional 5 days.

#### qPCR to measure AQP10 expression

Duodenum organoids embedded in Matrigel® were lysed in TRIzol (ThermoFisher, 15596026) and isolated with chloroform. RNA was precipitated with isopropanol, washed with 75% ethanol, air-dried, and resuspended in RNase-free water. RNA concentration and purity were assessed using a spectrophotometer. For reverse transcription, 1µg of total RNA was converted into cDNA using the NEB cDNA Synthesis Kit (NEB, E6560S). AQP10 cDNA was quantified on a CFX384 real-time cycler (Bio-Rad) with NEB Universal qPCR Master Mix (NEB, M3003L) and primers targeting AQP10. qPCR data was normalized to HPRT1 expression, and results presented as fold change of the lipid supplemented group relative to the matched non-lipid supplemented group. Primer sequences used Human HPRT; (Primer 1: 5’-CATTATGCTGAGGATTTGGAAAGG −3’, Primer 2: 5’-CTTGAGCACACAGAGGGCTACA −3’) and Human AQP10: (Primer 1: 5’-GACAGAGGGAGCAGTGAATAG −3’, Primer 2: 5’-GCCAGAAACATGGTGAAGAAG −3’)

### General Statistical Analysis

For correlation of participant metadata, variables were sorted into continuous, Boolean (yes/no survey data), ordinal (categorical data with ranked values) or nominal (categorical data without rank) data structures. Boolean and ordinal categories were re-encoded into numerical values based on the rank of each category, with True/Yes = 1 and False/No = 0. Nominal data was one-hot encoded such that dummy variables represented each unique category from the original variable. After encoding, correlation was performed across all variables using a spearman rank-order correlation.

For evaluating changes in cellular proportion across categorical variables, such as site, we performed either a Mann-Whitney U test (variables = 2) or a Kruskal-Wallis’s test, followed by Dunn’s post-hoc analysis (variables >2). For evaluating changes across continuous variables, such as age, a linear regression was calculated using the lm function in R. Unless otherwise specified, p values were corrected based on the false discovery rate estimated with the Benjamini and Hochberg correction (Benjamini and Hochberg, 1995), using p.adjust with the fdr method in R.

P value, n and additional summary statistics are provided within the figure itself and/or the corresponding figure legend where appropriate. Data visualization was performed in R, Python or Prism using the following packages: Seurat (v5.1.0), ggplot2 (v3.5.2), ggridges (v0.5.6) and ComplexHeatmap (v2.25.1).

## Acknowledgements

The authors are grateful to the individual participants and families who generously contributed their time and samples, without whom this research would not have been possible.

The authors thank the following endoscopists who collected specimens without whom this study would not have been possible: Adeel ur Rehman – Aga Khan University; Sonia Ballal, Beate Beinvogl, Elana Bern, Silvana Bonilla, Alexandra Carey, Denis Chang, Lauren Collen, Nan Du, Christopher Duggan, Scott Elisofon, Laurie Fishman, Alejandro Flores, Victor Fox, Amit Grover, Elizabeth Hait, Suzanna Hirsch, Lissette Jimenez, Stacy Kahn, Ishrat Mansuri, Maireade McSweeney, Claudio Morera, Melissa Musser, Peter Ngo, Samuel Nurko, Sara Rosenbaum, Eitan Rubenstein, Paul Rufo, Desiree Sierra-Velez, Jared Silverstein, Erin Syverson, Kate Templeton, Michaela Tracy, Andrew Werhman, Dascha Weir, Allison Wu, Jessica Yasuda, Jason Zhang, Lori Zimmerman, Naamah Zitomersky – Boston Children’s Hospital; Michael Nowicki, Sandra Camacho-Gomez – University of Mississippi Medical Center; Barrett Barnes, Frank Dipaola, Craig McKinney, Jeremy Middleton, Sean Moore, Eunice Odianse – University of Virginia.

## Author Contributions

JSC, SG, AH, MN, JOM, SR, JAS, PS, SBS, SS, JRT, LFW and FZ developed the concept and/or designed experiments.

MDA, CA, CB, DEB, HHB, JDB, NNB, PRB, XC, JSC, NSD, ARG, LPG, SCG, MG-J, LG-S, AH, JI, ZJ, CRJ, JL, CAM, NM, EN, FN, MN, JO-M, AP, KR, SFR, SR, SSR, DS, JAS, KS, LS, SS, ES-D, JRT, AU, TGW, MW, LFW, DZ and FZ acquired, analyzed and/or interpreted data. MDA, DEB, AJ, JL, JS and JRT were involved in the creation of new software used in this work.

HHB, JSC, JL, NM, FN, MN, JOM, DS, JAS, SS, JRT and LFW have drafted or substantially revised the manuscript.

## Competing Interest Declaration

The authors declare the following non-financial competing interests:

SS is a Board of Trustee member for Vital Pakistan and has a courtesy faculty appointment at the Aga Khan University.

The authors declare the following personal financial interests:

CRJ has served as an expert witness in pediatric litigation concerning osmotic demyelination syndrome and thiamine deficiency, unrelated to the subject of this work.

FN has consulted for PathAI.

JOM has consulted for Tessel Biosciences, Radera Biotherapeutics, and Passkey Therapeutics.

JAS has consulted for Alimentiv, Chugai Pharmaceutical Co. LTD, Mozart Therapeutics, Takeda Pharmaceuticals, Teva Pharmaceuticals, and Topas Therapeutics.

SS is co-Chair of IBD Plexus selection committee for the Crohn’s and Colitis Foundation (paid position).

MDA and JRT are named inventors on a patent application (PCT/US24/31703) related to the SPECTRE-Plex methods described in this manuscript.

JL, KR, JS and JRT are named inventors on provisional patents (US63/622,894, 63/622,925) related to software described in this manuscript.

## Additional Information

Supplemental Information is available for this paper.

Correspondence and requests for materials should be addressed to Jay R Thiagarajah at.

---

**Extended data 1:**
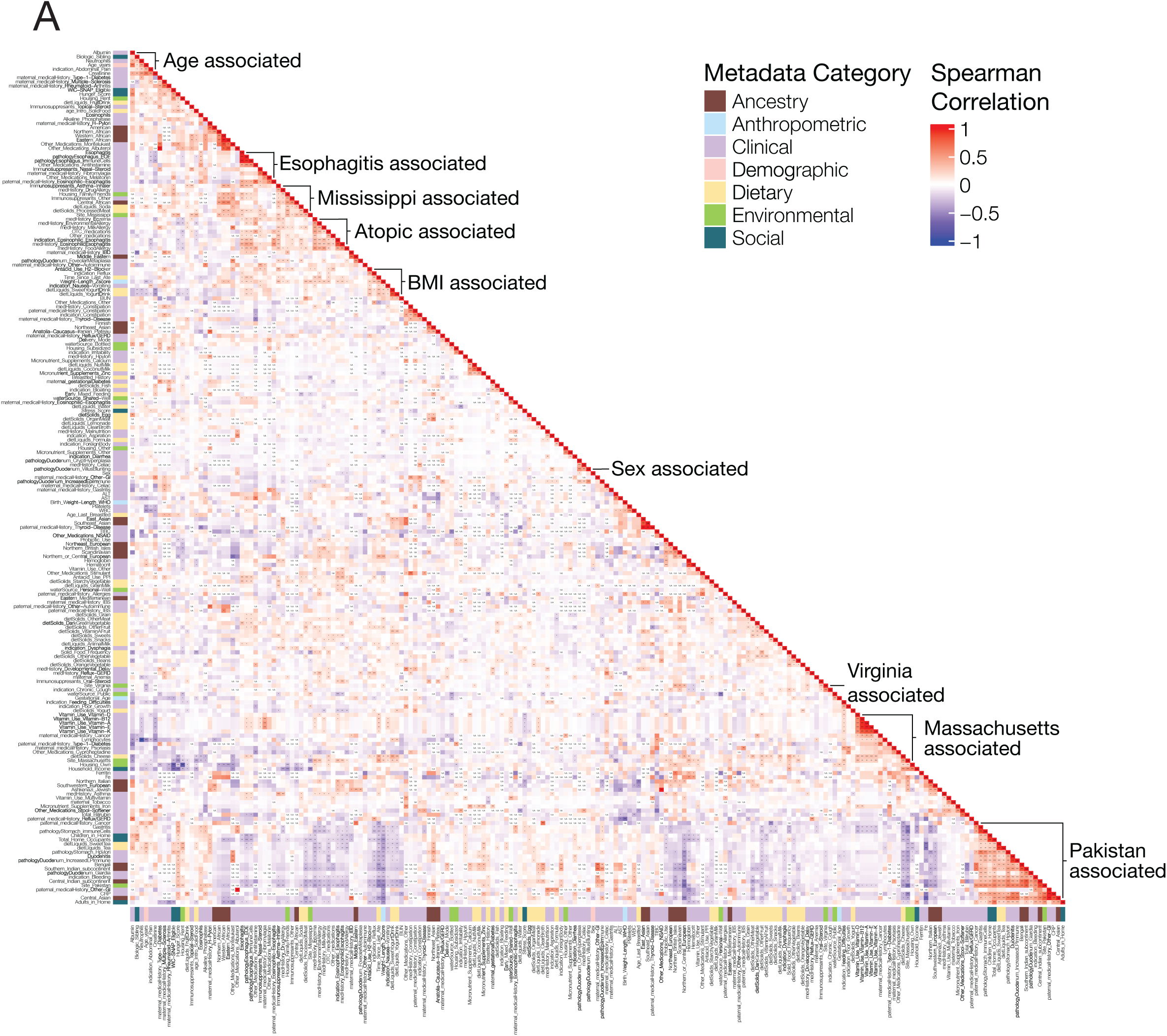
Metadata associations across Gut-AGE. **A.** Correlation matrix of all gut-AGE metadata, colored by data category and Spearman correlation coefficient. Correlated metadata was organized by performing hierarchical clustering and correlated modules were marked. NA denotes comparisons that do not have >3 pairwise comparisons. * p < 0.05, ** p < 0.01, ***p < 0.001, **** p < 0.0001.

**Extended data 2:**
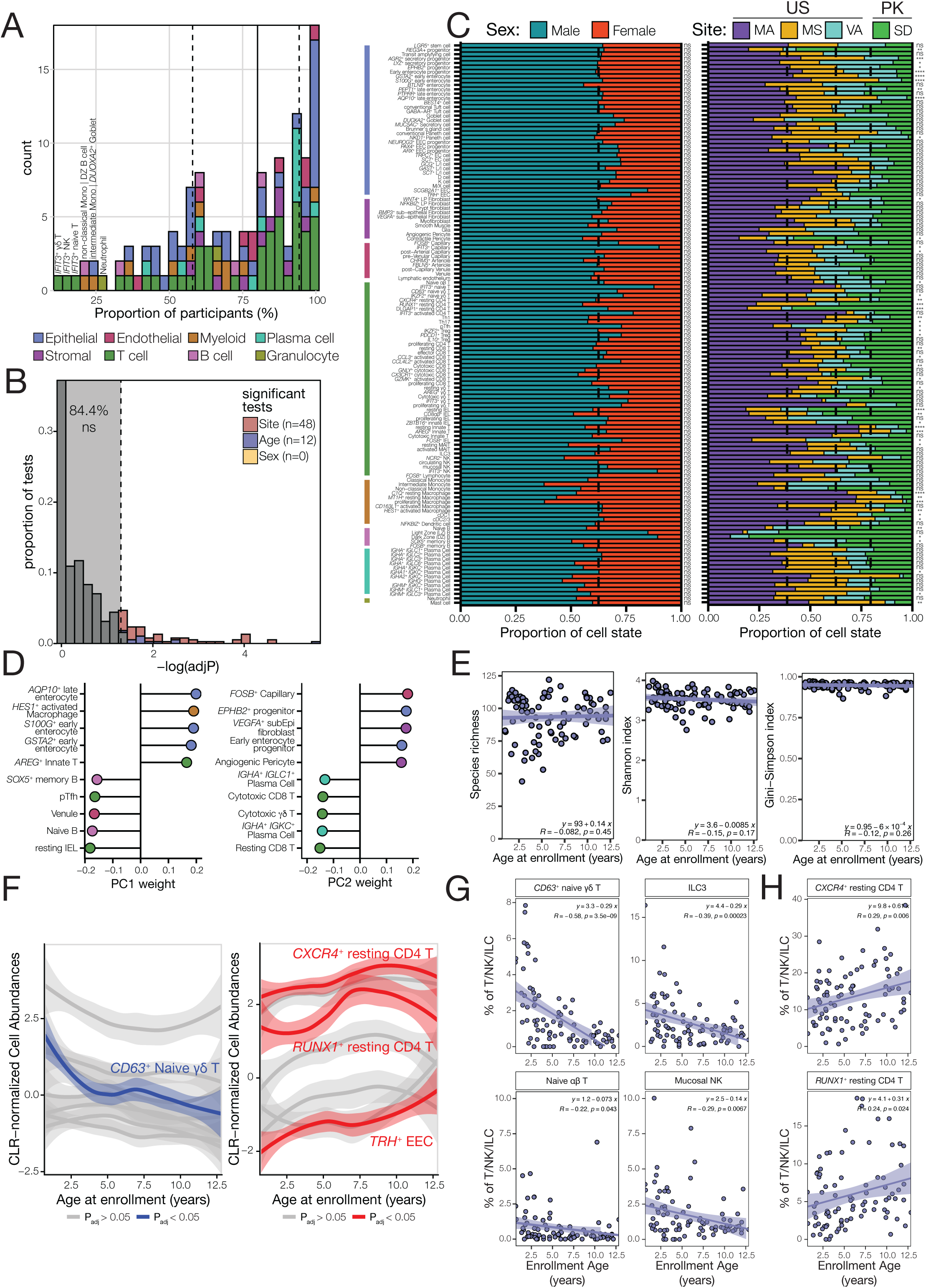
Most duodenal cell states are conserved across age and environment. **A.** Histogram of cell state representation across participants. For each cell state, the proportion of individuals with at least one cell detected was calculated as a product of the total number of participants. IQR (58.0-94.3%) is indicated by dotted lines and median (80.1%) is indicated by a solid line. Cell states are colored by broad cell identity and outlier populations are labelled. **B.** Proportion of cell states that were significantly associated with either site (red), age (blue) or sex (yellow). The dotted line denotes adjusted p value = 0.05. Cell abundance association with sex and site was statistically tested using a Wilcoxon rank-sum test. Cell abundance association with age was statistically tested using a linear regression. P values were adjusted using the Benjamini-Hochberg (BH) method. **C.** Stacked bar chart of the relative proportion of cell states across sex (left) and geography (right). The dotted line represents the overall patient distribution for each variable. The colored bars to the left correspond to broad cell identity. Significance is indicated to the right of each bar. **D.** Top 5 positive and top 5 negative cell abundance PC loadings for PC1 (left) and PC2 (right), colored by broad cell identity. **E.** Cellular diversity of participant biopsies across age. Species richness (left), Shannon index (middle) and Gini-Simpson index (right) were used to quantify biopsy diversity, with each cell state representing a unique species. Spearman rank correlation was used to calculate correlation coefficient. **F.** Proportional abundance plot of cell states either negatively (left) or positively (right) associated with age. Cell states with colored labels remain significant following p value adjustment (BH). **G.** Frequency plots of immune cell populations that are negatively associated with age as a proportion of total T/NK/ILC numbers. **H.** Frequency plot of T cell populations positively associated with age as a proportion of total T/NK/ILC numbers. ns p > 0.05, * p < 0.05, ** p < 0.01, ***p < 0.001, **** p < 0.0001.

**Extended data 3:**
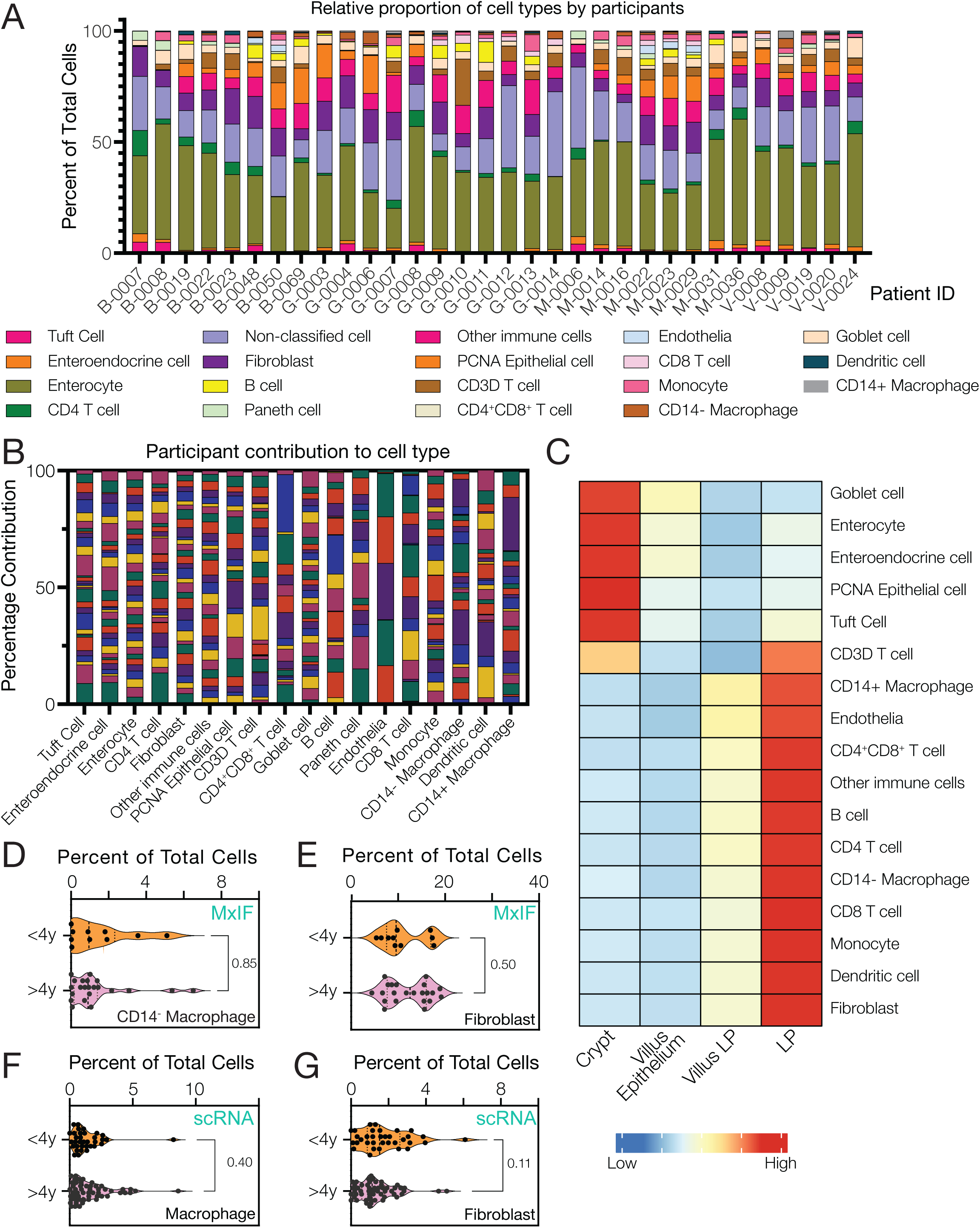
Multiplex immunofluorescence enables detection of distinct cell types across different morphological regions. **A.** Relative percentage of each cell type observed in the biopsy of every participant using MxIF. Each bar represents a single participant and each color a specific cell type. **B.** Plot showing the relative percentage of each cell type in every participant. Each bar represents a cell type, and the color represents the relative abundance of the cell type in each participant. **C.** Heatmap shows relative cell abundance of cell types in different morphological tissue structures within the duodenum. **D-G.** Percentage abundance of macrophages identified by MxIF (CD68^+^ CD14^-^) **(D)** or scRNA-seq **(F)** and fibroblasts identified by MxIF **(E)** or scRNA-seq **(G)** in young children (age < 4 years) and in older children (age > 4years) relative to all classified cells. Each dot represents a single participant

**Extended data 4:**
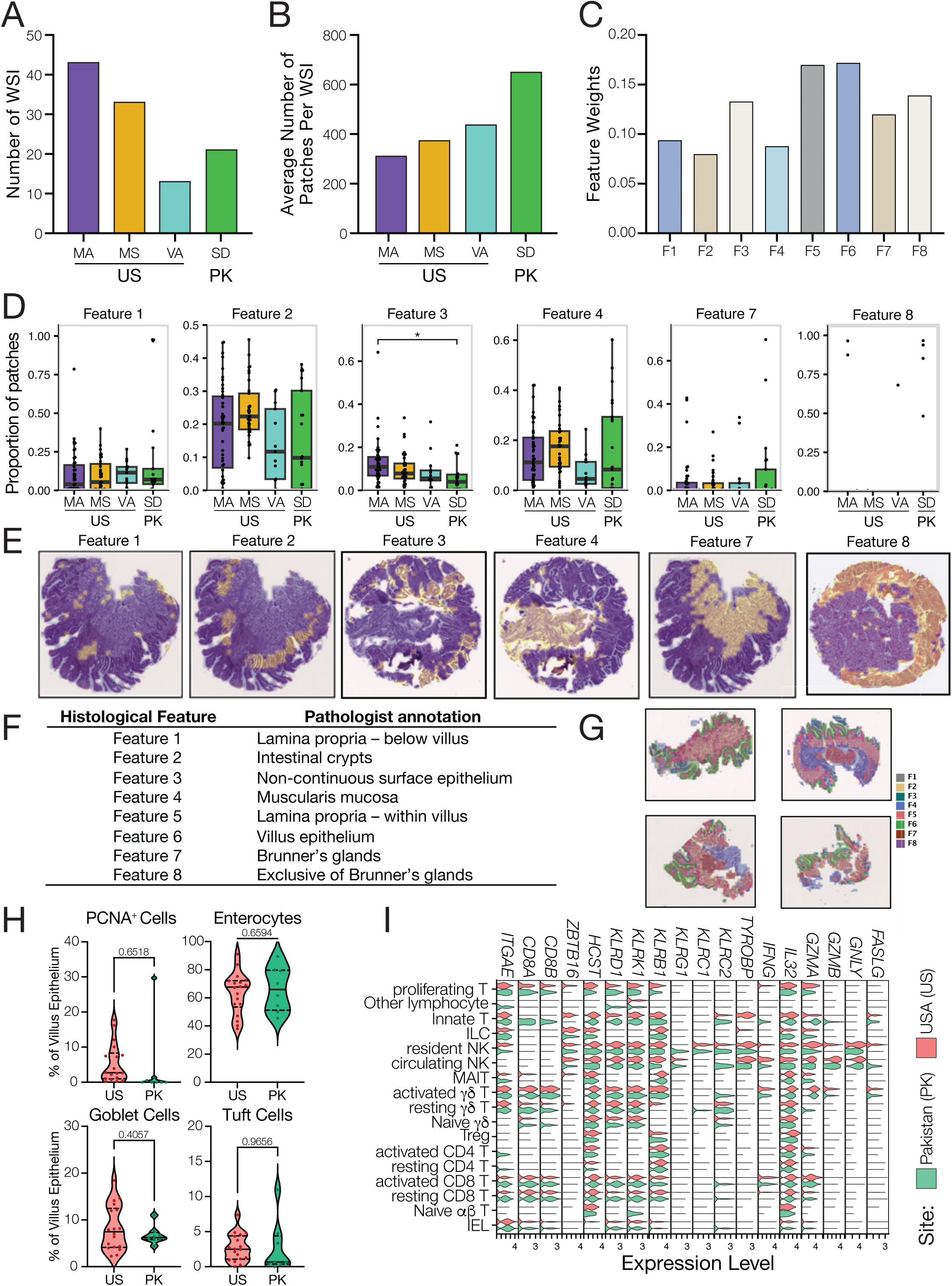
Features identified by patch-level feature extraction. **A.** Total number of whole slide images (WSI) processed for each site (one image per participant). **B.** Number of patches per WSI for each site. **C.** Feature importance in identifying sample site as determined by a random forest classifier entrained using Gut-AGE patch features. **D.** Comparison of proportional patch assignment across site. For each biopsy, feature proportion was calculated by dividing the number of patches assigned a specific feature by the total number of patches identified. Color corresponds to geographic sites **E.** Representative whole slide images of biopsies colored by regions associated with each annotated feature. **F.** Consensus annotations for each feature as determined by three independent pathologists. **G.** Representative biopsies segmented into patches and colored by feature. **H**. Frequency of various epithelial populations identified by multiplexed immunofluorescence by country of enrollment. **I.** Violin plot displaying the gene expression of IEL markers and features associated with IEL function across all T cell subsets. Color denotes country of enrollment. * p < 0.05, ** p < 0.01, ***p < 0.001, **** p < 0.0001.

**Extended data 5:**
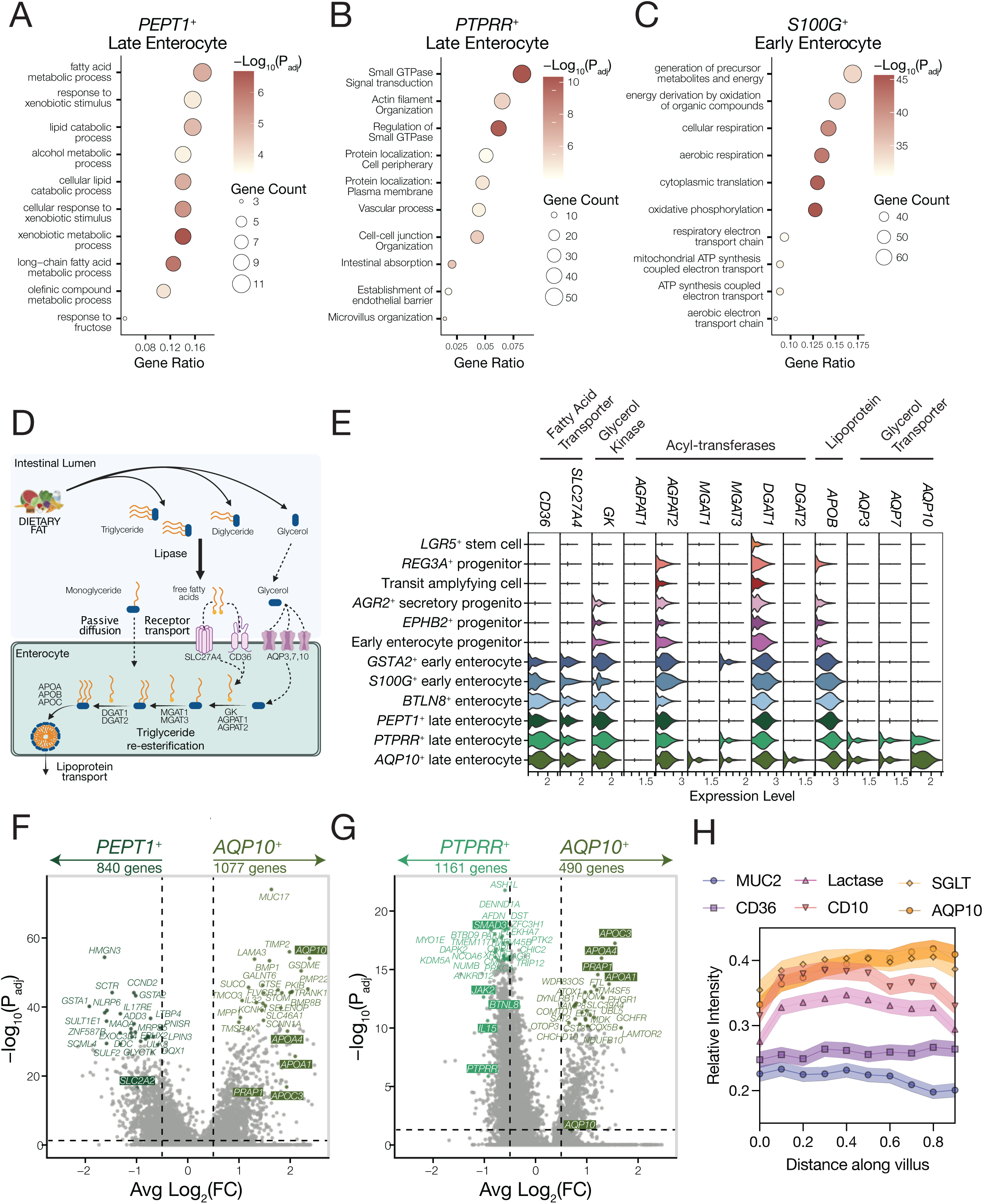
Late enterocyte states are functionally distinct. **A – C.** Top 10 GO terms enriched in genes significantly differentially expressed in major enterocyte subsets relative to all other epithelial states. P values were corrected for FDR using the BH method. **D**. Overview of the mechanism of dietary fat absorption from the lumen of the duodenum across enterocytes. **E.** Violin plot depicting expression of enzymes necessary for lipid absorption and digestion across enterocyte states. Color corresponds to enterocyte state. **F-G**. Volcano plot of genes differentially expressed between AQP10^+^ enterocytes and either PEPT1^+^ late enterocytes **(F)** or PTPRR^+^ late enterocytes **(G)**. A full list of differentially expressed genes can be found in **Supplementary table 8** (AQP10^+^ vs PEPT1^+^) and **Supplementary table 9** (AQP10^+^ vs PTPRR^+^). Differential gene expression was calculated using DEseq2 on samples pseudobulked by participant and cell identity. **H.** Quantification of fluorescence intensity of various epithelial markers across regions of the intestinal villus normalized to total villus intensity.

**Extended data 6:**
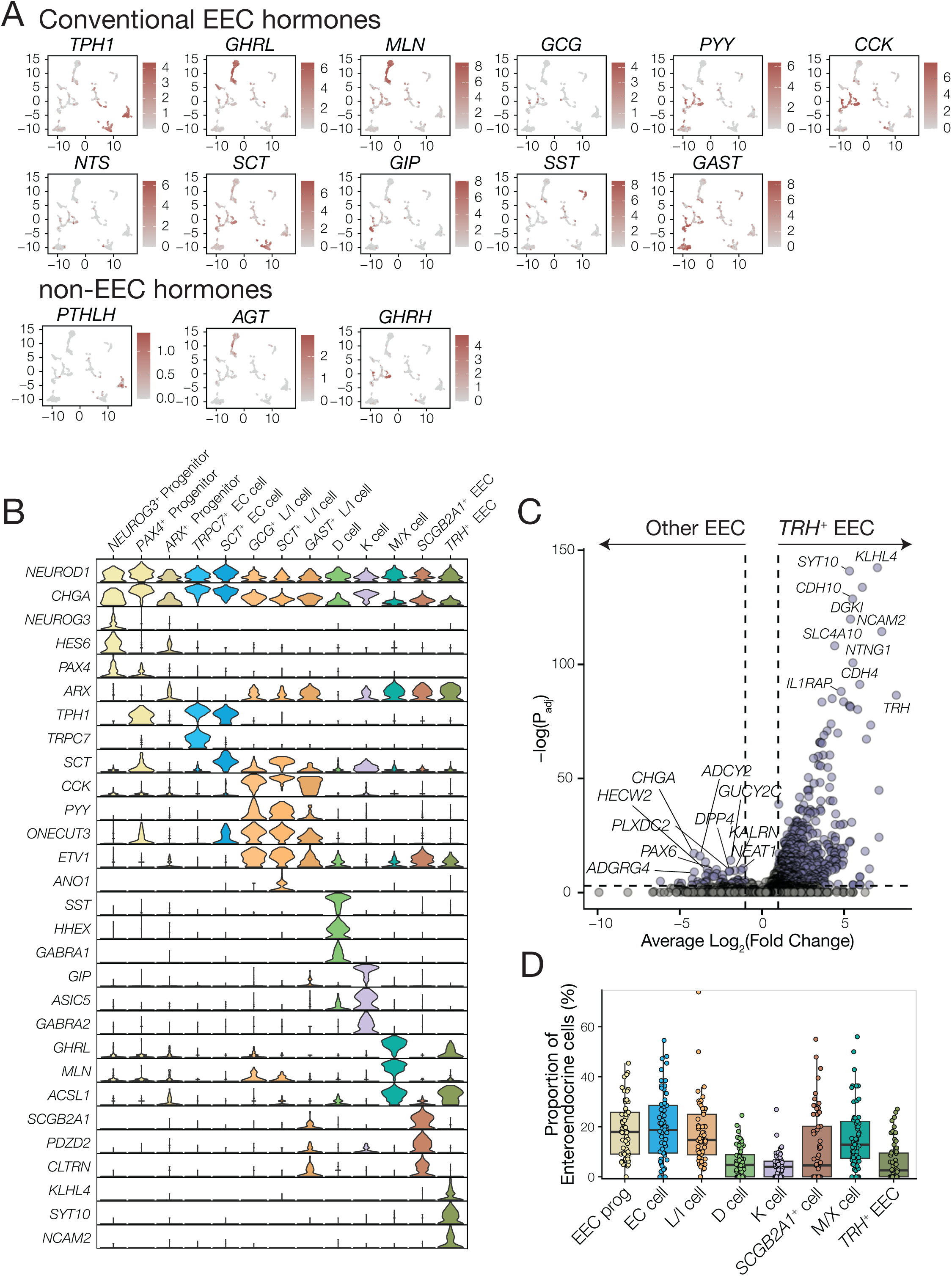
Enteroendocrine cell diversity within the pediatric duodenum. **A.** Expression of hormones previously associated with EEC function (conventional EEC hormones), as well as hormones not classically associated with EEC function (non-EEC hormones) overlaid on the UMAP representation of enteroendocrine cells **B.** Violin plot of top marker genes that define enteroendocrine subsets and states. Markers were identified from the top differentially expressed genes for each subset and state when compared to all other enteroendocrine cells. Color corresponds to enteroendocrine cell state. **C.** Volcano plot of genes upregulated in TRH^+^ enteroendocrine cells relative to all other enteroendocrine cells. The 10 genes with the lowest adjusted p value are highlighted. Differential gene expression was calculated using the Seurat FindMarkers function. A full list of differentially expressed genes can be found in **Supplementary table 10**. **D.** Frequency of enteroendocrine subtypes within the early childhood duodenum. EEC, enteroendocrine cell; EC, enterochromaffin cell.

**Extended data 7:**
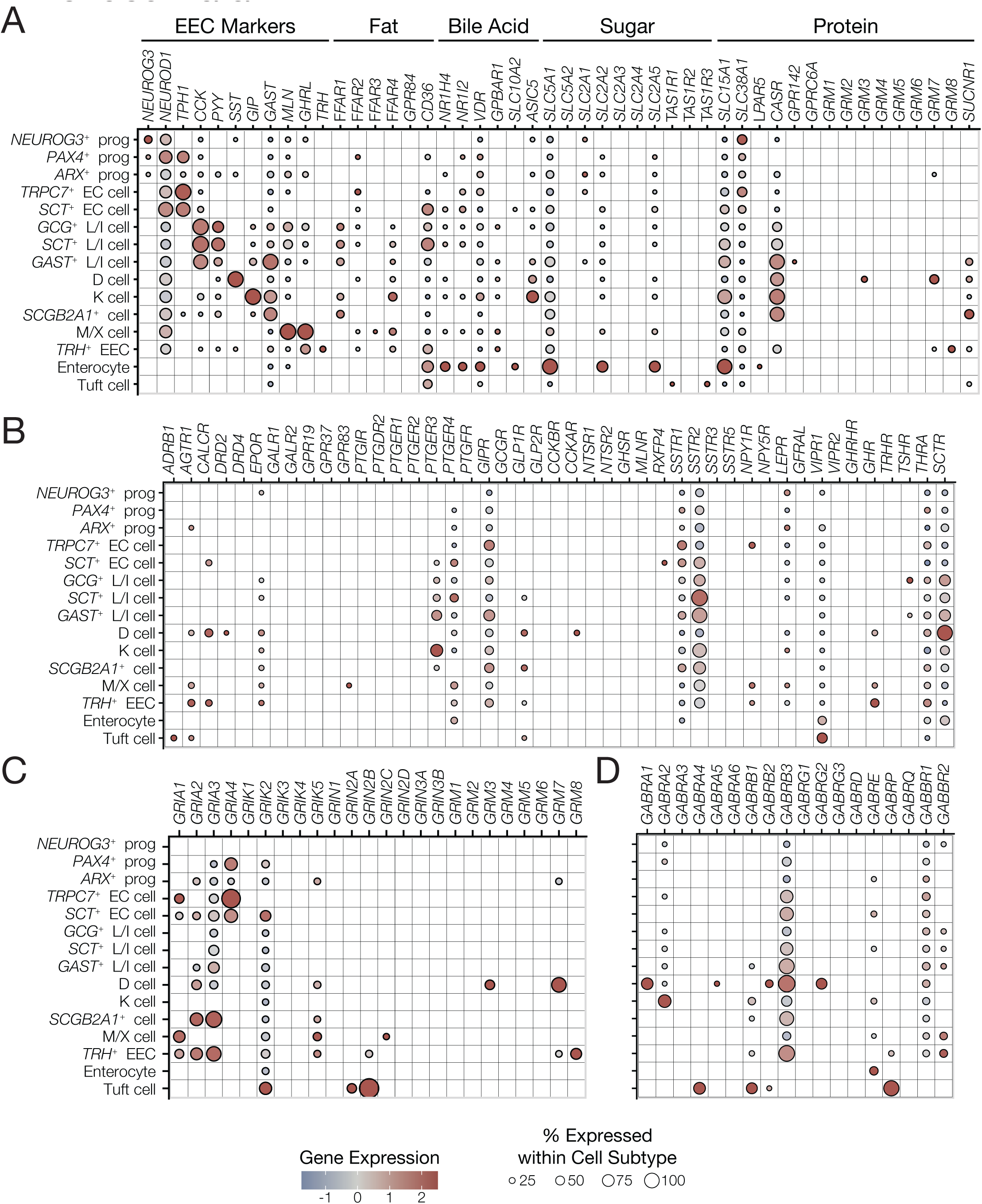
Receptor expression in duodenal enteroendocrine cells. **A.** Dot plot visualization of EEC markers and genes previously associated with the sensing of intestinal metabolites. **B**. Dot plot visualization of hormone receptor gene expression across EEC states. **C and D**. Dot plot visualization of glutamate receptors **(C)** and GABA receptors **(D)** gene expression across EEC states. Color corresponds to column-scaled z-score, and size corresponds to the proportion of cells positive for that gene. Prog, progenitor; EEC, enteroendocrine cell; EC, enterochromaffin cell.

**Extended data 8:**
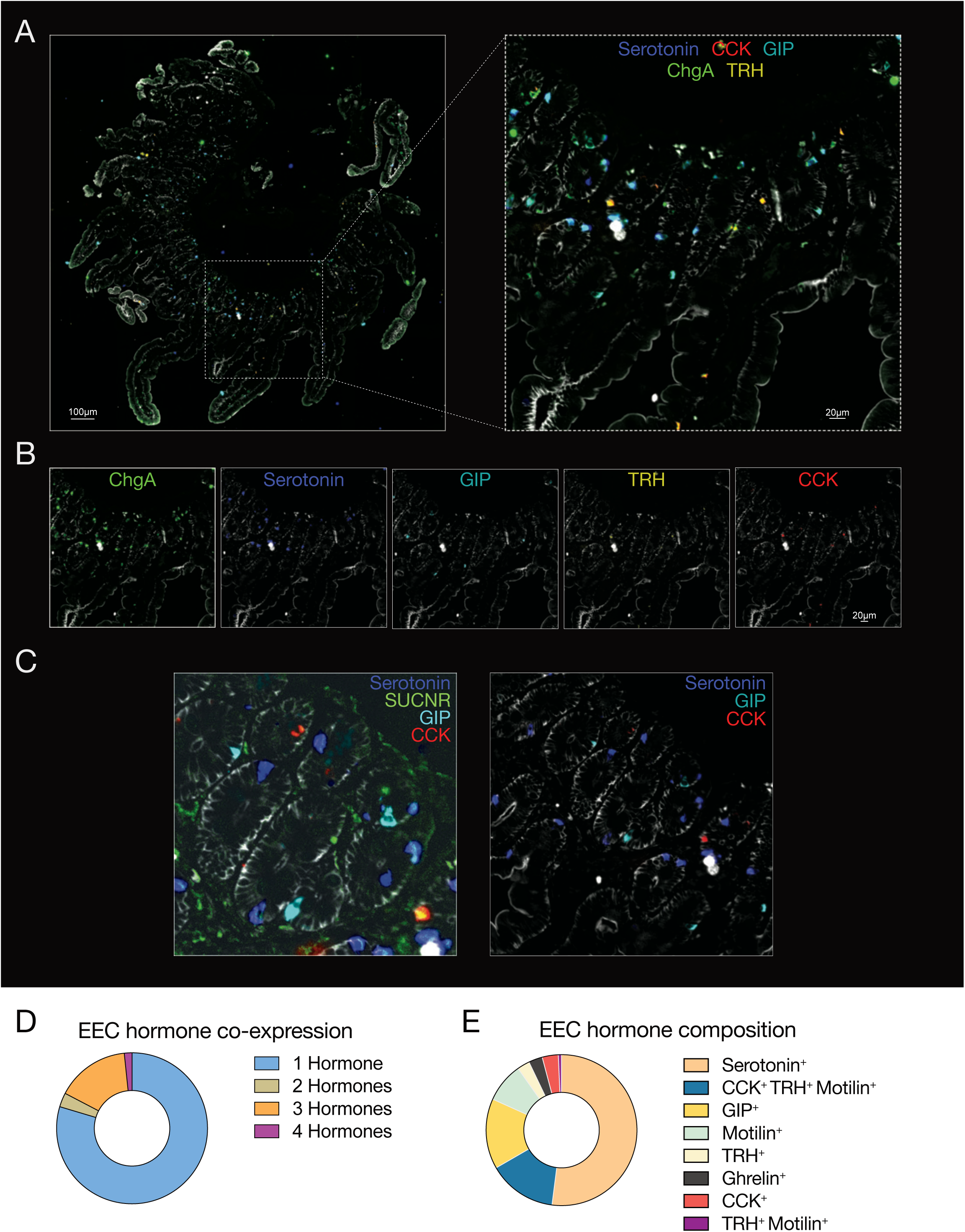
Spatial geography of enteroendocrine cells in duodenum. **A.** Representative MxIF image of a biopsy from the United States (Boston, Male, 3.0 years-old) with an expanded view stained with the following markers, Serotonin (violet), CCK (red), GIP (blue), ChgA (green) and TRH (yellow). **B.** Single channel decomposition of the expanded view from above showing the specific spatial localization of Serotonin (violet), CCK (red), GIP (blue), ChgA (green) and TRH (yellow). **C.** Expanded view of the same biopsy now displaying (left) Serotonin (violet), SUCNR (green), GIP (blue) and CCK (red) (right) Serotonin (violet), GIP (blue) and CCK (red). **D.** Pie chart showing the hormonal complexity of different EEC cells. Each slice represents the number of the hormones expressed in a cell while the size represents the relative abundance of EECs represented by the count. **E.** Pie chart showing the diversity of the hormones expressed by EECs. Each slice represents the hormones expressed while the size represents the relative abundance of this hormone in EECs.

## 1 Aims of the Glossary

The purpose of this glossary is to provide a clear and transparent rationale for the naming of each cell type, subtype and state in our dataset. Here we define a cell type as a broad, stable and developmentally distinct identity. An example of cell type would be an epithelial cell or a T cell. We divide these broad cell types into subtypes, which represent distinct and stable identities that are well supported within the literature. Subtypes possess agreed upon markers and functions. Finally, a cell state represents dynamic and potentially reversible response among cell subtypes. Cell states are denoted by their most informative marker gene, followed by their cell subtype designation. We also provide a brief overview on the population, any existing literature as well as marker genes (We have used the terms markers and genes interchangeably) that were informative in our annotations. We emphasize that all our definitions arise from iterative clustering of the transcriptomic data. Although we describe specific cellular nomenclatures, the underlying data fundamentally reflect mathematically derived clusters of individual cells. All subtypes and states are interpreted within this mathematical framework.

## 2 Stromal Cells

Stromal cells are a broad catchall for cell types that do not fit into epithelial, endothelial, nor immune categories. As such, they are *PTPRC*, *PECAM1*, and *EPCAM* negative. Subsets are fibroblasts (marked by the expression of *PDGFRA*), Pericytes, Glia and smooth muscle.

Quality control metrics (200 < nFeature_RNA *<* 4000 and percent.mt *<* 40%)

**Figure 1:**
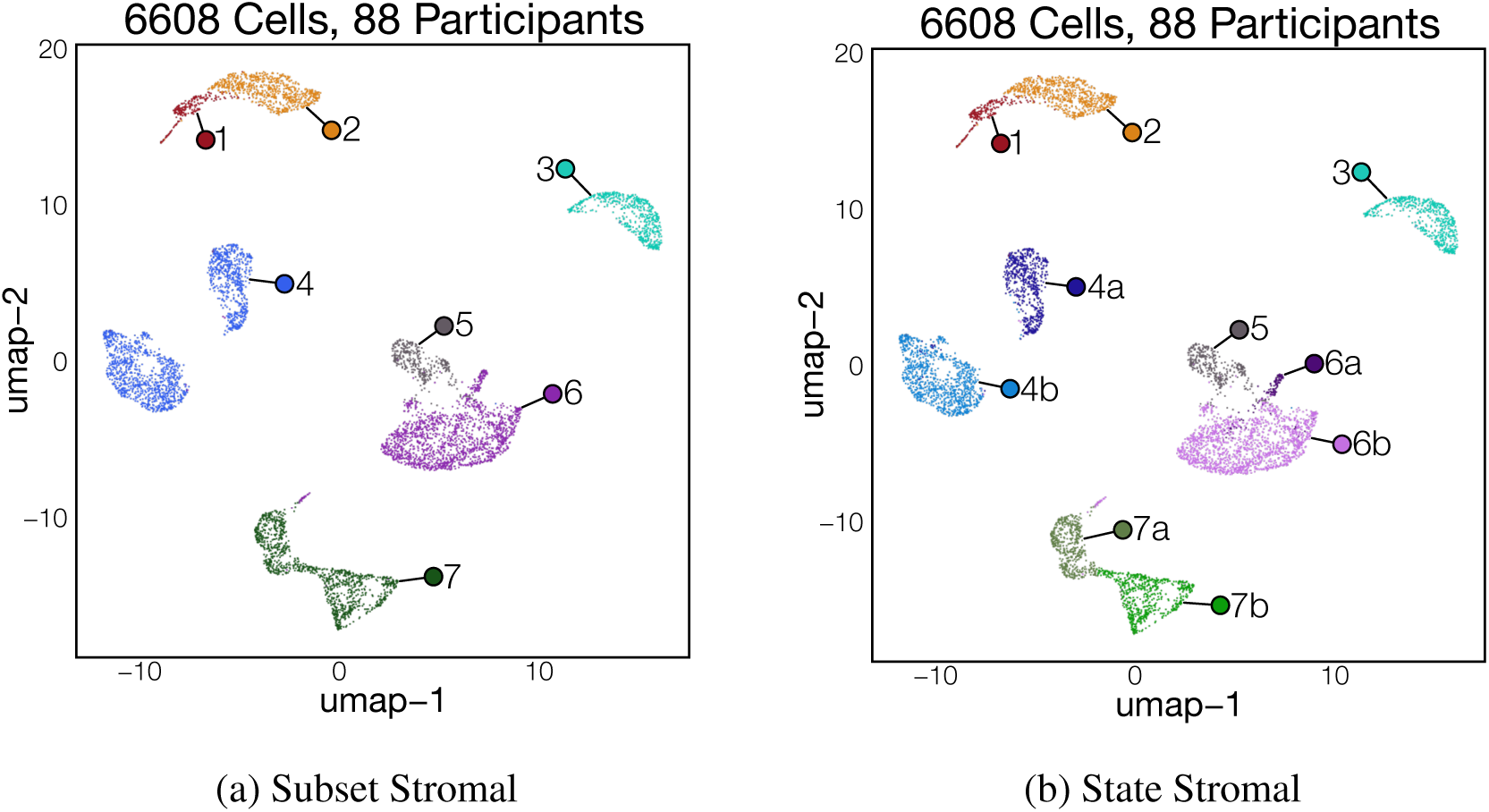
Stromal Cells

**1. Smooth Muscle:**

Cells in this cluster expresses high levels of *ACTA2*, *MYH11*, and *DES*, markers previously associated with both smooth muscle and myofibroblasts.[1, 2] Unlike myofibroblasts, smooth muscle cells express both *GREM1* and *GREM2*.[3] As biopsies do not capture full intestinal thickness, these cells are likely from the muscularis mucosa

**2. Myofibroblast:**

Like smooth muscle, myofibroblasts express high levels of *ACTA2*, *MYH11* and *DES*. They also express *FOXF1* and *HHIP*, but not *PDGFRA*.

**3. Glia:**

Enteric glia are stromal cells that actively support enteric neurons. They exist distributed with nerve fibers in the muscle as well as within the mucosa.[4] Enteric glia express *PLP1*, *S100B*, *CDH19* and *CRYAB*.[5] A small proportion of glia express *SOX10*, which may indicate glial progenitors.

**4. Pericyte:**

Pericytes are broadly defined by the expression of *PDGFRB* and *NOTCH3* and function in close association with endothelial cells.[6, 7]

**a. Contractile Pericytes:** These cells express high levels of *ACTA2* and *MYH11*, and uniquely express *MUSTN1*. Expression of these transcripts have been described in contractile pericytes that line arterioles.[8] Previous work describes contractile pericytes in humans and in mice.[2, 9, 10]
**b. Angiogenic Pericytes:** Defined by expression of *EBF1*, *ADGRF5* and *RGS5*. *RGS5* has been associated with angiogenic activity in pericytes.[11]

**5. Crypt fibroblast:**

Crypt firbroblasts express *PDGFRA* at low levels, and co-express *RSPO3* and *GREM1*. Prior work in the murine small intestine showed that these cells are proximal to intestinal crypts[12, 13] and are necessary for intestinal stem cell maintenance. Also referred to as S3 cells and Trophocytes.[1]

**6. Lamina propria Fibroblast:**

Both states of LP fibroblasts express *PDGFRA* albeit at lower levels than sub-epithelial fibroblasts. They also express *ADAMDEC1* and *LUM*. Prior work has shown broad distribution within the lamina propria. Also referred to as S1 fibroblasts.[1, 12]

**a. NFKBIZ**^+^ **LP Fibroblast:** High expression of inflammatory and early response genes.

Most express *FOS*, *JUN*, *NKFBIZ* and *SOCS3* which is suggestive of an activated state. Notably, this population is rare and was not present in all biopsies

**a. b. WNT4**^+^ **LP Fibroblasts:** Cells in this state comprise the majority of LP fibroblasts and

were the only cells that expressed *WNT4* in our dataset. Prior studies have shown that *WNT4* expressing fibroblasts are distributed within the lamina propria above intestinal crypts.[12]

**7. Sub-epithelial Fibroblast:**

Cells annotated as this subtype express high levels of *PDGFRA* in addition to *F3*, *NRG1* and *SOX6*. Furthermore, these cells express the majority of BMPs, including *BMP2*, *BMP3*, *BMP4*, *BMP5*, *BMP6* and *BMP7*. Prior work has shown these cells are near the epithelial lining and are necessary for differentiation of the epithelium.[13] Previously referred to as S2, Telocytes, and WNT5B^+^ Fibroblasts.[12, 13, 14, 1]

**a. BMP3**^+^ **sub-epithelial Fibroblast:** In addition to being the primary source of *BMP3*, cells in this state express *VSTM2A*, *PCSK6* and *TLL2*.
**b. VEGFA**^+^ **sub-epithelial Fibroblast:** Cells in this state express *VEGFA*, which has previously been associated with fibroblasts localized to the villus tip[15]. They also express *TRPA1* and *NPY*.

## 3 Endothelial cells

This broad category encompasses all cells expressing the endothelial marker *PECAM1* (*CD31*) which are negative for other lineage markers such as *PTPRC* and *EPCAM*. It includes endothelial cells that make up the vasculature and lymphatics of the duodenum.

Quality control metrics (200 *<* nFeature_RNA *<* 4000 and percent.mt *<* 40%)

**Figure 2:**
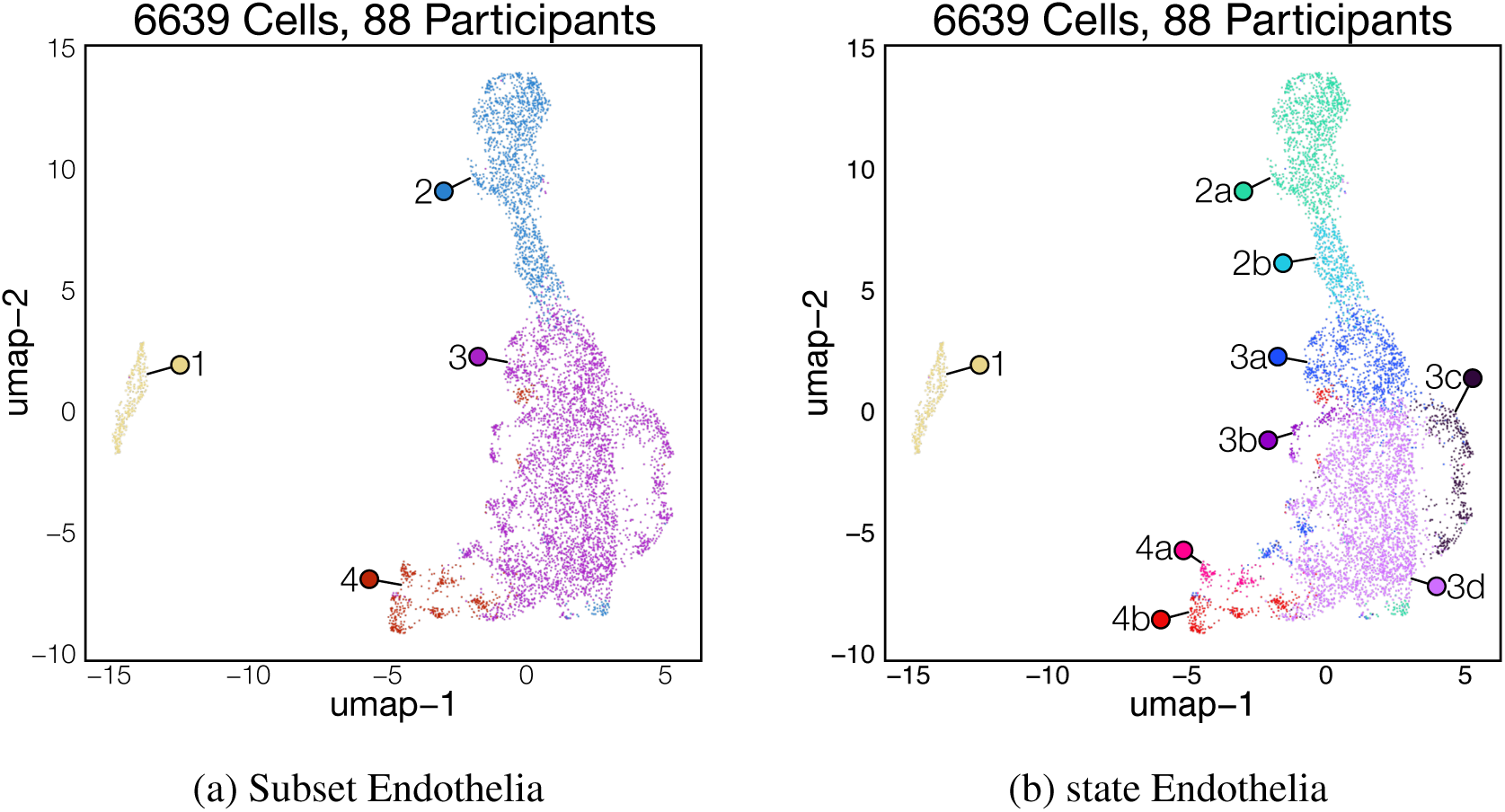
Endothelia

**1. Lymphatic endothelium:**

Cells within this cluster express high levels of *LYVE1*, *CCL1*, and *PROX1*, markers associated with the lymphatic endothelium.[16, 17] Within the duodenum, lymphatics assist in fat digestion by absorbing chylomicrons.[18]

**2. Venular endothelium:**

Venular endothelia express high levels of *ACKR1*, *MADCAM1*, and various selectins including *SELP* and *SELE*. This is consistent with prior literature describing venules as the primary site of immune cell interaction and entry.[19]

**a. Venule:** These cells have elevated levels of *ADGRG6*, *IL1R1* and *POSTN*. Within other tissues, expression of these genes increases with transition from venules to large veins, indicating that these cells may form larger vessels than other venule states.[20]
**b. Post-Capillary Venule:** In this state, expression of venule marker genes is lower relative to venule state. In addition, in the UMAP space this cluster is more proximal to the cell cluster annotated as capillary which indicate it may be a post-capillary venule. These cells also express *CCL14* and *CLU*.

**3. Capillary:**

All capillary cells expressed *CA4*, *RGCC* and *INSR* which are known to be expressed primarily in capillaries found throughout the body.[21, 16]

**a. Pre-Venular Capillary:** Cells in this state express capillary makers but are closer to venules along a calculated trajectory.[20] They also express *RBP7* and *AQP1*.
**b. IFIT3**^+^ **Capillary:** Cells in this state express *IFIT3* and other interferon stimulated genes (ISG), including *ISG15* and *IDO1*. These cells were present in a small subset of individuals and may be indicative of an active response to enteric infection.
**c. FOSB**^+^ **Capillary:** This state is defined by expression of *FOS*, *JUN* and *NKFBIZ* which indicates that these capillaries may be activated and/or stressed.
**d. Post-Arterial Capillary:** These cells express capillary makers but are closer to venules along a calculated trajectory.[20] They also express *IGFBP3*, *P2RY8* and *LNX1*.

**4. Arteriole:**

Arterioles broadly expressed *VEGFC* and *SEMA3G* which were positively associated with arteries and arterioles in prior atlases.[20, 16]

**a. CHRM3**^+^ **Arteriole:** Arteriole state defined by expression of *CHRM3*, a muscarinic acetylcholine receptor. In murine studies, *M3R* was shown to be necessary for artery dilation.[22] Cells in this state also express *ADAMTS6* and *SSUH2*.
**b. FBLN5**^+^ **Arteriole:** Defined by high expression of *FBLN5* and *SULF1*. Previously, *SULF1* was associated with larger arteries and arterioles.[16] This state may represent larger vessels relative to the other arteriole state.

## 4 B Cells

Cells within this broad categorization express the broad immune marker *PTPRC* (*CD45*) as well as *CD19* and *MS4A1* (*CD20*) which are canonical B cell markers in flow cytometry. B cells also express *CD79A*, *CD79B*, and *PAX5*. [23, 24, 25] Notably, almost all B cells in our dataset express *CD69*, which is likely necessary for their retention within the intestine.[26] There is also a high degree of *VPREB3* expression among all B cells, a surrogate light chain involved in BCR selection during B cell development. Although B cell development has been observed in the intestine, we do not detect any expression of *RAG1* or *RAG2*.[27]

Quality control metrics (200 *<* nFeature_RNA *<* 5000 and percent.mt *<* 25%)

**Figure 3:**
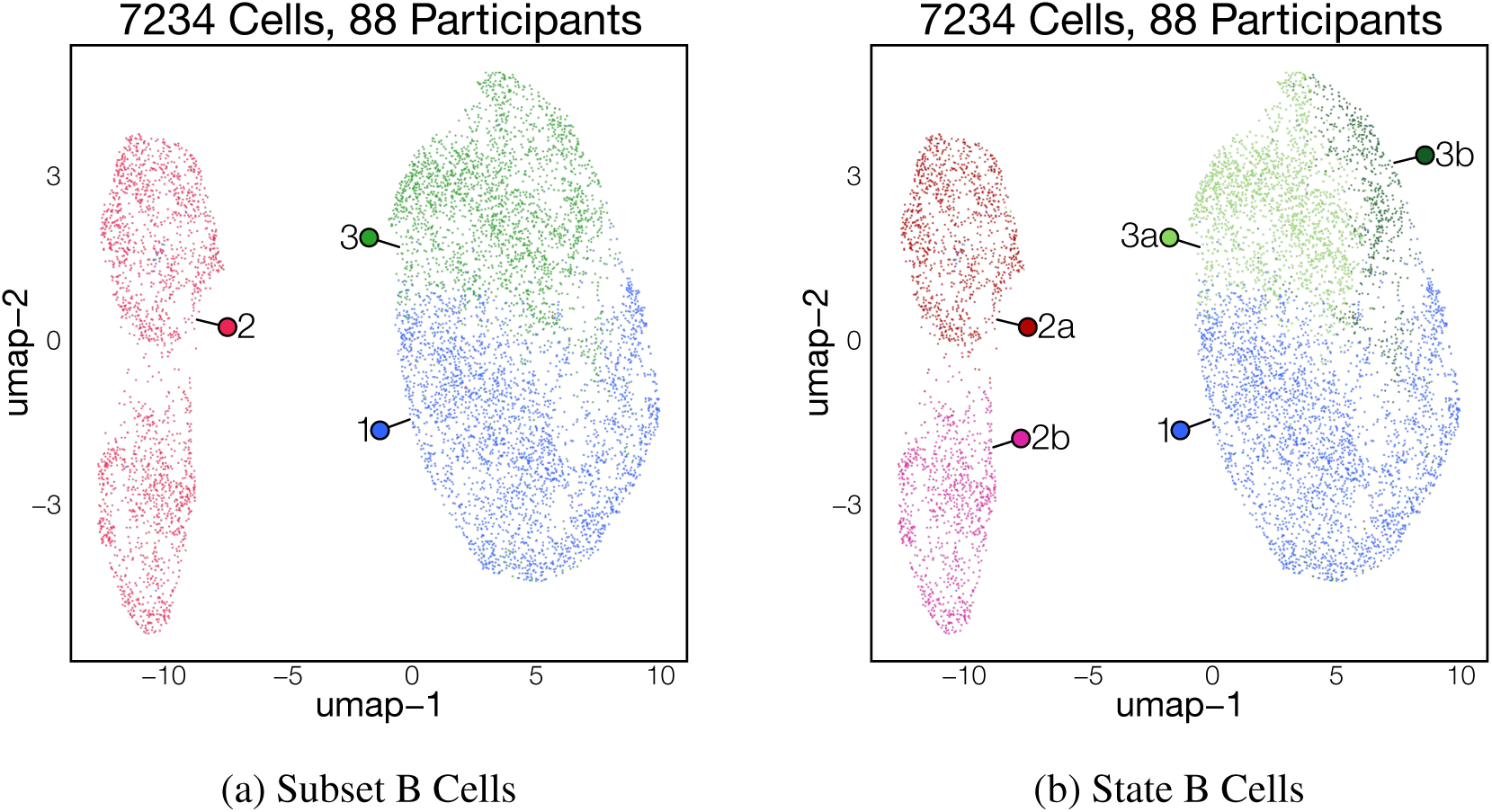
B Cells

**1. Naïve B:**

This subset has high expression of both *IGHD* and *IGHM* in the absence of any other BCR isotypes. Furthermore, there was expression of *FCER2* (*CD23*), *SELL* and a lack of *CD27* expression. These genes indicate a naïve B cell that has yet to recognize its cognate antigen.

**2. Germinal Center (GC) B:**

Germinal center B cells express *AICDA* (*AID*), a key mediator of somatic hypermutation, at varying levels depending on their location within the germinal center. Furthermore, there is evidence of *CD40* response with expression of *TRAF1* across cells. At this cluster level we find evidence of both light zone and dark zone GC B cells.

**a. Light Zone (LZ) B:** These cells express *AICDA*, albeit at lower levels than dark zone B cells. Light zone B cells are positive for *CD40* and *LMO2*.[28]
**b. Dark Zone (DZ) B:** In addition to expressing higher levels of *AICDA*, dark zone B cells have a strong proliferation signature enriched in *MKI67*.

**3. Memory B:**

Memory B cells have previously encountered their cognate antigen, so they have differentiated, undergone somatic hypermutation and possibly also class switch recombination. Memory B cells are defined by expression of CD27.[29] In our dataset, they also had high expression of *CD69* and *BCL2*.[30]

**a. FOSB**^+^ **memory B:** Cells in this state express memory B cell markers and are enriched for early response genes *FOSB*, *JUNB*, *NR4A1*, and many others. These genes are rapidly induced upon BCR stimulation,[31] suggesting that these cells are recently activated B cells. However, these genes can be induced by several signals, including stress during single cell dissociation.[32]
**b. SOX5**^+^ **memory B:** Cells in this state express high levels of *SOX5*, *RUNX2* and *FCRL4*.[33] FCRL4^+^ B cells have been described as a distinct memory population localized to mucosal tissues.[33, 34]

## 5 Plasma Cells

Plasma cells were defined by high expression of both heavy chain and light chain BCR genes. Most also express high levels of *JCHAIN*, which serves as a linker for polymeric IgA and IgM antibodies. Upon subsequent sub-clustering, plasma cells primarily separated based on the type of heavy and light chains used, so they were named based on this feature. Notably, they do not express *PTPRC* (*CD45*), which has been associated with long-lived tissue resident plasma cells.[35] Plasma cell states are predominantly differentiated by divergent heavy and light chain expression, likely reflecting antigen specificity and/or genetic differences in VDJ recombination rather than cell state differences.

Quality control metrics (200 *<* nFeature_RNA *<* 5000 and percent.mt *<* 20%)

**Figure 4:**
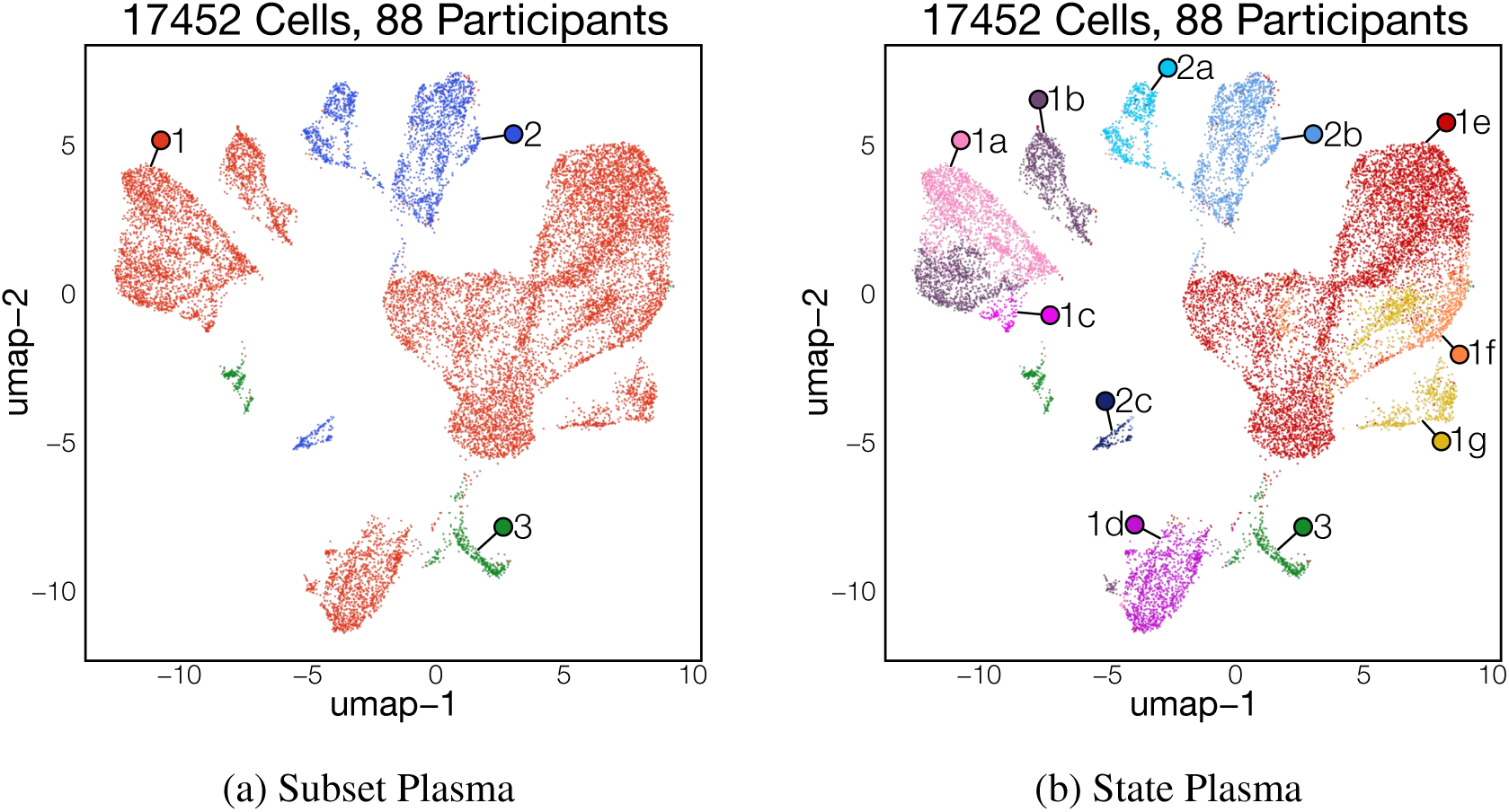
Plasma Cells

**1. IgA Plasma cell:**

IgA plasma cells had high levels of either *IGHA1* or *IGHA2*, *JCHAIN* and either lambda or kappa light chains. These plasma cells make up most plasma cells identified within the duodenum.

a. **IGHA**^+^ **IGLC2**^+^ **Plasma Cell**
b. **IGHA**^+^ **IGLC3**^+^ **Plasma Cell**
c. **IGHA**^+^ **IGLC6**^+^ **Plasma Cell**
d. **IGHA**^+^ **IGLC1**^+^ **Plasma Cell**
e. **IGHA1**^+^ **IGKC**^+^ **Plasma Cell**
f. **IGHA2**^+^ **IGKC**^+^ **Plasma Cell**
g. **IGHA**^+^ **IGKC**^+^ **Plasma Cell**

**2. IgM Plasma cell:**

Cells within this cluster expressed high levels of *IGHM* and *JCHAIN*. Notably these cells also express *CCL3* and *KLHL14*, which was also observed in bone marrow IgM^+^ plasma cells.[36]

a. **IGHM**^+^ **IGLC3**^+^ **Plasma Cell**
b. **IGHM**^+^ **IGKC**^+^ **Plasma Cell**
c. **IGHM**^+^ **IGLC1**^+^ **Plasma Cell**

**3. IgG Plasma cell:**

IgG Plasma cells express high levels of *IGHG*. Their relative rarity compared to other plasma cell subsets precluded higher resolution identification based on gamma chain usage. Consequently, this could be a mixed population of IgG^+^ plasma cells.

## 6 Granulocytes

Granulocytes were not initially identified as a separate population. They could only be identified by sub-clustering of unrelated populations (TNK ILC and Myeloid).

Quality control metrics (200 *<* nFeature_RNA *<* 4000 and percent.mt *<* 25%)

**Figure 5:**
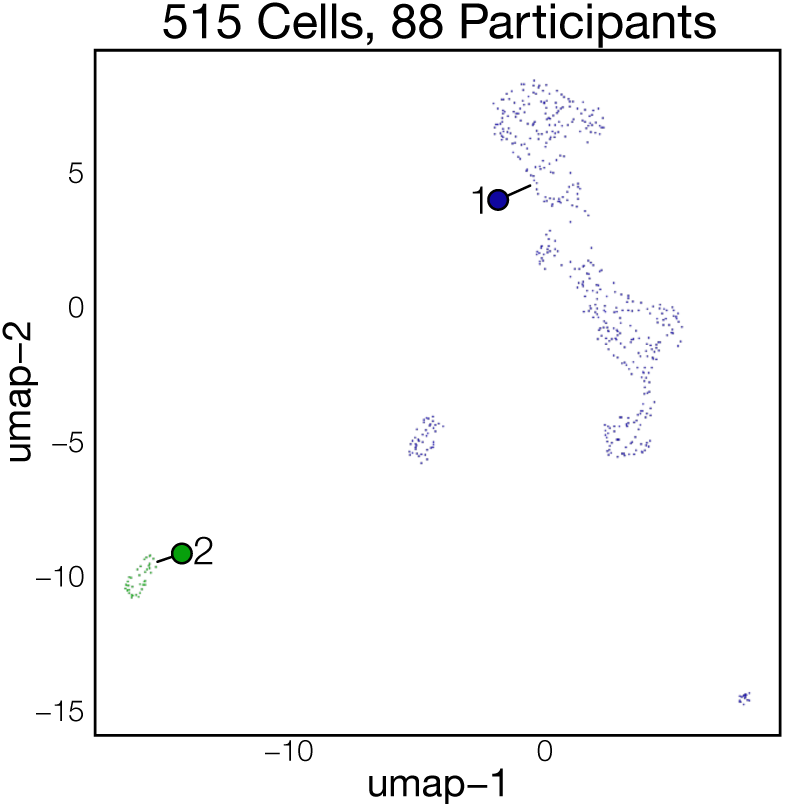
Granulocytes

**1. Mast cells:**

Although mast cells initially grouped with T/NK/ILC cells, sub-clustering revealed that their expression signature is distinct from lymphocytes. Mast cells are defined by high expression of mast cell granule genes such as *CPA3*, *TPSB2*, and *TPSAB1*. They also express high levels of the cell surface marker *KIT* and of *HDC*, the enzyme necessary to synthesize histamine.

**2. Neutrophils:**

Neutrophils express high levels of *CSF3R*, *CYBB*, *S100A8* and *S100A9*.[37] They were rare in the duodenum of our cohort of relatively healthy children and initially grouped with Myeloid cells.

## 7 Myeloid Cells

Myeloid cells are a diverse immune cell population (PTPRC^+^) comprised of macrophages, monocytes and dendritic cells. Myeloid cells are defined by high *ITGAM* expression. The majority also express *CD14*.

Quality control metrics (200 *<* nFeature_RNA *<* 5000 and percent.mt *<* 25%)

**Figure 6:**
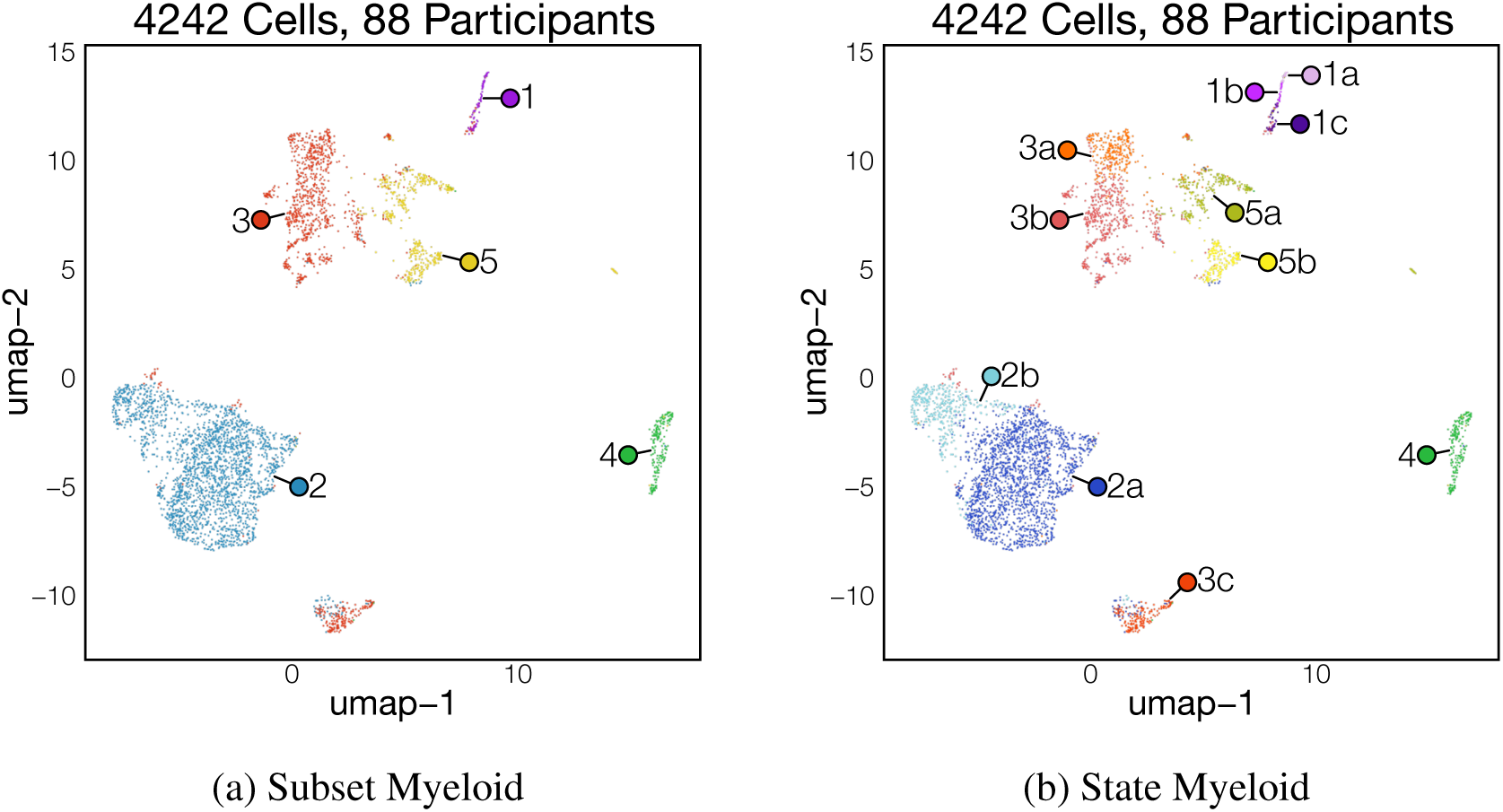
Myeloid Cells

**1. Monocyte**

Prior atlases have defined monocytes as myeloid cells with high expression of *S100A8*, *S100A9* and *VCAN*.[38] Using these markers, we identify 3 monocyte states, which we further define using existing monocyte nomenclature based on the presence of *CD14* and *FCGR3A* transcripts. As our cohort consists of uninflamed duodenum, monocytes are a rare population and are a minority of identified myeloid cells.

**a. Intermediate Monocyte** Intermediate monocytes co-express *CD14* and *FCGR3A*. We identify a cluster of cells that are positive for these markers, and have high expression of *IL21R*.[39, 40]
**b. Non-classical Monocyte** Non-classical monocytes have previously been identified by positive expression of *FCGR3A* (*CD16*).[41, 38] These cells expressed *FCGR3A* at higher levels than intermediate monocytes defined previously.
**c. Classical Monocyte** Classical monocytes have been defined in humans by the presence of *CD14* and absence of *CD16* (encoded by *FCGR3A*).[41] We use the expression of these markers to define classical monocytes. In addition, we note that in agreement with prior studies, cells within this cluster demonstrate higher expression of *CCR2* relative to other identified monocyte states.[42, 43]

**2. Resting Macrophages:**

Resting macrophages have high expression of *CSF1R*, *CD68*, *CD14* and evidence of MHC class II expression, with both *CD74* and *HLA-DRA*. Relative to other macrophage subsets, the resting population had lower RNA counts and were less diverse, which may indicate a relatively quiescent population.

**a. C1Q**^+^ **resting Macrophage:** Cells in this state express high levels of the C1 receptor complex, including *C1q*, *C1r* and *C1s*. Macrophages are the primary source of C1 within the intestine where complement components are produced at basal levels to protect against enteric infection. [44]
**b. MT1H**^+^ **resting Macrophage:** These cells broadly express metallothionein genes, including *MT1H*, *MT1G*, and *MT2A*. Macrophages high for metallothionein have previously been observed but their exact role in health is not well understood.[45]

**3. Activated Macrophages:**

Activated macrophages express traditional macrophage markers and have relatively high RNA counts which may indicate a more transcriptionally active state.

**a. CD163L1**^+^ **activated Macrophage:** *CD163L1* upregulation has been associated with both *m-CSF* and *IL10* responses in macrophages.[46, 47] *CD163L1* expression in macrophages has been associated with a tissue-resident anti-inflammatory population. Cells in this state also express *ITGA9* and *MERTK*.
**b. HES1**^+^ **activated Macrophage:** *HES1* has been used previously in multiple tissue types to identify a macrophage state with an M2-skewed phenotype.[48, 45] Cells in this state also express high levels of *CCL3/4*, *PRDM1* and *MRC1*.
**c. Proliferating Macrophage:** These macrophages with a strong proliferation signature. Top differentially expressed genes include *MKI67*, *TOP2A* and *PCLAF*.

**4. cDC1:**

These cells express dendritic cell markers *FLT3* and *ITGAX* as well as genes previously used to identify cDC1 cells, including *IDO1*, *XCR1*, *CLEC9A* and *BATF3*.[49] These cells are capable of cross-presenting foreign antigen to CD8 T cells.

**5. cDC2/3:**

cDC2/3 cells are a subset of conventional/myeloid derived dendritic cells that express *IRF4* and *CD1C* in addition to the broad dendritic cell markers *FLT3* and *ITGAX*.[50, 51] In our dataset, these cells also express *CD207*, a marker associated with skin dendritic cells, but also described within the intestine.[52, 53]

**a. NFKBIZ**^+^ **Dendritic cell** These cells express high levels of *NFKBIZ*, a suppressor of the NFKB pathway, upregulated following NFKB signaling.[54] This may indicate that these cells have recently been activated through an NFKB-dependent pathway. These cells express lower levels of *CD1C* relative to other cDC2/3 states.
**b. cDC2/3** These cells express transcripts previously associated with cDC2/3s, including *CD1C* and *FCER1A*.[38] Upon performing differential gene expression, there was minimal evidence of any additional activation signatures, which may indicate that these cells represents a resting population of cDC2/3.

## 8 Epithelial Cells

The epithelium makes up the majority of cell types captured within our dataset of the pediatric duodenum. Epithelial cells were defined here based on their expression of *EPCAM* in the absence of any markers specific to other lineages. Epithelial cells, especially enterocytes, typically had higher mitochondrial content relative to other cell types. In contrast, crypts tend to have higher ribosomal RNA content.[55, 56] Generous filtration criteria were applied to reduce the risk of removing epithelial cell subtypes.

Quality control metrics (200 *<* nFeature_RNA *<* 5000 and percent.mt *<* 60%)

### 8.1 Progenitors and absorptive epithelium

**Figure 7:**
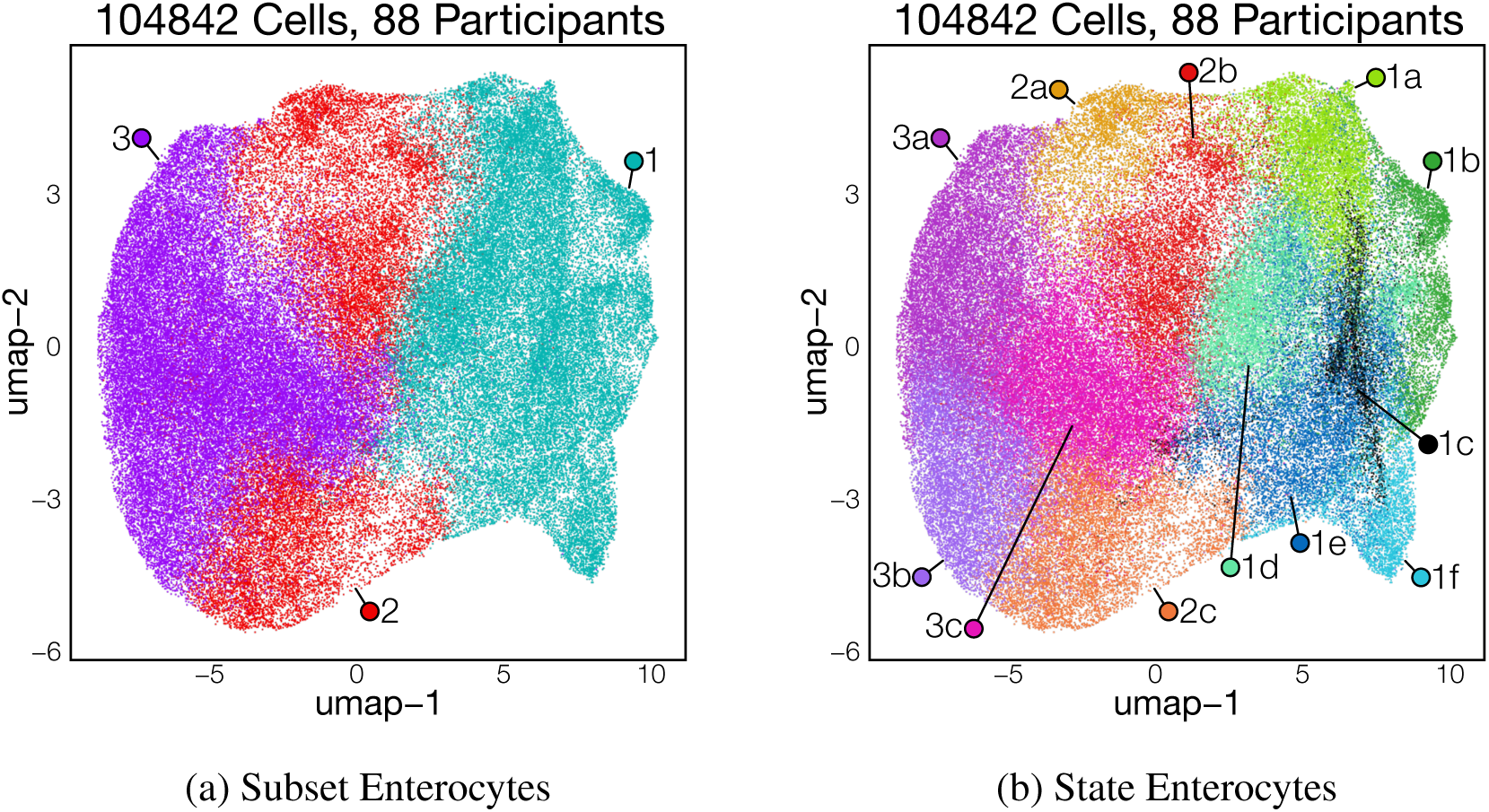
Epithelial Cells

**1. Epithelial progenitor:**

Cells within this cluster do not express major epithelial lineage markers, including *SLC15A1* (Enterocytes), *MUC2* (Goblet cells), *CHGA* (Enteroendocrine cells), *LYZ* (Paneth cells) and *POU2F3* (Tuft cells). Rather, they broadly express crypt markers *OLFM4*, *SLC12A2* and *SOX9*,[57, 58] indicating that they are primarily uncommitted epithelial progenitors. However, only a small subset of cells were positive for the intestinal stem cell marker LGR5.[59] These cells are primarily localized to the villus crypt.[55]

**a. AGR2**^+^ **Secretory progenitor:** These cells express *AGR2*, a gene previously associated with intestinal secretory cells and mucous production.[60, 61]
**b. Transit amplifying cells:** Transit amplifying cells had an active proliferative signature with high expression of *MKI67* and *TOP2A*. Notably, this was the only epithelial cell with a strong proliferation signature in our dataset.
**c. REG3A**^+^ **Progenitor:** These progenitor cells express high levels of the antimicrobial peptide *REG3A* which has been shown to be upregulated following intestinal damage in inflammatory bowel disease and in graft vs host disease.[62, 63] Thus, these cells are likely an injury responsive subset. REG3A^+^ progenitors also express detectable levels of *LGR5* and *ASIC2*, suggesting WNT activation and stem-like properties. Notably, this cell state is significantly enriched in biopsies from Pakistan.
**d. Early enterocyte progenitor:** These cells express crypt-associated markers in conjunction with the upregulation of enterocyte markers. Early enterocyte progenitor cells also express HES1, a Notch-signaling response gene[64] that inhibits *ATOH1* expression and secretory cell formation.[65]
**e. EPHB2**^+^ **Progenitor:** *EPHB2* has been shown to be localized to the crypt, where it functions to help restrict intermingling between crypt and villi localized cells.[66] Therefore, these cells may play a role in villus-crypt demarcation. Cells in this state are defined by expression of *BCL2* and *CDCA7*.
**f. LGR5**^+^ **Stem cell:** These cells are the primary source of *LGR5*, *ASCL2* and *SMOC2*, genes previously associated with the intestinal stem cell (ISC).[59, 58] Approximately 80% of cells in this state express *LGR5*, indicating that most are ISCs. ISCs are a self-renewing population that can give rise to all differentiated epithelial populations of the duodenum[67]

**2. Early Enterocyte:**

This subset is comprised of cells that have recently committed to the enterocyte lineage and are therefore relatively early in terms of enterocyte differentiation. Prior work has suggested that cells with this transcriptional signature tend to be spatially located towards the top of crypts or the bottom of villi. Early enterocytes express the pan-enterocyte marker *SLC15A1*, as well as early enterocyte markers *SLC2A2* and *SLC2A5*.[55] They also express high levels of *CA2* and *MGAM*.

**a. S100G**^+^ **Early enterocyte:** Cells in this state have high expression of *S100G*, a gene believed to be associated with calcium absorption in the proximal intestine.[68]
**b. GSTA2**^+^ **Early enterocyte:** These cells express both *GSTA1* and *GSTA2*, which has previously been shown to be expressed within the duodenum.[69] They also have elevated levels of *KHK*[70] and are the earliest enterocyte population along the pseudotime trajectory that expresses *GLUT2*, possibly indicating a precursor population of specialized carbohydrate absorbing enterocytes.
**c. BTNL8**^+^ **enterocyte:** Cells in this state express both *BTNL3* and *BTNL8*, two receptors necessary for epithelial interaction with *γδ* T cells in the healthy intestine.[71, 72] These cells may therefore represent a resident cell state that mediates interactions with intestinal *γδ* T cells. BTNL8^+^ cells also express higher levels of *TRPM6*, a channel necessary for magnesium absorption.[73]

**3. Late Enterocyte:**

As enterocytes differentiate and progress toward the villus tip, they mature into cell-states specialized for nutrient and electrolyte absorption. This includes cells that exhibit a transcriptional profile associated with carbohydrate absorption and cells defined by transcripts necessary for dietary fat absorption.[55] Late enterocytes express high levels of apoproteins, including *APOC3*, *APOB* and *APOA4*, as well as *CD36*.[74] Consistent with previous work, we also find high expression of purine metabolism genes, *ENPP3* and *SLC28A1*. Interestingly, all late enterocyte states in our dataset express SLC46A1, the folate transporter.[75]

**a. AQP10**^+^ **Late enterocyte:** *AQP10* encodes a pH sensitive aquaglyercoporin channel, that is optimally gated in the range of the physiological pH of the duodenal lumen. Cells in this state are highly specialized for the absorption of the enzymatic products of triglyceride hydrolysis including free glycerol. In addition to *AQP10*, these cells express the aquaglyceroporins *AQP7* and *AQP9*. Among late enterocytes, these cells express elevated levels of APO-family proteins further indicating a highly specialized role in fat absorption.
**b. PTPRR**^+^ **Late enterocyte:** These cells express the tyrosine phosphatase, *PTPRR*, which can suppress WNT signaling.[76] They also express high levels of *IL15* and BTNL8, suggestive of possible role in immune-epithelial cell interaction. [77, 72] They also express the genes required for triglyceride re-esterification indicating a role in lipid absorption.
**c. PEPT1**^+^ **Late enterocyte:** Although SLC15A1 (PEPT1) is broadly expressed in all enterocytes, cells in this state expressed higher levels of *PEPT1* in conjunction with *SLC2A2*, *APOB* and *CD36*. Therefore, these enterocytes are likely represent a multifunctional absorptive population.

### 8.2 Secretory Epithelium

**Figure 8:**
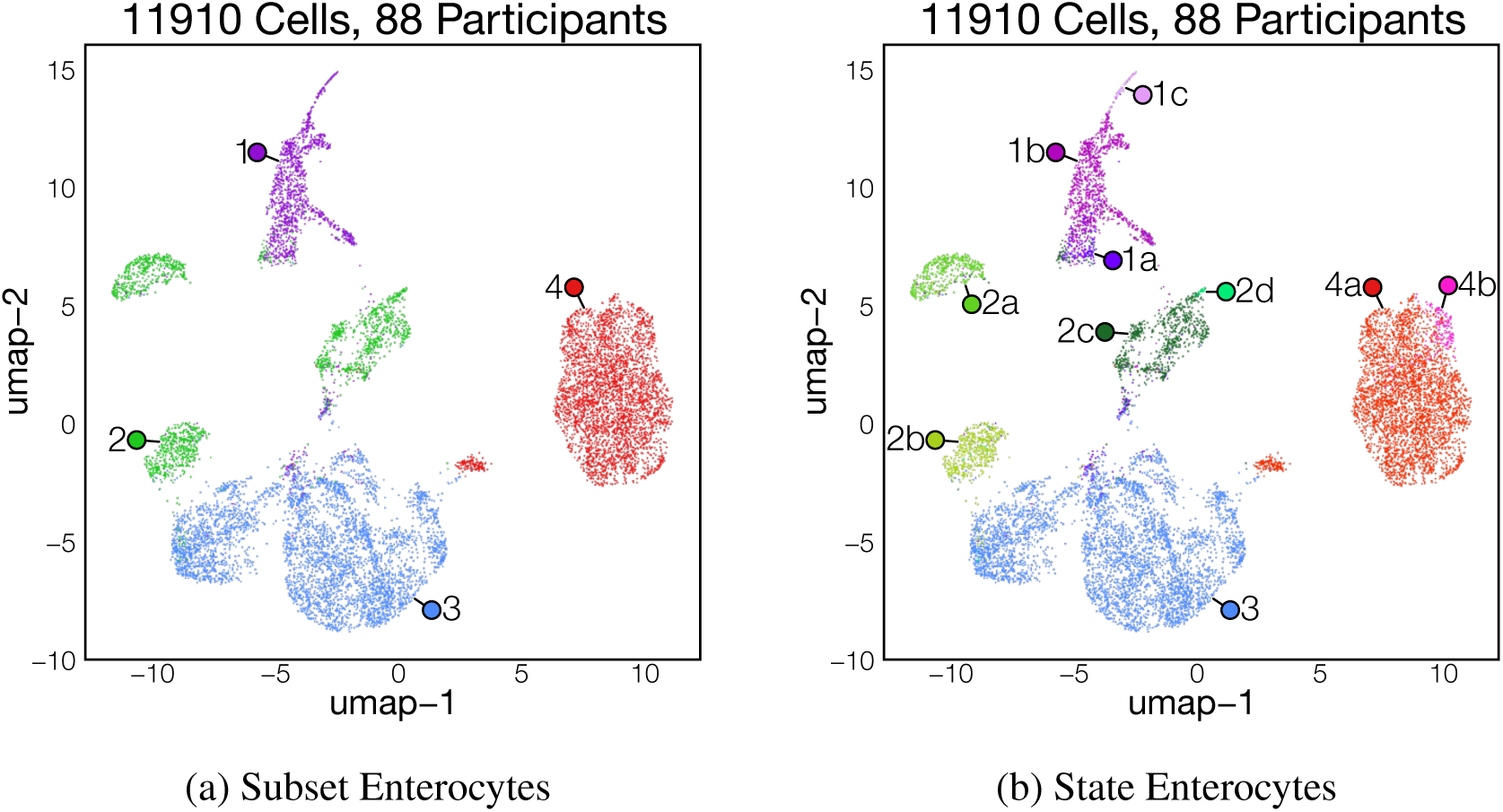
Epithelial Cells

**1. Paneth cells:**

Paneth cells are a subset of epithelial cell that reside within intestinal crypts. These are known to secrete antimicrobial peptides which function maintain a barrier against microbial species.[78] Within the epithelial context, Paneth cells uniquely express high levels of *LYZ*, *DEFA5*, *DEFA6* and other secreted defense factors.[79, 80]

**a. LYZ**^+^ **secretory progenitor:** Cells in this state express *OLFM4* and *LGR5*, as well as Paneth cell markers, including *DEFA6* and *LYZ*, as well as *SOX9*.[81, 82] Therefore, they are likely early Paneth cell progenitors. This state is also enriched for the transcription factors *SOX4*, which has been implicated in ATOH1-independent secretory development and may represent an alternative lineage.[83] Accordingly, these cells do not express *ATOH1*.
**b. Conventional Paneth cell:** These cells make up much of the Paneth cell subset and express markers associated with canonical Paneth cell function.
**c. NKD1**^+^ **Paneth cell:** This state corresponds to a minority of Paneth cells with high expression of the WNT-inhibitor *NKD1*.[84, 85] These cells also have high expression of *CRACR2A*, *CTNNA2*, *HPN* and *PGM5*. The exact role of this cell state is not understood.

**2. Mucous-producing cells:**

This broad subset includes multiple mucin-secreting secretory epithelial cells. As a secretory epithelial cell, mucous-producing cells express secretory-associated genes including *TFF1*, *TFF2*, *TFF3*,[86] *FCGBP*[87], *SPDEF*[88] as well as various types of mucins. Secretions of these cells establish the duodenal mucous layer and provides a protective barrier against microbial threats.[89] They can be further categorized based upon their secretory profile.

**a. Brunner’s gland cells:** Unique to the duodenum, Brunner’s glands are a specialized mucin-producing cell type that primarily produces *MUC6*.[90, 91] We define Brunner’s gland cells as the cells that express *MUC6*, *TFF2*, *PGC* and *AQP5*.[74, 92]
**b. MUC5AC**^+^ **mucous cell:** These cells express high levels of *REG4*, *TFF1* and *MUC5AC*. Both *TFF1* and *MUC5AC* have been attributed to the surface mucous cells of the stomach.[93, 94, 95] In contrast, *REG4* has been described as a marker of deep-crypt secretory cells within the colon.[96] Thus, these are likely a rare duodenal secretory cell with similarity to gastric surface mucous cells.
**c. Goblet cells:** Throughout the GI tract, goblet cells express high levels of the secreted mucin, *MUC2*.[97] In agreement with other scRNA-seq studies, we identify goblet cells based on their high expression of *MUC2*, *TFF3*, *FCGBP* and *CLCA1*.
**d. DUOXA2**^+^ **Goblet cell:** This state is defined by the expression of *DUOXA2*, *DUOX2*, *MUC2* and *IL1B* and represents a rare subset of secretory cells in a small number of participants. Within the intestine, *DUOXA2* expression is associated with dysbiosis and injury[98] and is upregulated in IBD patients.[99] So, this state may signify epithelial damage in the few enrolled participants in whose biopsies these cells were found

**3. BEST4 Enterocytes:**

BEST4 enterocytes are found throughout the intestinal tract and marked by expression of *BEST4*, *CA7* and *SPIB*.[100, 101, 102] Functionally, BEST4 enterocytes have been shown to regulate intestinal fluid efflux using *CFTR* and other ion transporters.[103]

**4. Tuft Cells:**

A specialized epithelial subset that expresses chemosensory receptors, which sense luminal contents and coordinate downstream immune responses through effector cytokines, such as IL25.[104, 105, 106] Tuft cells are described in multiple tissues and share the common markers *POU2F3*, *TRPM5*, *ALOX5*, *GFI1B* and *DCLK1*.[105, 104] Although recent work has identified two major subtypes of tuft cell, in our dataset we identified only one major subset with a minor state.[107] As these differences are associated with position along the crypt-villus axis, it is likely that this cluster of cells consists of a spectrum of Tuft-1/Tuft-2. Furthermore, *IL25* is often ascribed as a marker of tuft cells. However in our cohort, we see very little evidence of tonic IL-25 production.

**a. Conventional Tuft cell:** This subset expresses genes associated with classical tuft cell function as described above and includes most tuft cells in our dataset.
**b. GABA-A**^+^ **Tuft cell:** Cells in this state express high levels of GABA-A receptor subunits, *GABRA4* and *GABRB1*. In mice, GABA activation signature in tuft cells was associated with a high fat diet.[108] However, how GABA may regulate tuft cell function remains an active area of study.

### 8.3 Enteroendocrine cells

Enteroendocrine cells (EECs) are a rare and highly heterogeneous subset of secretory epithelium. They secrete various hormones in response to diet and environmentally derived metabolites. These hormones regulate physiological responses such as gut motility, hunger/satiation and other factors involved in the gut-brain axis. EECs can be classified on the basis of the production of a primary hormone. Broadly, all EECs express *CHGA* and the transcription factor *NEUROD1*.[109, 110]

**Figure 9:**
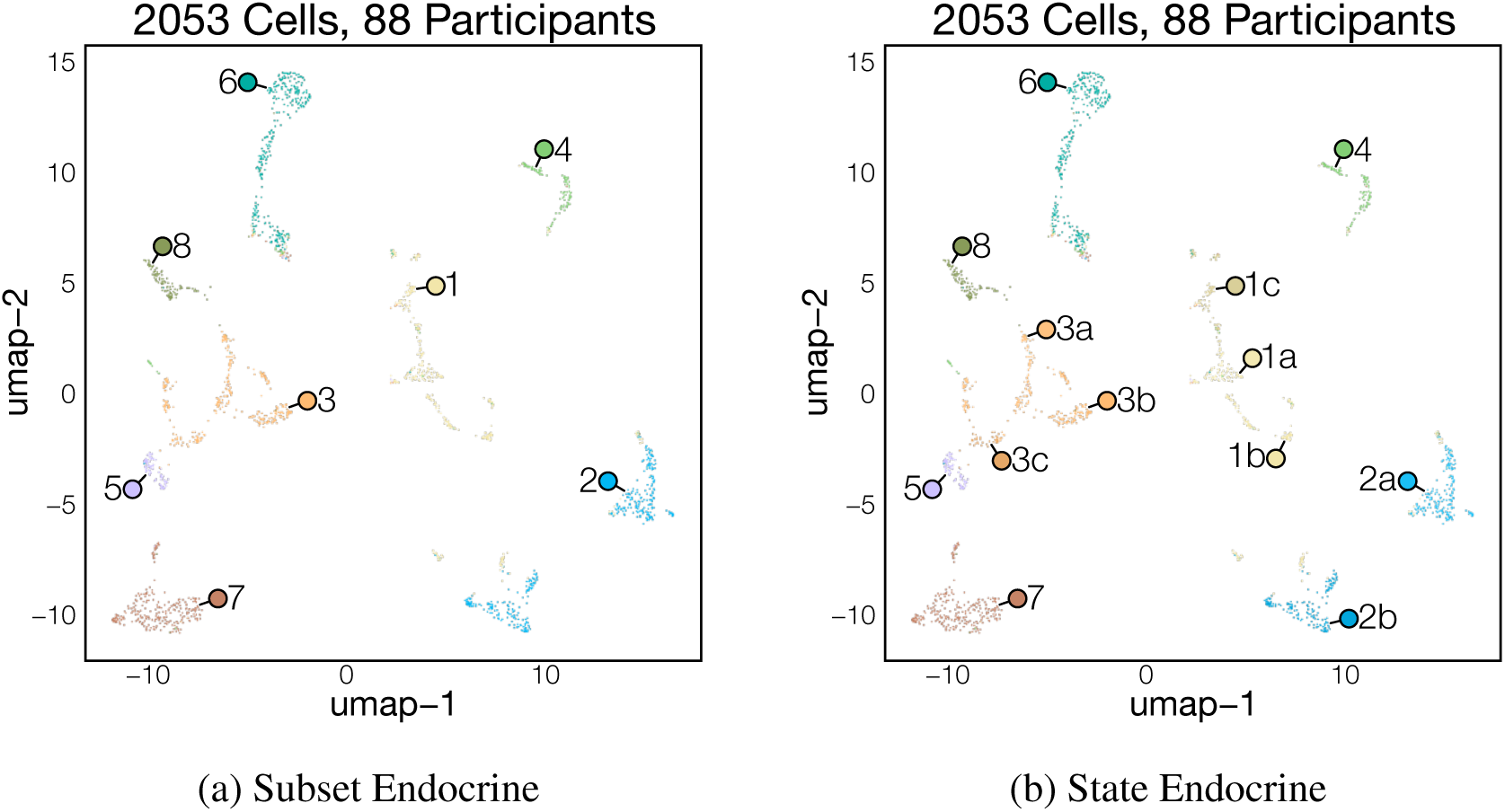
Endocrine Cells

**1. EC progenitors:**

EEC progenitors are defined by expression of *NEUROG3* and other transcription factors involved in cell fate commitment such as *NKX2-2*, *NEUROD1* and *INSM1*. *NEUROG3* is transiently upregulated to specify the endocrine cell lineage and is downregulated upon cell fate commitment.[111, 112, 113] EEC progenitors do not express endocrine hormone genes as they are expressed upon maturation. These cells also have *DLL1* and other genes previously shown to be expressed early in EEC differentiation, including *DLL1*.[67]

**a. NEUROG3**^+^ **progenitor:** These cells express high levels of *NEUROG3* and *NEUROD1* and are likely to be the earliest progenitors that have committed to the enteroendocrine lineage.
**b. PAX4**^+^ **progenitor:** These cells express *PAX4* but not *ARX*, a transcription factor implicated in the development of enterochromaffin cells.[114] Furthermore, a small percentage of PAX4^+^ progenitors express *NEUROG3* and *TPH1*, an enzyme necessary to synthesize serotonin. Therefore, these cells are likely early progenitors of the enterochromaffin lineage.
**c. ARX**^+^ **progenitor:** These cells express high levels of *ARX*, a transcription factor implicated in the specification of peptide hormone producing enteroendocrine subsets.[115] However, they do not express all peptide hormone so they likely represent an early progenitor for many of the peptide hormone producing subsets.

**2. Enterochromaffin cell:**

Enterochromaffin cells synthesize and secrete serotonin and can be identified by the expression of *TPH1*, the enzyme necessary for serotonin synthesis, as well as by elevated expression of *CHGA*.[116] They are localized throughout the gastrointestinal tract[117, 118, 119]

**a. TRPC7**^+^ **Enterochromaffin cell:** These cells uniquely express *TRPC7*; a cation channel activated by diacyl glycerol.[120]
**b. SCT**^+^ **Enterochromaffin cell:** These cells co-express the hormone *SCT* and *TPH1*. In mice, enterochromaffin cells upregulate *SCT* as they move up the villus axis. Therefore, this state likely represents enterochromaffin cells located within the villus.[121]

**3. L/I cell:**

Nearly all participant biopsies contained enteroendocrine cells that co-expresses *CCK* and *PYY*. When expressed alone/in isolation, these genes mark I and L cells respectively. Notably, co-expression of *CCK* with other enteroendocrine hormones has been described.[122, 123] Both *CCK* and *PYY* are anorexigenic and suppress appetite. More specifically, *CCK* functions to promote gallbladder contraction to assist in intestinal digestion.[124]] L/I cells also express high levels of *ONECUT3*, a transcription factor previously associated with I cells.[112, 67] These cells can be further clustered by the expression of a third hormone. Notably, GIP was not expressed by any L/I cell states in our dataset.[122]

**a. GCG**^+^ **L/I cell:** L cells that express *GCG* (which is cleaved into *GLP1*, *GLP2* and other hormones) are much lower in frequency in the duodenum relative to the ileum or colon.[125, 119, 117] Nevertheless, we did identify cells that co-express *GCG* with *PYY* and *CCK* in some participant biopsies.
**b. SCT**^+^ **L/I cell:** Similar to enterochromaffin cells, L/I cells can express *SCT* and this has also been observed to correlate with position along the villus. Therefore, these cells are likely positioned near the villus tip relative to other L/I cells.[121] Interestingly, cells in this state also express high levels of *IL33* and *GHRH*.
**c. GAST**^+^ **L/I cell:** In the stomach, *GAST* expression marks G cells and regulates the secretion of gastric acid.[126] G cells in the duodenum have been prescribed previously[127, 119], however GAST^+^ L/I cells in our dataset also co-express *CCK* and *PYY*. They also express *SPX*, *SYT1* and *ADGRL3*.

**4. D cell:**

D cells are enteroendocrine cells that express the peptide hormone *SST* which inhibits many aspects of metabolism, including the release of hormones from other enteroendocrine cells and neurons.[128]D cells have been previously defined by their exclusive expression of the transcription factor *HHEX*.[116] Within our dataset, *GABRA1*, *GABRB2* and *GABRG2* are also expressed in D cells.

**5. K cell:**

K cells express *GIP* and are enriched in the proximal small intestine.[119] GIP enhances glucose-dependent insulin release and promotes the storage of fat in adipocytes.[124] In addition to *GIP*, K cells can be identified by expression of the bile acid sensor *BASIC* (ASIC5)

**6. M/X cell:**

Some enteroendocrine cells co-express *MLN* (M cells) and *GHRL* (X cells). This has been observed previously and these were designated as M/X cells[116, 129] Ghrelin release promotes the sensation of hunger in anticipation of meals.[124] Function of motilin is less characterized due to the absence of a functional gene in rodents.[130, 131] Like ghrelin, it is released during fasting and has been implicated in inducing hunger sensation.[131] Motilin induces smooth muscle contraction and this has been suggested to increase gastric emptying and overall gastrointestinal motility.[132] M/X cells express *ACSL1*, *IL20RA* and *BHMT* in addition to *MLN* and *GHHL*.

**7. SCGB2A1^+^ cell:**

Cells in this cluster have not been described previously. They express high levels of *SCGB2A1*, which encodes for mammaglobin, a secretoglobin found within the mammary gland. The exact function of mammoglobin is not well understood. SCGB2A1^+^ cells also express high levels of the succinate receptor (*SUCNR1*), which has been shown to modulate Tuft cell secretory activity.[133]

**8. TRH^+^ cell:**

Cells in this state express *NEUROD1* and the hypothalamic hormone *TRH*. They also uniquely express *KLHL4*, *SYT10*, *CDH10* and *NCAM2* along with low levels of hormones previously associated with enteroendocrine function. Within the central nervous system, TRH functions to indirectly induce the secretion of thyroid hormone and regulate systemic metabolism.[134] The exact role of these previously undescribed enteroendocrine cells is currently unknown.

## 9 T/NK/ILC

Cells which express high levels of *PTPRC* clustered together. These consist of T cells, NK cells and ILCs, with a majority being T cells. Therefore, the top differentially expressed genes consists of T cell markers, including *SKAP1*, *CD247* and *CD3E*. By further clustering, we separate the different lymphocytes into their respective cell subtypes.

Quality control metrics (200 *<* nFeature_RNA *<* 4000 and percent.mt *<* 25%)

### 9.1 CD4 T cells

**Figure 10:**
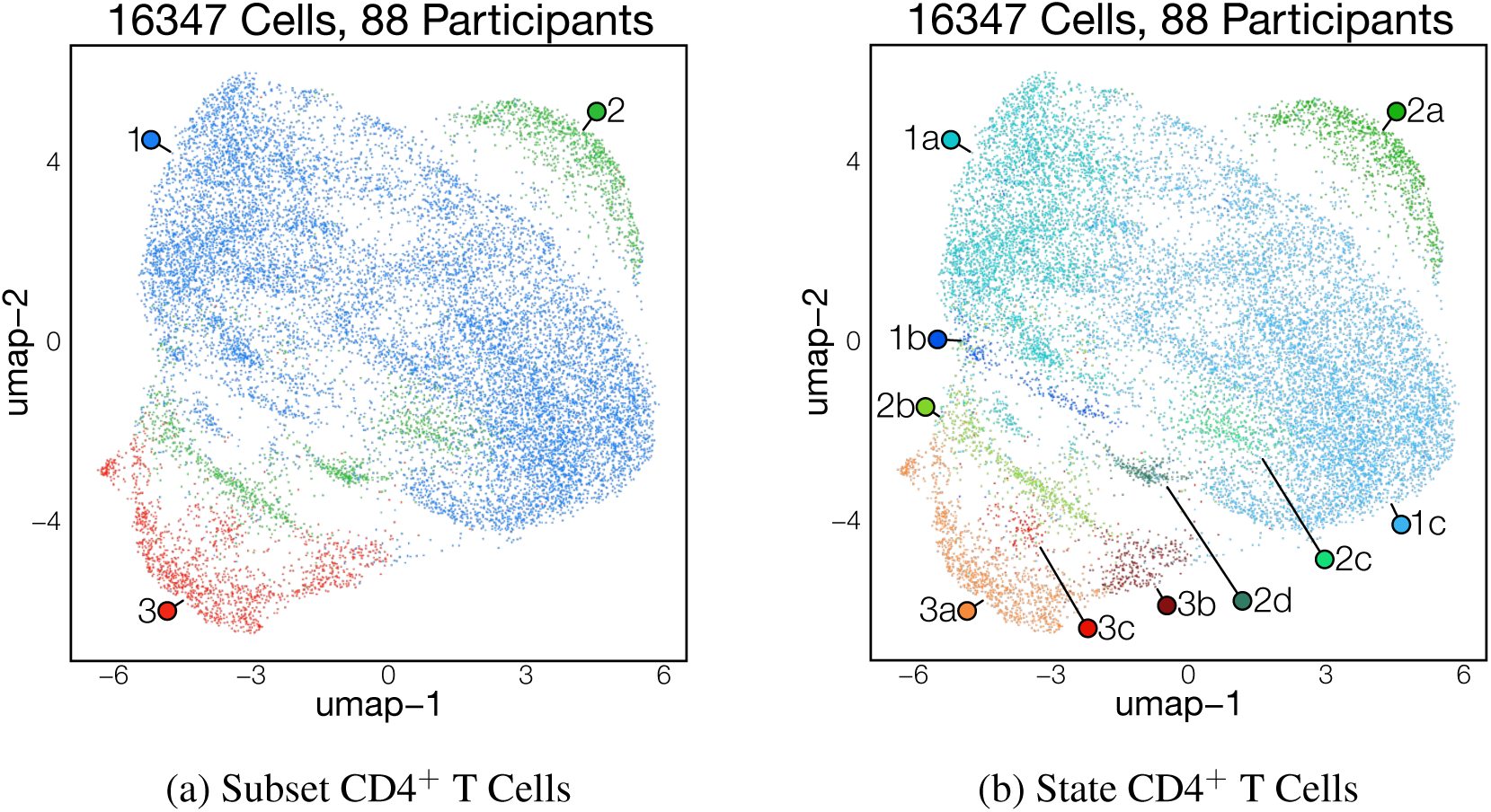
CD4^+^ T Cells

**1. Resting CD4 T:**

These cells express both *CD4* and *TRAC*. In contrast to activated T cell subsets they do not express cytokines previously associated with CD4 T cell effector function nor do they express *CCR7* or *SELL*, which are used to define naïve and central memory T cell subsets. Resting CD4 T cells express high levels of transcription factors and chemokine receptors, which may indicate they were previously activated. Notably, cells within this subtype express *ICOS*, *CD40LG* and *CCR6*. In other cell atlases, resting Th17 cells were defined by *CcR6* expression.[135]

**a. RUNX1**^+^ **resting CD4 T:** These CD4 T cells have high expression of *RUNX1* and *RUNX2*. In mouse models, *RUNX1* has been shown to be necessary for maturation of peripheral CD4 T cells.[136] RUNX1^+^ resting CD4 T cells also express *GRIP1*, *PTPN13* and *ADAM12*. These may be resting Th1 cells as *ADAM12* has been implicated as a costimulatory molecule in Th1 formation.[137]
**b. DLGAP1**^+^ **resting CD4 T:** *DLGAP1* has been implicated in post-synaptic densities of neurons. Its exact role in T cells is unknown.[138]
**c. CXCR4**^+^ **resting CD4 T:** These are the majority of resting CD4 T cells. The ligand for *CXCR4* is *CXCL12*, which is highly expressed in stromal cells across many tissues, including the intestine.[139] Cells in this state are also enriched for *CD40LG*, *GPR183* and *IL4I1*. *IL4I* is a marker of human Th17 T cells, so these are likely resting Th17 cells[140]

**2. Activated CD4 T:**

This subtype includes antigen experienced CD4 cells that can be binned into conventional helper T cell subsets. These subsets were primarily defined by active cytokine expression.

**a. Th17:** In scRNA-seq, Th17 cells express high levels of *IL17A*, *IL23R* and *IL22* which are cytokines associated with type-17 inflammation.[141] Notably, tonic IL22 functions to maintain barrier function of the intestinal epithelium. Global depletion leads to increased bacterial translocation in both health and disease.[142, 143] These cells also express *CCR6*, *IL4I* and *RORC*.
**b. pTfh:** Peripheral Tfh-like (pTfh) cells express *PDCD1*, *IL21* and *CXCR5*, which are markers of Tfh cells.[144] Tfh cells are primarily found within secondary lymphoid tissue, including specialized gut-associated lymphoid tissue, where they support B cell activation and maturation.[144, 145] However, the duodenum has a significantly lower density of both Peyer’s patches and isolated lymphoid follicles relative to the rest of the intestine, with these structures being almost completely absent from the proximal duodenum.[146, 147] While pTfh cells have been described in inflammatory diseases[148, 149], their presence in the healthy duodenum warrants further investigation.
**c. Th1:** Within our dataset, Th1 cells are defined as CD4 T cells that express *IFNG* and *IL12RB2*. Th1 also express chemokine receptors *CXCR6* and *CCR2* and have elevated levels of *CD40LG*.
**d. IFIT3**^+^ **activated CD4 T:** This state was found in a small number of study participants. These cells are marked by interferon-response genes and may indicate an active response to a viral infection.

**3. Treg:**

Tregs express the master transcription factor *FOXP3* and are known to have an immunoregulatory role mediated by *CTLA4*, *IL10* and *IL2RA*. We defined Tregs as CD4 T cells that express *FOXP3*. In our dataset, all Treg states expressed elevated levels of *CTLA4* and *IL2RA*. Notably, we do not observe cluster of *RORC* cells expressing Tregs previously described in other segments of the intestine.[150, 151] In our dataset, *RORC* transcripts were not well detected.

**a. IKZF2**^+^ **Treg:** IKZF2 (Helios) encodes a transcription factor which functionally stabilizes Tregs.[152] Within the intestine, Helios has been shown to mark a specific population of thymic-derived Tregs.[153, 154] We defined IKZF2^+^ Tregs as CD4 T cells that express *IKZF2* and *FOXP3*.
**b. PDCD1**^+^ **Treg:** A small subset of Tregs that did not express *IKZF2* had elevated levels of *PDCD1*. These cells are rich in ribosomal genes, which may indicate a more quiescent state. They also express *CCR7* and *SELL*, which may indicate a resting-like state.
**c. IL10**^+^ **Treg:** These Tregs express high levels of *IL10*, a cytokine released upon activation of the cell to suppress inflammation particularly in the intestine.[155, 156, 157] *IL10* expression may indicate a more effector-like phenotype. However, a recent study in mice demonstrated that a subset of colonic Tregs constitutively express IL10.[158]

### 9.2 CD8 T cells

**Figure 11:**
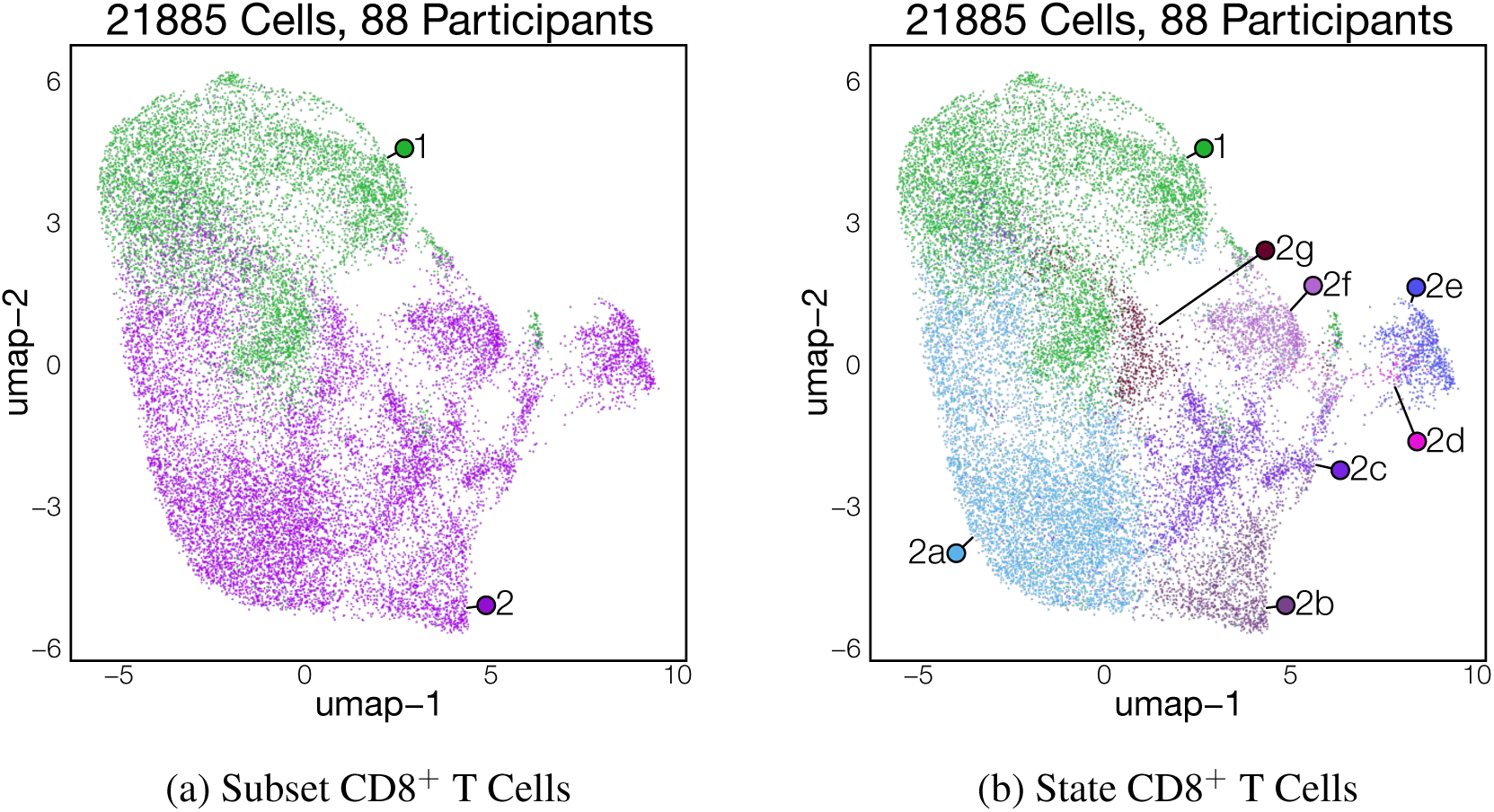
CD8^+^ Cells

**1. Resting CD8 T**

CD8 T cells were broadly defined as *αβ* T cells (positive for both *TRAC* and *TRBC1*, negative for *TRDC* and *TRGC2*), that express *CD8A* and *CD8B*. Resting CD8 T cells had a strong ribosomal gene signature which has been implicated in poised-like states.[159, 160]Despite the absence of effector gene expression, these cells express both *CD69* and *ITGAE*, indicating a Trm state.[161]

**2. Activated CD8 T:**

Activated or effector CD8 cells express inflammatory cytokines, chemokines and cytotoxic granule contents including *IFNG*, *GZMA*, *GZMK* and *CCL4*. They can be further classified by the other effector molecules that they express.

**a. Effector CD8 T:** CD8 T cells with low granzyme expression in the presence of *IFNG* and *KLRK1*, suggesting an activated phenotype.
**b. CCL4L2**^+^ **activated CD8 T:** CD8 T cells in this state have high expression of *IFNG*, *CCL3*, *CCL4*, *CCL3L2*, *CCL4L2* and *CCL5*, but express low levels of *GZMA* and *GZMB*.
**c. CCL3**^+^ **activated CD8 T:** Similar to other activated CD8 T cell states, these cells express *CCL3* in the absence of *CCL3L2* and *CCL4L2*. Cells in this state express *GZMA* and *MHC class I* genes.
**d. CX3CR1**^+^ **Cytotoxic CD8 T:** *CX3CR1* has previously been used as a marker of effector T cell differentiation, with CX3CR1^+^ effector T cells giving rise to both peripheral memory and effector memory CD8 T cells[162, 163] Accordingly, this state has a peripheral memory phenotype in which cells express *SELL*. Consistent with prior studies, these cells are highly cytotoxic with expression of *GNLY*, *GZMB* and *PRF1*.
**e. GNLY**^+^ **Cytotoxic CD8 T:** Although all cytotoxic CD8 T cells express *GZMB*, *GZMA* and *PRF1*, only CXCR1^+^ Cytotoxic CD8s and cells within this subset express GNLY. GNLY^+^ cytotoxic CD8 T also express *ZNF683* (Hobit). Hobit is present in long-lived effector CD8 T cells and regulates tissue residency in conjunction with *PRDM1* (Blimp1).[164, 165] This likely reflects a highly cytotoxic state of tissue resident CD8 T cells.
**f. GZMK**^+^ **activated CD8 T:** GZMK^+^ T cells have been described extensively in the context of inflammatory disease such as rheumatoid arthritis and IBD.[166] Unlike *GZMA* and *GZMB*, *GZMK* has little cytotoxic potential, and instead activates the complement pathway.[167] Here we find that GZMK^+^ T cells are present at steady state in the duodenum.
**g. Cytotoxic CD8 T:** Cytotoxic CD8 T cells express high levels of *GZMA* and *GZMB* in the absence of *GNLY*. This state is likely closely related to the GNLY^+^ state and may represent a less mature cell state.

### 9.3 *γδ* T cells

*γδ* T cells were defined by co-expression of *TRDC* with either *TRGC2* or *TRGC1*. It was important to consider all TCR constant regions, as ILCs and NK cells were found to have high expression of the delta chain but not the gamma chain, and CD8 T cells had high expression of the gamma chain but not the delta chain.[168]

**Figure 12:**
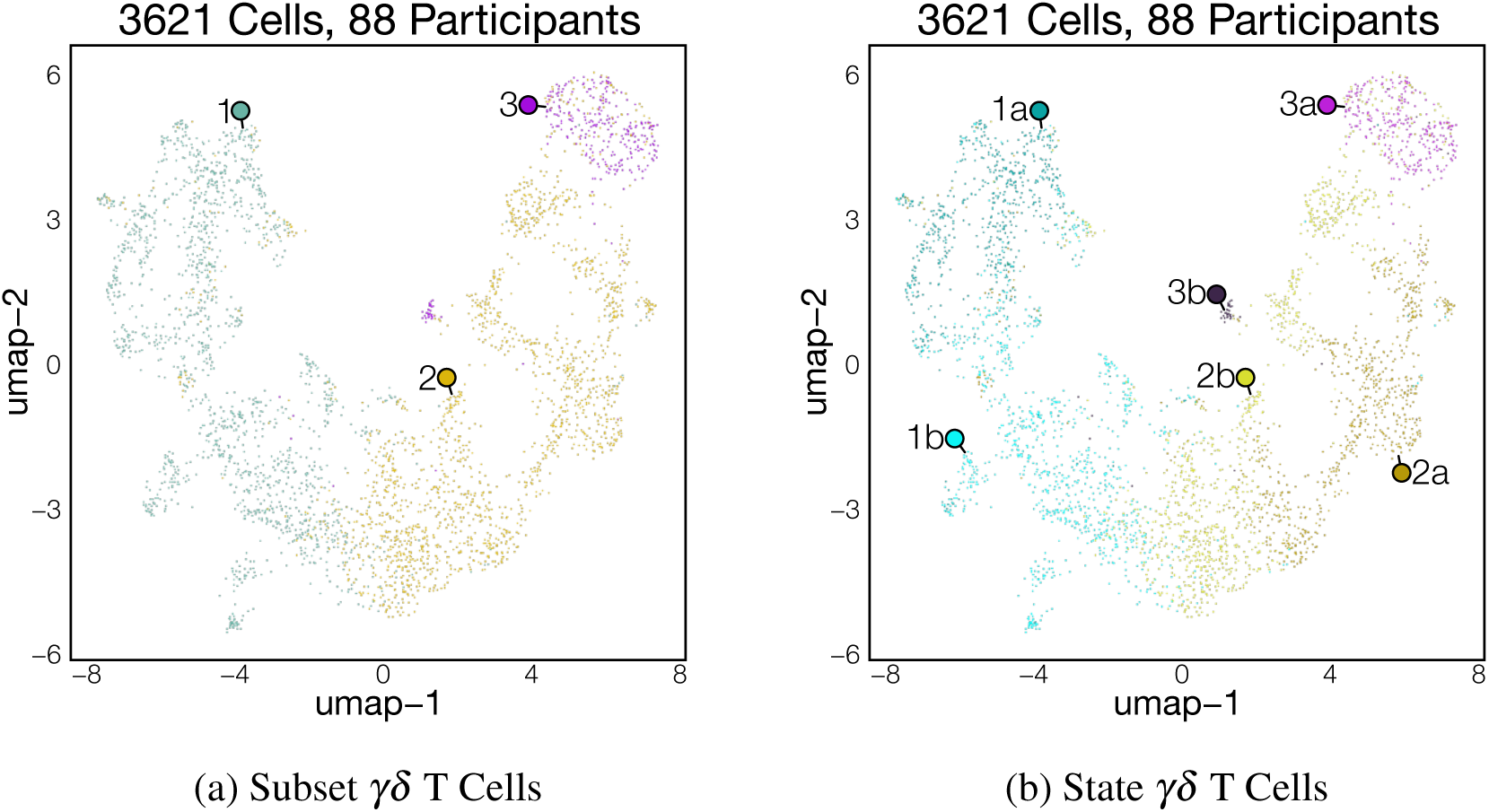
*γδ* Cells

**1. Naïve *γδ* T:**

Naïve *γδ* T cells have previously been defined by the expression of *TCF7* and *LEF1* in the absence of effector genes.[169]

**a. IKZF2**^+^ **naïve** *γδ*: IKZF2 (Helios) is highly expressed in this states. Helios expression has been described in a subset of circulating *γδ* T cells, where it may mark recent activation.[170] Cells in this state also express *RTKN2* and *TIAM1*.
**b. CD63**^+^ **naïve** *γδ*: Cells in this state express *CD63* and Hobit (*ZNF683*). *CD63* has been shown to be upregulated on T cells following antigen stimulation, where it can act as a costimulatory molecule.[171] Considering this as well as the role of Hobit in maintaining tissue residency, these cells may be a recently activated naïve population.[172]

**2. Resting *γδ* T:**

Similar to resting CD8 T cells, resting *γδ* T cells express cytotoxic granule genes at low levels. They express high levels of the chemokine ligands *XCL1* and *XCL2*. *XCL1* has been shown to be necessary to position cDC1 populations in the murine intestine and may be important for tolerance induction.[173, 174]

**a. AREG**^+^ *γδ* **T:** *AREG* encodes a cytokine implicated in epithelial regeneration and disease tolerance.[175, 176] Therefore, this *γδ* T cell state may be a tissue resident population critical for the maintenance of intestinal tolerance. Cells in this state also express the innate receptor genes *KLRC3* and *FCER1G*.
**b. Resting** *γδ* **T:** Cells in this state express high levels of *XCL1* and *XCL2* in the absence of *AREG*. They are the majority of the resting *γδ* T cells in our dataset.

**3. Activated *γδ* T:**

Relative to resting cells, activated *γδ* T cells express high levels of *GZMA* and *GZMB* as well as the cytotoxic receptor *NKG7*. This cytotoxic gene expression profile may reflect recent receptor activation.

**a. Cytotoxic***γδ* **T:** The majority of cells within this subset are highly cytotoxic and express innate killing receptors, such as *FASLG*, *NKG7*, *KLRD1*, *KLRC3* and *KLRC2*.
**b. IFIT3**^+^ *γδ* **T:** This rare state of activated *γδ* T cell was found in a small number of study participants. These cells are marked by interferon-response genes and may indicate an active response to a viral infection.

### 9.4 ILCs/NKs

**Figure 13:**
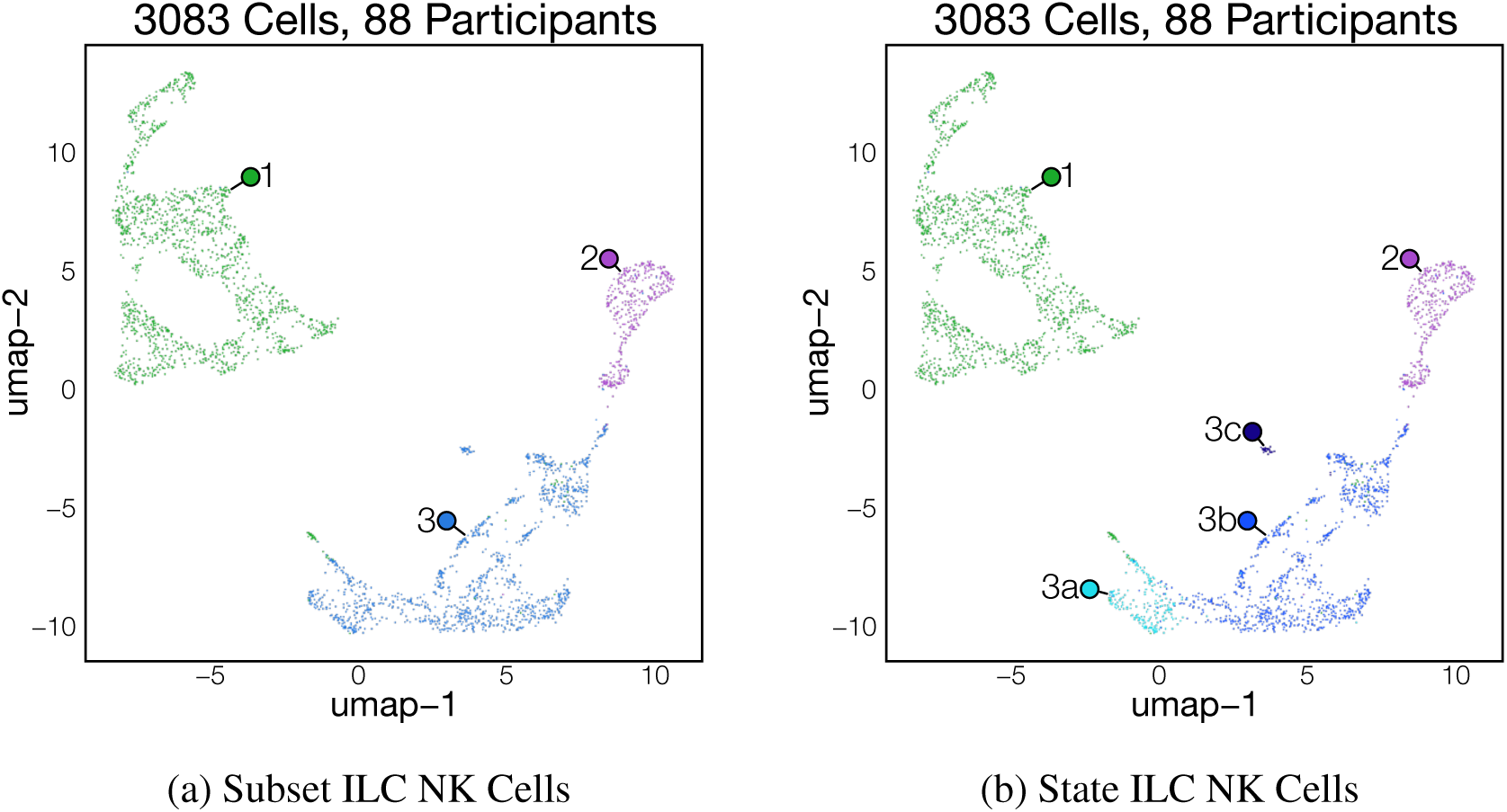
ILC NK Cells

**1. ILC3:**

ILCs initially clustered with T lymphocytes even though they do not express *TRAC* and *TRGC2*. ILCs have high expression of *KIT*. The majority of cells in this subset also express *IL23R*, *RORC* and *CCR6*, indicating that they are ILC3s.

**2. Circulating NK:**

This subset represents NK cells that have recently entered the intestine and/or NK cells that were within the microvasculature at the time of sample collection. Circulating NK cells express *FCGR3A*, *CX3CR1*, and have significantly elevated levels of *GZMB*, *PRF1* and *GNLY*.

**3. Resident NK:**

NK cells are a cytotoxic population of innate lymphocyte that express *NCAM1* and cytotoxic granule contents such ae *GZMA* and *GNLY* in the absence of *TRAC* and *TRGC2*. Resident NK cells also express *FCER1G* and *TYROBP*. In contrast to the circulating population, resident NK cells do not express *FCGR3A*.

**a. NCR2**^+^ **NK:**: NCR2 or NKp44 positive NK cells have previously been described in the healthy adult intestine, where they were the majority of NK cells.[177, 178] These cells express lower levels of *NCAM1* and may correspond to the CD56^dull^ population.[179] NCR2^+^ NK cells were primarily found to be localized within the intestinal epithelium.[178]
**b. Mucosal NK:** These cells express high levels of *NCAM1* as well as numerous chemokines, including *XCL1*, *CCL3* and *CCL4*. Mucosal NK cells express higher levels of the transcription factor *EOMES* than other NK cells. Prior description of an EOMES^high^ intestinal NK cell population used *KLRC1*, which is expressed in this state in our dataset. Prior work has described, a mucosal NK cell population expressing similar markers, to decrease in frequency during infancy, an observation that we corroborate in our dataset.[178]
**c. IFIT3**^+^ **NK:** Biopsies from a small number of participants had a strong interferon signature in multiple cell populations, including NK cells. This may indicate that these individuals had an active infection at the time of biopsy collection.

### 9.5 Other lymphocytes

**Figure 14:**
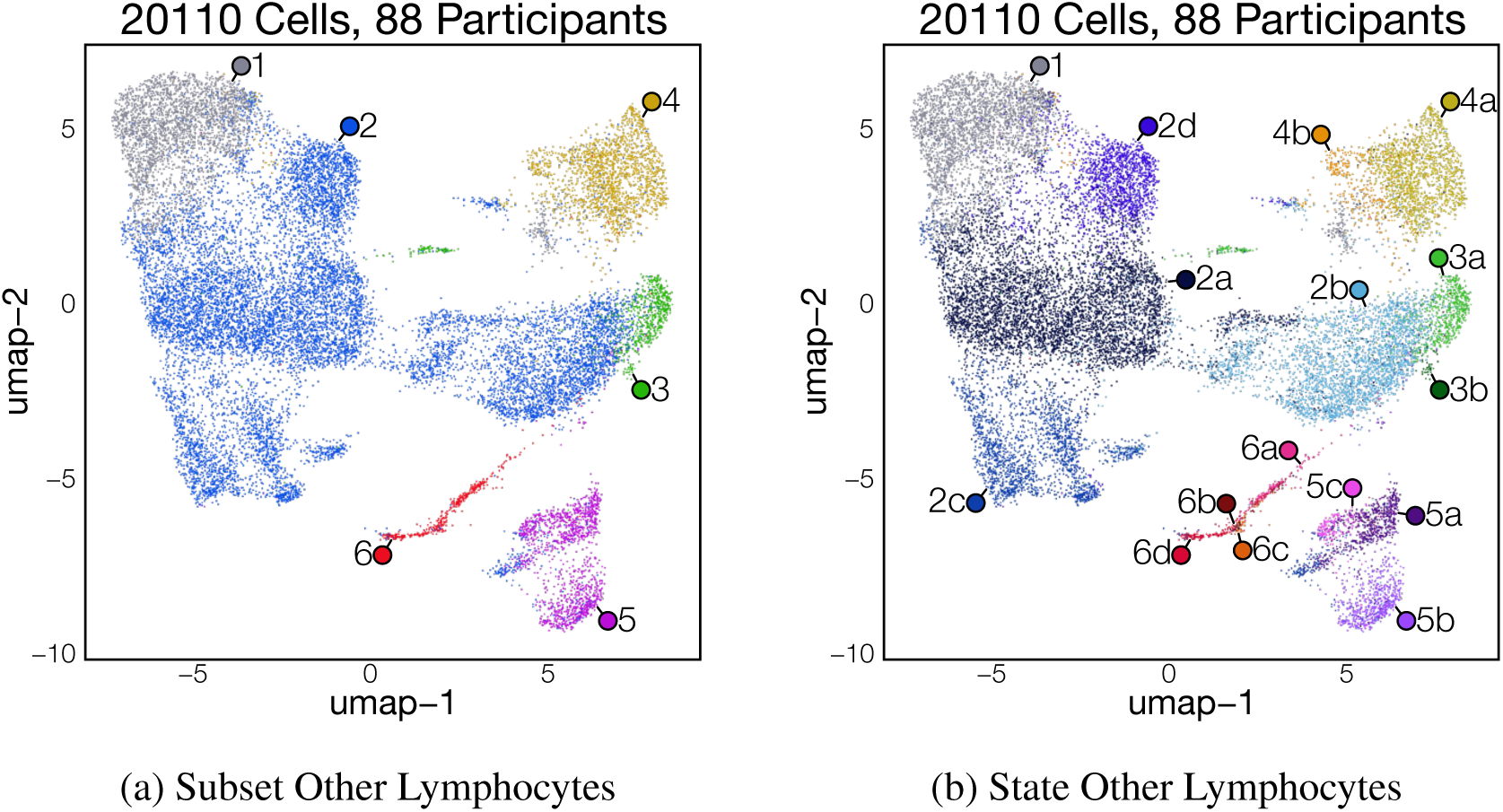
Other Lymphocytes

**1. FOSB^+^ Lymphocyte:**

This subset may represent low quality cells as it is difficult to distinguish what type of lymphocyte these cells may correspond to. Furthermore, these cells have high expression of various lncRNAs, *FOS*, *FOSB* and *JUN*, which indicates they may be an activated cell.

**2. IELs:**

Intraepithelial lymphocytes (IELs) are a subset of T cells that exist within the epithelial compartment of the intestine and other mucosal sites. IELs were defined by the high expression of ITGAE, an adhesion molecule necessary to enter the epithelial layer.[180] Prior work investigating IELs using flow cytometry or scRNA-seq has demonstrated that they are het-erogenous populations composed of both *αβ* and *γδ* T cells. Furthermore, they consist of thymic derived or natural populations as well as peripherally induced populations.

**a. Resting IEL:** Similar to T cells within the lamina propria, Resting IELs were defined based on an elevated presence of ribosomal genes. They also have elevated *ITGAE* expression relative to other resting lymphocyte states.
**a. CD8***αβ* ^+^ **IEL:** IELs that express *CD8A* and *CD8B* are a subset of induced or conventional IEL, that develops from classical CD8 T cells upon encountering a foreign antigen.[180] Cells in this state are defined by high expression of *CD8A*, *CD8B* and *ITGAE*.
**a. ZBTB16**^+^ **innate IEL:** Unconventional or natural IELs develop directly from thymic T cells upon encountering self-antigen.[180] As a result, they have innate-like properties and express the innate T cell master transcription factor *ZBTB16* or *PLZF*. Unlike ILC1s, these cells express *TRAC*, but are negative for *CD4* and *CD8*, indicating that they may be double negative populations.
**a. FOSB**^+^ **IEL:** Cells in this state are similar to FOSB^+^ lymphocytes and are likely a recently activated population.

**3. Naïve *αβ* T:**

Naïve *αβ* T cells express high levels of *SELL* and *CCR7*, two factors necessary for migration to lymph nodes. Similar to naïve *γδ* T cells, they expressed high levels of *LEF1* and *TCF7*. They do not express the residency marker *CD69*.

**a. Naïve** *αβ* **T:** The majority of Naïve *αβ* T cells have no discernible state and are high for ribosomal genes as previously reported. This state includes both CD8 and CD4 T cells, as both co-receptors are expressed. [5, 181] Although naïve T cells have been shown to primarily occupy secondary lymphoid tissue, previous publications have demonstrated that naïve T cells seed the tissue at various points during development and inflammation.
**b. IFIT3**^+^ **naïve T:** *IFIT3* is an interferon response gene and defines an interferon responsive naïve T cells. These cells were present in only a small subset of individuals.

**4. MAIT cell:**

Mucosal-associated invariant T (MAIT) cells are a subset of T cells prevalent at mucosal sites, such as the gastrointestinal tract. MAIT cells express a limited diversity of TCR alpha chains that allow them to detect antigen in the context of the MHC I-like molecule *MR1*.[182] MAIT cells are innate-like T cells and can be activated through TCR-dependent and TCR-independent mechanisms. They initiate a rapid response upon activation.[183] MAIT cells have been defined in scRNA-seq by expression of *TRAC*, *KLRB1* and *SLC4A1*, which have been previously shown to be highly expressed in cells that recognize MR-1. These cells also express *ZBTB16*.[184]

**a. Resting MAIT:** Similar to CD4 and CD8 T cells, resting MAITs do not express cytokine genes. Instead, they express cytokine receptors, chemokine receptors and various transcription factors. Notably, these cells also express high levels of *CCR6*, *IL7R* and *IL23R*, indicating that they may be poised toward Il17 production.
**b. Activated MAIT:** In contrast, activated MAIT cells express various cytokines through which they mediate effector functions. Activated MAIT cells express *IL17A* and *TNF*, in addition to classical MAIT markers.

**5. Innate T:**

Innate T cells are a less understood subset of T cell that we defined by expression of *ZBTB16*, *FCER1G*, and *TYROBP*. This subset has been described in the context of cancer, where it was demonstrated to have an important cytotoxic role.[185] Although they express high levels of *TRDC* and *TRAC*, Innate T cells do not express either *TRGC1* or *TRGC2*, indicating that they are likely *αβ* T cells.

**a. Resting Innate T:** Cells in this state express high levels of *FCER1G*, *TYROBP*, *KRT81* and *KRT86*. However, they express very little cytokines or other effector molecules.
**a. Cytotoxic Innate T:** Upon activation, these cells have been shown to have high cytotoxic potential.[185] Innate T cells in this state express high levels of *GZMA* and *GZMB* in the absence of *GNLY*.
**b. AREG**^+^ **Innate T:** This activated innate T cell state expresses *AREG* and has a cytotoxic profile. *AREG* has been shown to have an important role in wound repair and tissue tolerance.[186]

**6. Proliferating T:**

In addition to resting and activated states, we also observed a distinct subset of cells that have high expression of *MKI67*, *TOP2A*, *STMN1* and many other genes involved in DNA replication and mitosis. These proliferating T cells tend to cluster together due to this strong transcriptional signature; however, the identities can be further defined by performing sub-clustering on the proliferating cells and looking for genes used to define previous cell type annotation. In our dataset we identified proliferating cells from multiple different subtypes of T cells.

**a. Proliferating CD8 T**
**b. Proliferating CD4 T**

c. Proliferating *γδ* T

d. **Proliferating IEL**

