## Supplementary Methods and Figures for "Development of the Early Childhood Duodenum across Ancestry, Geography and Environment"

### Supplementary Data 1

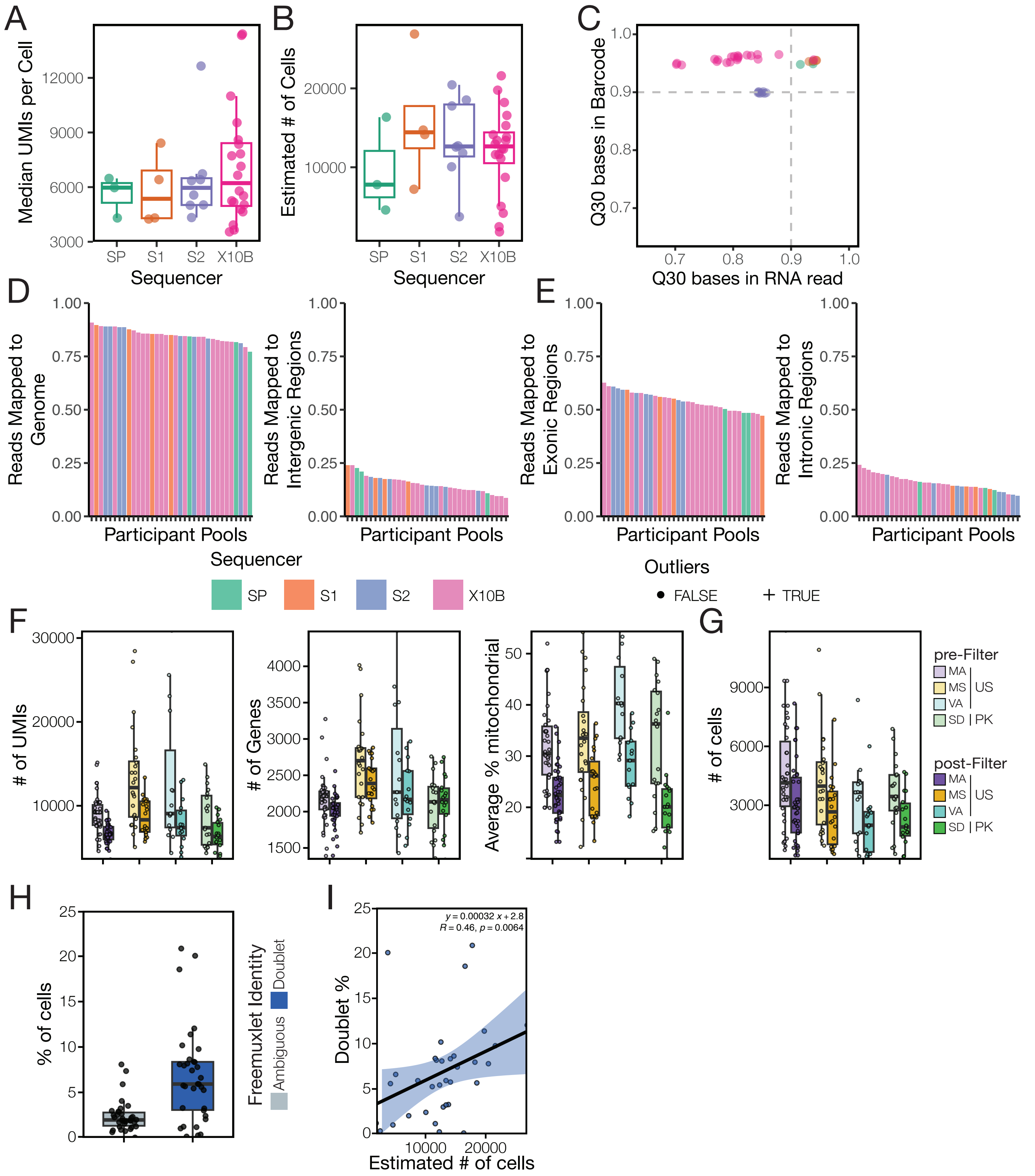


**Supplementary Data 1:** ***Quality control metrics of Gut-AGE scRNA-seq.*** **A and B**. Median unique molecular identifiers (UMIs) detected per cell **(A)** and Estimated number of cells per pool **(B)**. Each point represents a participant pool and color corresponds to the sequencing platform. **C.** The proportion of bases with a Q30 score (sequencing error rate < 1/1000) in the sequenced RNA read versus the sequenced cell barcode. Each point represents a participant pool and color corresponds to the sequencing platform. **D.** The proportion of total reads mapped to the genome (left) versus intergenic regions (right). **E.** The proportion of gene-aligned reads that align to exons (left) or introns (right) before and after filtering of low-quality cells. **F-G.** The average UMIs, unique genes and mitochondrial percentage identified per cell for each participant **(F)** and the total number of cells identified per participant **(G)** before and after filtering of low-quality cells. Participants were colored by sample site and filtration status. H. The proportion of cells per pool that were either unable to be attributed to a participant (ambiguous, grey) or that contained identifying features of 2 or more participants (doublet, blue). I. Estimated number of cells per participant pool versus the proportion of cells called a doublet by Freemuxlet. Correlation coefficient and statistical significance was calculated using a Spearman's rank correlation test.

### Supplementary Data 2

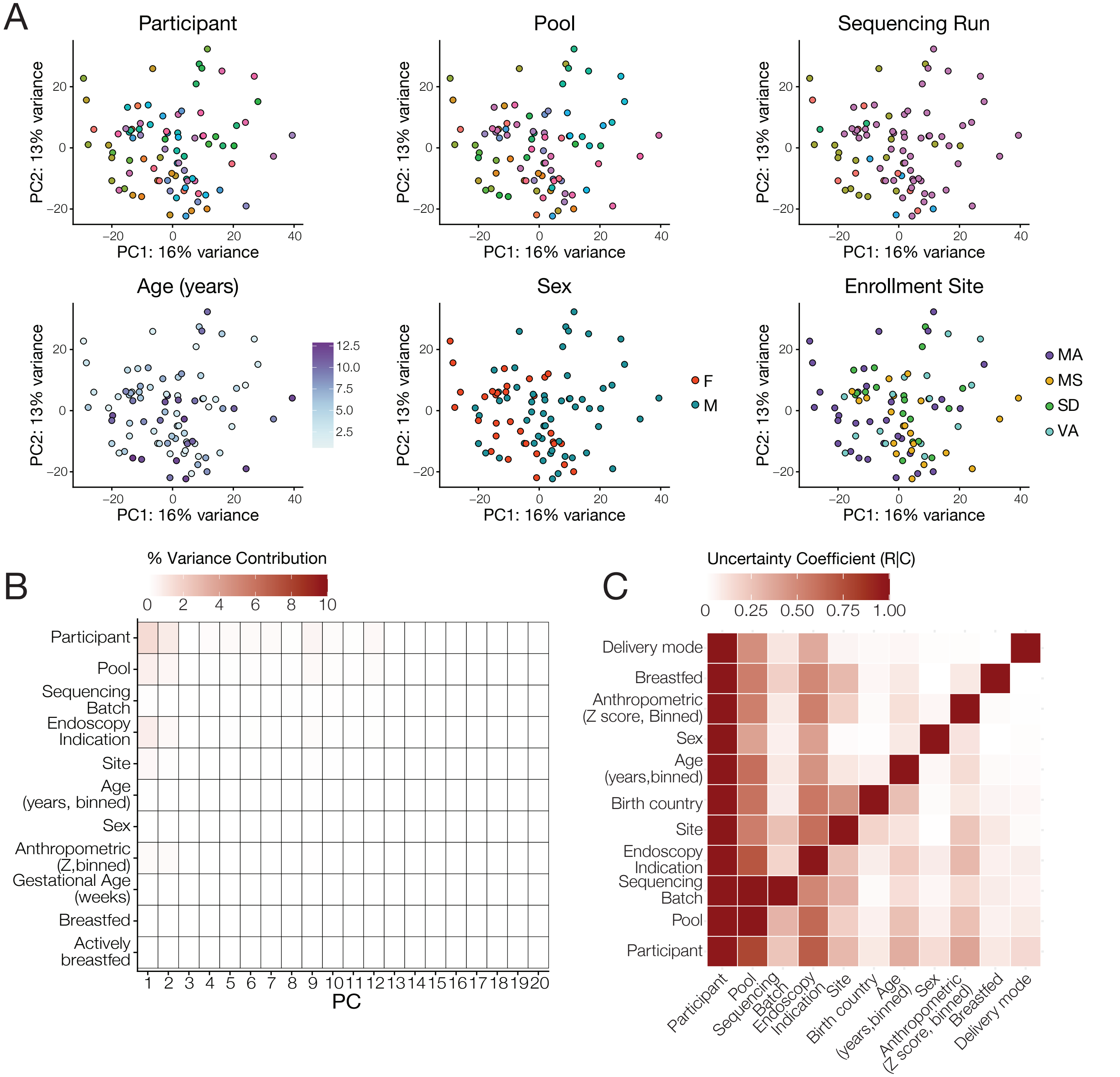


**Supplementary Data 2:** ***Effect of technical variables on scRNA-seq data variance.*** **A.** Principal component plots generated from pseudobulked participant transcriptomic data colored by different variables in the dataset. Top: Colored by technical variables, participant ID (top left), pool (top middle) and sequencing run (top right). Bottom: Colored by biological variables age (bottom left), sex (bottom middle) and site (bottom right). **B.** Heatmap measuring percentage contribution of major technical and biological covariates to the total variance captured by the principal components (PCs) from scRNA-seq. **C.** Heatmap of the proportion of row variable (R) information explained by the column variable (C) as measured by the uncertainty coefficient (Theil's U) for major categorical covariates.

### Supplementary Data 3

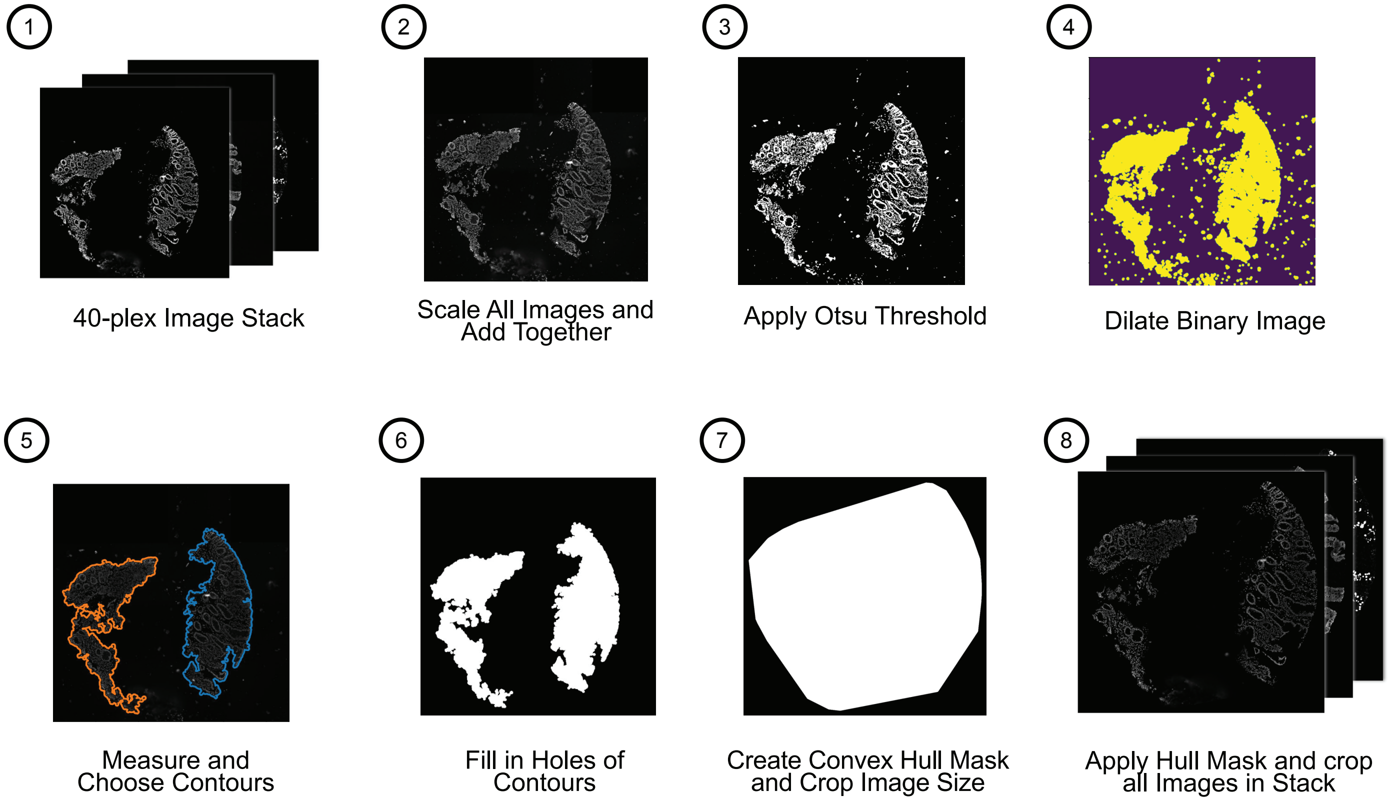


**Supplementary Data 3:** ***Generating the tissue mask.*** Sequence of steps involved in image pre-processing performed prior to cell segmentation to reduce non-tissue signal and remove excess pixels from the images.

### Supplementary Data 4

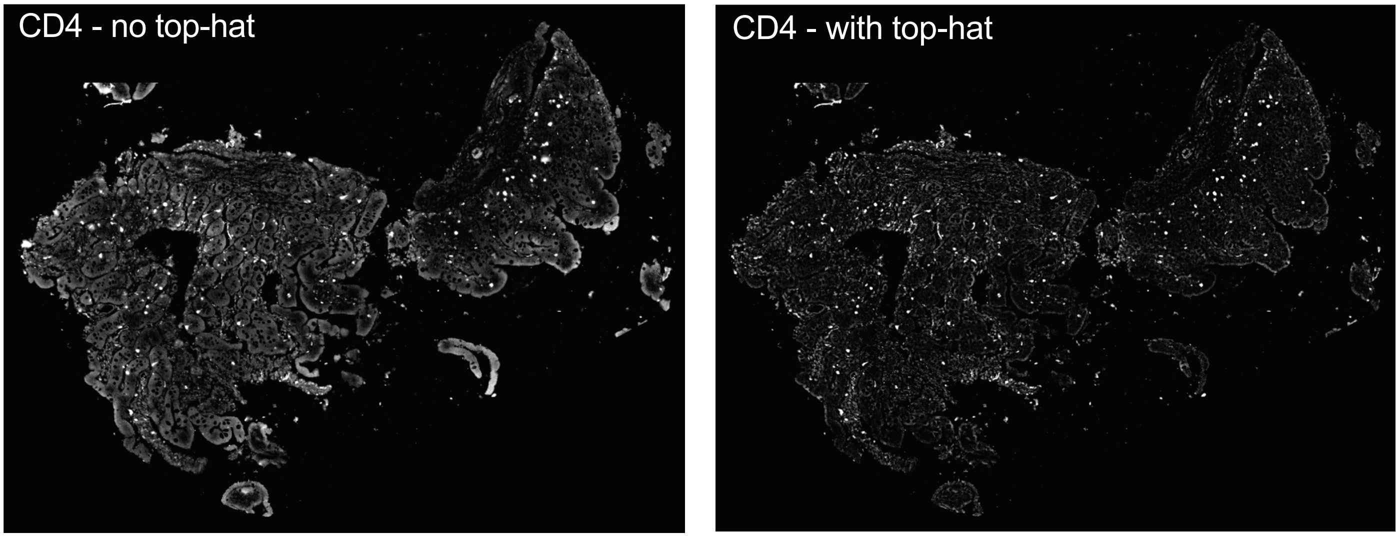


**Supplementary Data 4:** ***Effect of top-hat filter.*** Representative images of a biopsy stained with CD4 (grey) without (left) and with (right) a top-hat filter.

### Supplementary Data 5

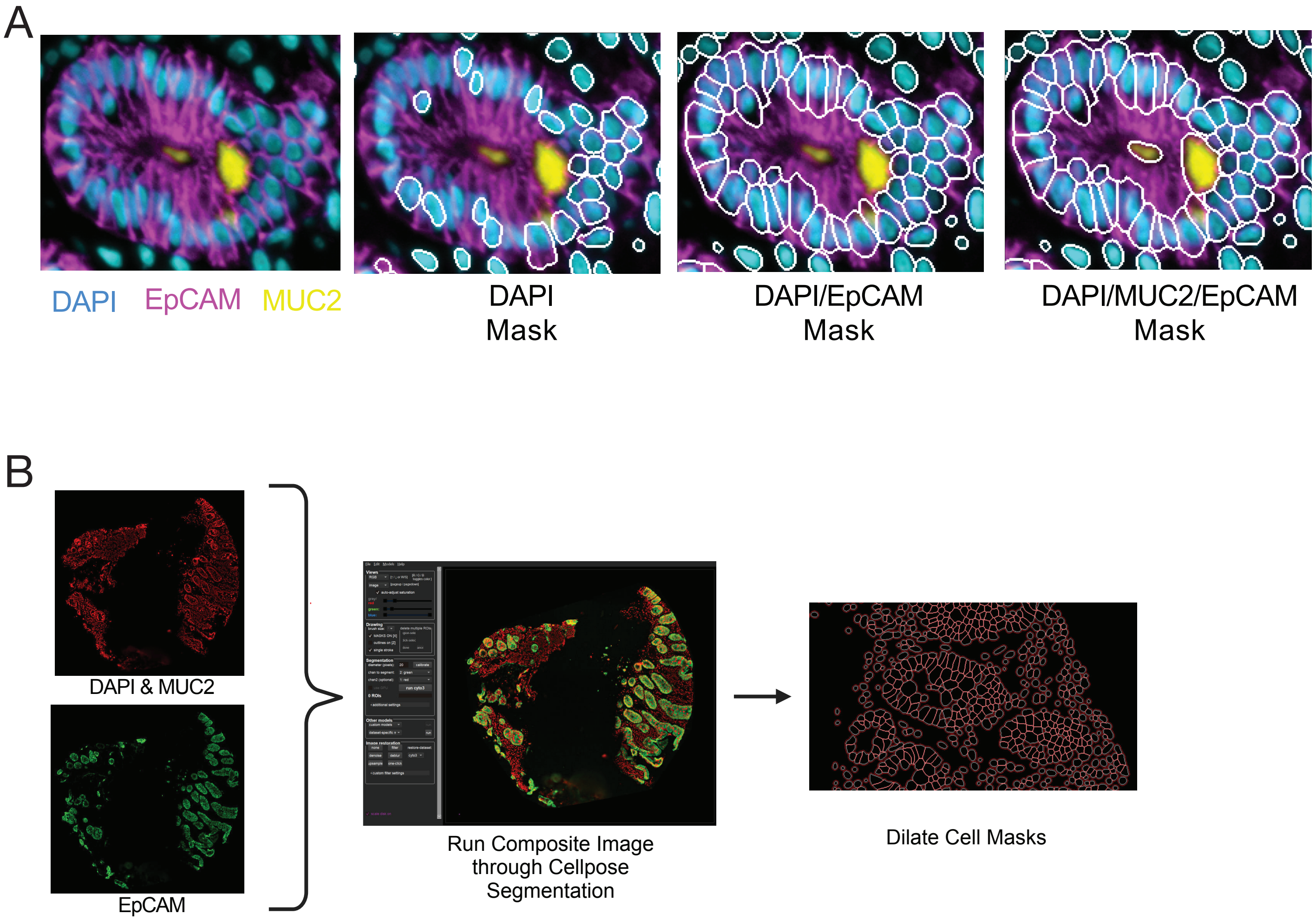


**Supplementary Data 5:** ***Cell segmentation using Cellpose.*** **A.** Representative images demonstrating how the choice of input stain(s) for cell mask generation in Cellpose impacts segmentation. The first panel shows tissue stained with DAPI (blue), EPCAM (pink) and MUC2(yellow). In the subsequent images, the segmentation mask (white) is overlaid when DAPI (left), DAPI and EPCAM (middle), or DAPI, EPCAM and MUC2 (right) are used as the input stains. **B.** Schematic overview of cell segmentation process. DAPI/MUC2 and EPCAM stained images are input to Cellpose which uses the composite image for segmentation, then the resulting cell masks are dilated for subsequent analysis.

### Supplementary Data 6

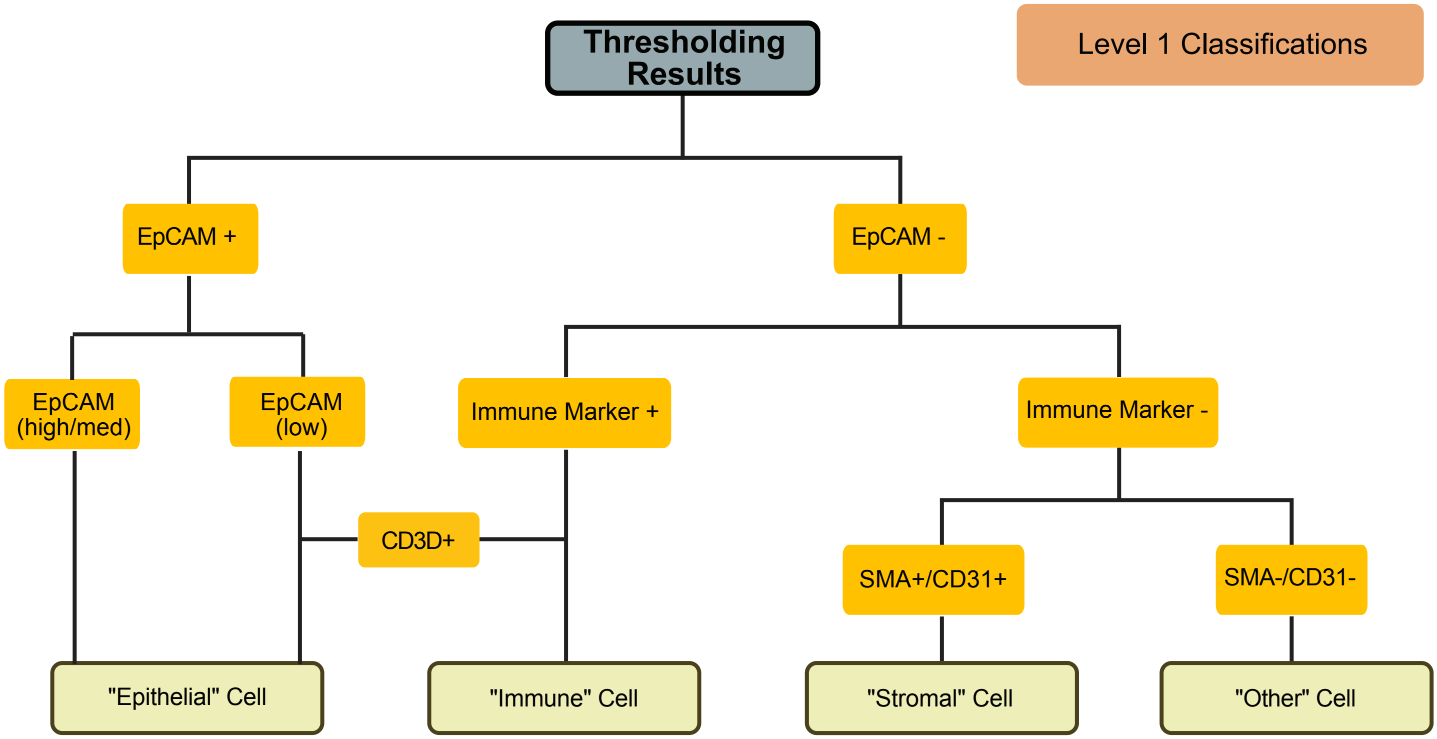


**Supplementary Data 6:** ***Level 1 Cell Type Classification.*** Decision tree for level 1 cell type classifications, which groups each cell into one of 4 major categories (epithelial, immune, stromal, others).

### Supplementary Data 7

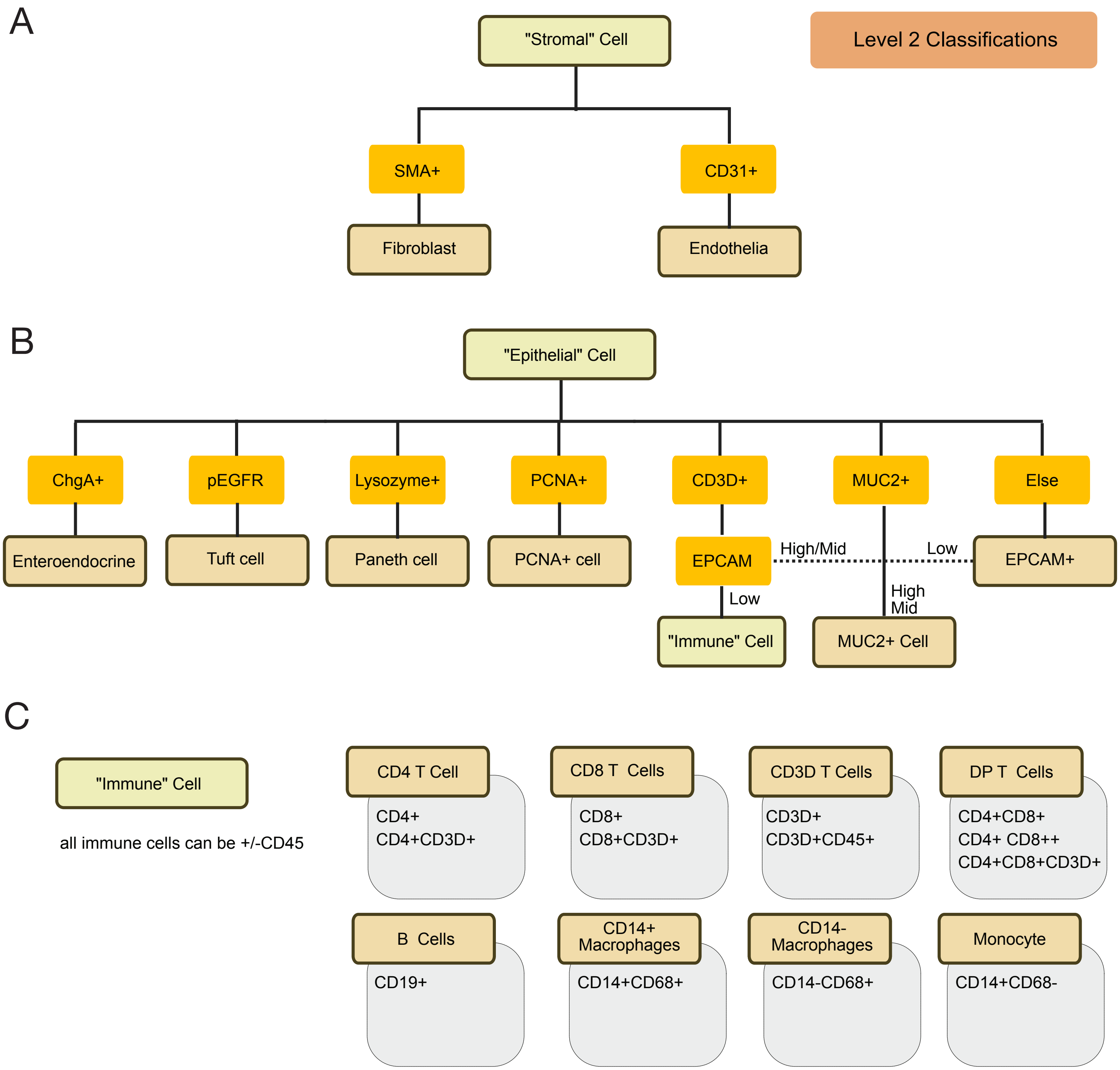


**Supplementary Data 7:** ***Level 2 Cell Type Classification.*** These decision trees are used to classify Level 2 cell types within the three major categories established by the preceding Level 1 classification. Three decision trees were used in level 2 classification, one each for **(A)** stromal cells **(B)** epithelial cells and **(C)** immune cells.
